# Personalized reference genome-based pipeline reveals comprehensive haplotype-resolved views of cancer genomes

**DOI:** 10.64898/2026.05.28.728591

**Authors:** Yoshitaka Sakamoto, Yotaro Ochi, Yasunori Kogure, Shota Kato, Aiko Sato-Otsubo, Masahiro Sugawa, Yosuke Tanaka, Taro Tsujimura, Takashi Mikami, Genta Nagae, Kenichi Chiba, Ai Okada, Yu Ito, Hajime Suzuki, Takuya Yamamoto, Hiroyuki Aburatani, Yuhki Koga, Itaru Kato, Junko Takita, Hiroyuki Mano, Seishi Ogawa, Keisuke Kataoka, Motohiro Kato, Yuichi Shiraishi

**Author notes:** Corresponding author: Yuichi Shiraishi, Ph.D., 5-1-1, Tsukiji, Chuo-ku, Tokyo, 104-0045, Japan.

## Abstract

Cancer genome analysis relies on standard human reference genomes, but detecting somatic alterations in highly repetitive or individual-specific regions remains challenging. We developed the Personalized Reference genome-based Cancer Genome Analysis Pipeline (PRCGAP, https://github.com/yos-sk/PRCGAP), to our knowledge, the first comprehensive pipeline integrating haplotype-resolved analyses of somatic point mutations, structural variants, copy number, and DNA methylation on personalized diploid reference genomes, with each variant phased and annotated by genomic features. We applied PRCGAP to eight tumor-normal cell line pairs and three newly collected pediatric B-cell acute lymphoblastic leukemia (B-ALL) clinical samples. PRCGAP recovered most standard reference-based variants (86.7–96.4% across variant types), validated by orthogonal short-read sequencing, while additionally detecting variants that standard references missed (23.4% of SNVs and 32.6% of SVs). Notably, PRCGAP uncovered variants in centromeric and telomeric regions, and somatic single-nucleotide variants were enriched in centromere dip regions, the putative kinetochore formation sites. Using PRCGAP outputs, we identified L1 retrotransposition source sites absent from standard references and showed that a B-ALL IGH::*DUX4* fusion arose from a *DUX4* pseudogene within an internal D4Z4 repeat unit, rather than the canonical full-length *DUX4* gene. Within a single haplotype-resolved framework, PRCGAP opens regions of the cancer genome that are inaccessible to standard references.

## Introduction

Somatic variants, including single nucleotide variants (SNVs), short insertions/deletions (indels), and structural variants (SVs), drive carcinogenesis and cancer progression^1–3^. A common approach for detecting these variants relies on mapping sequencing data to standard human reference genomes, such as GRCh38 or T2T-CHM13^4^. However, highly repetitive regions such as centromeres, segmental duplications, and telomeric sequences, as well as regions harboring ethnic or individual variation, are commonly referred to as genomic “dark matter”, in which accurate variant detection is challenging.

Recent advances in long-read sequencing technologies, including PacBio HiFi and Oxford Nanopore Technologies ultra-long reads (ONT-UL), have enabled the construction of the first complete haploid human reference, T2T-CHM13^5^. Combined with phasing information from trio, Hi-C, or Pore-C data, these technologies now allow the scalable construction of nearly complete diploid human genomes using de novo assembly tools such as hifiasm and Verkko^6–8^. These advances continue to drive the construction of pangenome references representing diverse populations^9–12^.

These long-read assembly approaches have recently been applied to cancer genome analysis. Personalized reference genomes constructed from matched normal samples by de novo assembly have improved somatic variant calling accuracy and uncovered variants residing in sequences absent from standard references^13–15^. Diploid de novo assembly-based approaches have been shown to substantially reduce false-positive somatic SV calls^16–18^. However, existing approaches have either performed integrated analysis of a single sample without providing a pipeline, or provided pipelines targeting only one class of somatic alteration, such as SVs. A comprehensive pipeline that brings the major classes of somatic alterations, including haplotype-resolved point mutations, SVs, copy-number alterations, and DNA methylation, together on the same personalized diploid reference genome has not yet been established. Moreover, none of these approaches has been systematically applied to clinical cancer samples.

Here, we present the Personalized Reference genome-based Cancer Genome Analysis Pipeline (PRCGAP), to our knowledge, the first comprehensive cancer genome analysis pipeline based on personalized diploid reference genomes (Fig. 1). Using a personalized diploid reference genome, PRCGAP maps tumor and normal long-read data in a haplotype-aware manner using haplotype-specific markers (21-mers) and performs downstream analyses including somatic point mutation and SV calling, haplotype-resolved copy-number profiling, and DNA methylation analysis. For each detected variant, PRCGAP automatically assigns the haplotype of origin, annotates it by overlap with genomic features, and lifts the coordinates over to standard references (GRCh38 and T2T-CHM13).

**Figure 1.**
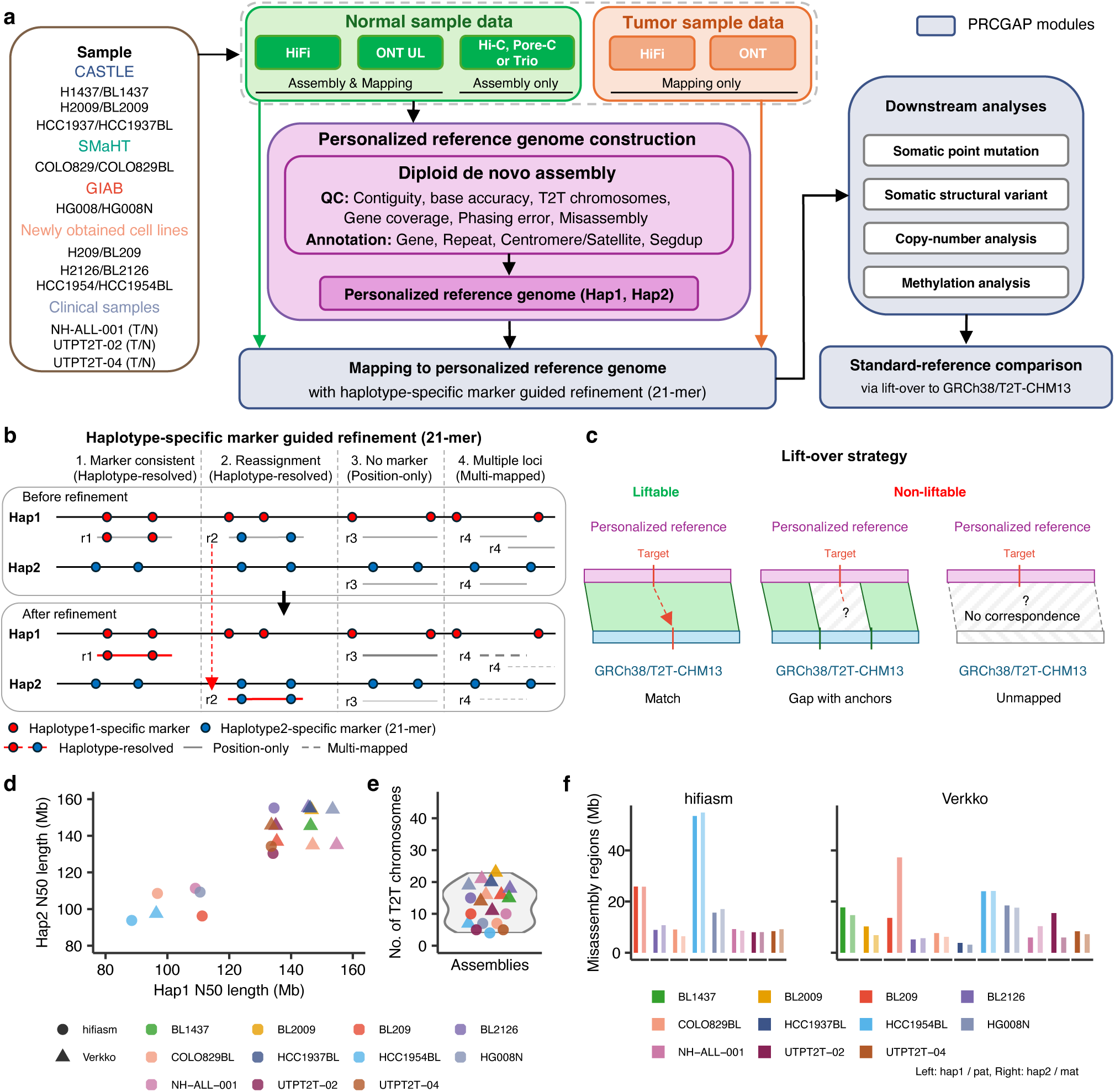
Overview of the Personalized Reference genome-based Cancer Genome Analysis Pipeline (PRCGAP) and evaluation of the constructed personalized reference genomes. (a) Schematic overview of PRCGAP and this study. The personalized reference genome is quality-checked and annotated with gene models, repeat elements (RepeatMasker), centromere/satellite, and segmental duplications (Segdup). See Methods for details. (b) Haplotype-specific marker guided refinement (21-mer). Four representative cases (1–4; reads r1–r4) are shown before (top) and after (bottom) refinement on the haplotype 1 (Hap1) and haplotype 2 (Hap2) references: (1) Marker consistent, where markers match the alignment (Haplotype-resolved, red); (2) Reassignment, where markers indicate the opposite haplotype and the alignment is reassigned (red dashed arrow; Haplotype-resolved); (3) No marker, where reads align to both haplotypes without haplotype-specific markers to determine the haplotype of origin (Position-only, gray solid; see Supplementary Fig. 3a for detailed classification criteria); and (4) Multiple loci (Multi-mapped, gray dashed). After refinement, the selected alignment is shown as a thicker line and unselected alignments as thinner, lighter ghost lines. (c) Lift-over strategy that enables comparison of PRCGAP results with standard-reference-based variant calling results and identification of variants inaccessible to such analyses. Coordinates are liftable when they align to GRCh38/T2T-CHM13-based coordinates (Match) and non-liftable when they do not. Non-liftable coordinates have two states: “Gap with anchors”, where flanking coordinates are liftable, and “Unmapped”, where they do not align to any GRCh38/T2T-CHM13-based coordinates. (d) Contig N50 lengths of haplotype 1 against haplotype 2 for each assembly. Point shapes indicate the assembler (circle, hifiasm; triangle, Verkko) and colors indicate the sample. (e) Distribution of the number of telomere-to-telomere (T2T) chromosomes per assembly. Individual points represent assemblies, with shape and color denoting the assembler and sample as in d. (f) Total length (Mb) of misassembly regions per haplotype, estimated by integrating results from Flagger, Inspector, and NucFlag for hifiasm (left) and Verkko (right). For each sample, the left bar corresponds to haplotype 1 (hap1)/paternal (pat) and the right bar to haplotype 2 (hap2)/maternal (mat).

In this study, we constructed personalized reference genomes by de novo assembly from matched normal samples and rigorously evaluated and annotated every assembly for genes, repeats, centromeres, and segmental duplications. We then applied PRCGAP to eight matched tumor-normal cell line pairs, comprising five publicly available datasets and three newly generated in this study^14,19–21^. In addition, we applied PRCGAP to newly generated data from three pediatric B-cell acute lymphoblastic leukemia (B-ALL) clinical samples with distinct molecular features. We demonstrated that PRCGAP identified somatic variants in centromeric and telomeric regions that are challenging to analyze by standard reference-based approaches. Building on these PRCGAP outputs and the personalized reference genomes, we identified somatic L1 retrotransposition source sites absent from GRCh38 and T2T-CHM13 and elucidated the genomic structure of the IGH::*DUX4* fusion gene, an oncogenic event in B-ALL, demonstrating that the fusion arose from a *DUX4* pseudogene within an internal D4Z4 repeat unit of the macrosatellite array, rather than the canonical full-length *DUX4* gene.

## Results

### Personalized reference genome-based cancer genome analysis pipeline (PRCGAP)

To perform cancer genome analyses based on a personalized reference genome, we developed a new pipeline called Personalized Reference genome-based Cancer Genome Analysis Pipeline (PRCGAP, Methods, Fig. 1a). PRCGAP uses a personalized diploid reference genome assembled from matched normal samples with hifiasm^7^ or Verkko^8^ using long-read sequencing data, including HiFi and ONT-UL data together with Hi-C, Pore-C, or trio phasing data (Fig. 1a). Long-read sequencing data from both tumor and normal samples are then mapped to this reference in a haplotype-aware manner using haplotype-specific markers (21-mers, Fig. 1b). Each mapped read is classified into one of three categories (Fig. 1b, Methods): “haplotype-resolved” (uniquely mapped to one haplotype), “position-only” (the read maps to a unique position but contains no haplotype-specific markers, so its haplotype cannot be determined), or “multi-mapped” (neither position nor haplotype uniquely determined). Using these mappings, PRCGAP performs downstream analyses, including somatic point mutation and SV calling, copy-number analysis, and methylation analysis.

Somatic variants are called using haplotype-resolved and position-only reads, while multi-mapped reads generally do not contribute since they have low mapping quality. Haplotype-resolved reads are used to phase each variant to its haplotype of origin, whereas position-only reads can also align to the same position on the other haplotype, generating redundant unphased calls. PRCGAP consolidates these redundant calls by comparing variant-flanking sequences between haplotypes (Methods). In addition, PRCGAP filters out false-positive calls arising from read mis-mapping or assembly errors (Methods). To enable comparison with standard-reference-based variant calling and to identify variants inaccessible to such analyses, PRCGAP lifts over genomic coordinates from the personalized reference genome to GRCh38 and T2T-CHM13, classifying each as liftable or non-liftable (Fig. 1c, Methods).

### Construction and evaluation of personalized reference genomes

We applied PRCGAP to eight cancer cell lines and three clinical samples, together with their matched normal samples (Fig. 1a and Supplementary Table 1). The cell lines included three pairs from the Cancer Standards Long-read Evaluation panel (H1437, H2009, and HCC1937)^19^, COLO829 from the Somatic Mosaicism across Human Tissues consortium^14,21^, HG008 from the Genome in a Bottle consortium^20^, and three newly sequenced pairs (H209, H2126, and HCC1954). The clinical samples included three newly generated ALL samples (NH-ALL-001, UTPT2T-02, and UTPT2T-04). De novo assemblies of the matched normal samples were generated using hifiasm and/or Verkko depending on the available phasing data type (Methods). Across all datasets, the assemblies yielded total contig sizes of approximately 3 Gb with high contiguity (median N50, 110.9 and 145.7 Mb for hifiasm and Verkko, respectively; Fig. 1d, Supplementary Table 2) and high accuracy (median QV, 60.5 and 60.7 for hifiasm and Verkko, respectively; Supplementary Fig. 1a). The assemblies contained a median of 9 (range, 4–15) and 16 (range, 7–23) T2T chromosomes for hifiasm and Verkko, respectively (Fig. 1e). The assemblies also correctly covered approximately 99% of known single-copy genes^22^ (Supplementary Fig. 1b and Supplementary Tables 2 and 3). Phasing accuracy of the diploid assemblies, evaluated using Hi-C or trio data when applicable (Methods), showed a median switch error rate of 0.17% (ranges, 0.04–0.72% for hifiasm and 0.06–0.63% for Verkko) and a median Hamming error rate of 0.59% (ranges, 0.15–1.26% for hifiasm and 0.19–2.11% for Verkko; Supplementary Fig. 1c). We estimated that a median of 99.7% (98.4–99.9%) of each haplotype assembly was correctly assembled using Flagger^9^, Inspector^23^, and NucFlag^24,25^ (Fig. 1f, Supplementary Tables 4–6, Supplementary Figs. 2a–c).

We further evaluated repeat annotations, including centromeres and segmental duplications. The assemblies harbored a median of 35 accurately assembled centromeres and exhibited a segmental duplication ratio of 6.17% (Supplementary Fig. 2d, e). These results were consistent with or superior to those reported in previous studies^26,27^. Taken together, these evaluations confirmed that both hifiasm and Verkko produce high-quality assemblies suitable as personalized reference genomes. Between the two assemblers, Verkko achieved higher contiguity and T2T completeness with comparable base accuracy and phasing accuracy. We therefore used Verkko assemblies as personalized reference genomes for downstream analyses in PRCGAP.

### Haplotype-aware mapping to the personalized reference genomes

After the construction of the personalized reference genomes, PRCGAP performed haplotype-aware mapping of long-read sequencing data from both tumor and normal samples (Fig. 1b, Supplementary Fig. 3a). Across HiFi and ONT alignments, a median of 78.1% of mapped reads were haplotype-resolved with high confidence (Supplementary Fig. 3b), 19.9% were position-only, and 1.9% were multi-mapped (Supplementary Table 7). The multi-mapped proportion was substantially lower than the 7.1% obtained when mapping to T2T-CHM13, demonstrating the benefit of mapping to personalized reference genomes using haplotype-specific markers (Supplementary Fig. 3c and Supplementary Table 7).

We further evaluated mappability of the personalized reference genomes by calculating the multi-mapped read ratio in 10-kb windows. Even when using personalized reference genomes, multi-mapped reads were enriched in parts of centromeric or satellite regions and segmental duplications (Supplementary Fig. 3d, e and Supplementary Table 8). For centromere or satellite regions, the multi-mapping ratio was extremely high in ribosomal DNA regions (Supplementary Fig. 3d). Segmental duplications with high similarity tended to have higher proportions of multi-mapped reads (Supplementary Fig. 3e). These results indicate that PRCGAP enables haplotype-aware read mapping across centromeric and other repetitive regions, with only specific subregions of insufficient sequence divergence (such as rDNA arrays) remaining challenging.

### Somatic variant detection in cancer cell lines

From these haplotype-aware mappings, PRCGAP detected somatic point mutations and SVs in the eight cancer cell lines using DeepSomatic^19^ and nanomonsv^28^, respectively. For each variant, we examined whether it was phased to a haplotype (“phasability”) and whether its genomic coordinates were liftable to GRCh38 or T2T-CHM13 (“liftability”; Fig. 1c). Because SVs involve two breakpoints, we additionally introduced a “Partial” category in each classification: for phasability, when only one breakpoint was phased; for liftability, when only one breakpoint was liftable.

PRCGAP detected a median of 35,251 SNVs (range, 10,180–144,968) and 4,635 short indels (range, 916–8,252; Fig. 2a and Supplementary Fig. 4a). Over 95% of SNVs and short indels were phasable and over 74% of their genomic coordinates were liftable to GRCh38- or T2T-CHM13-based coordinates (Fig. 2b, Supplementary Figs. 4b and 5a, b, and Supplementary Table 9). PRCGAP also detected a median of 382 SVs (range, 61–1,106; Fig. 2c). Over 76% of SVs were phasable (range, 76.0–93.8%) and over 70% of their genomic coordinates were liftable to GRCh38- or T2T-CHM13-based coordinates (range, 70.9–95.0%; Fig. 2d, Supplementary Fig. 5c, and Supplementary Table 9).

**Figure 2.**
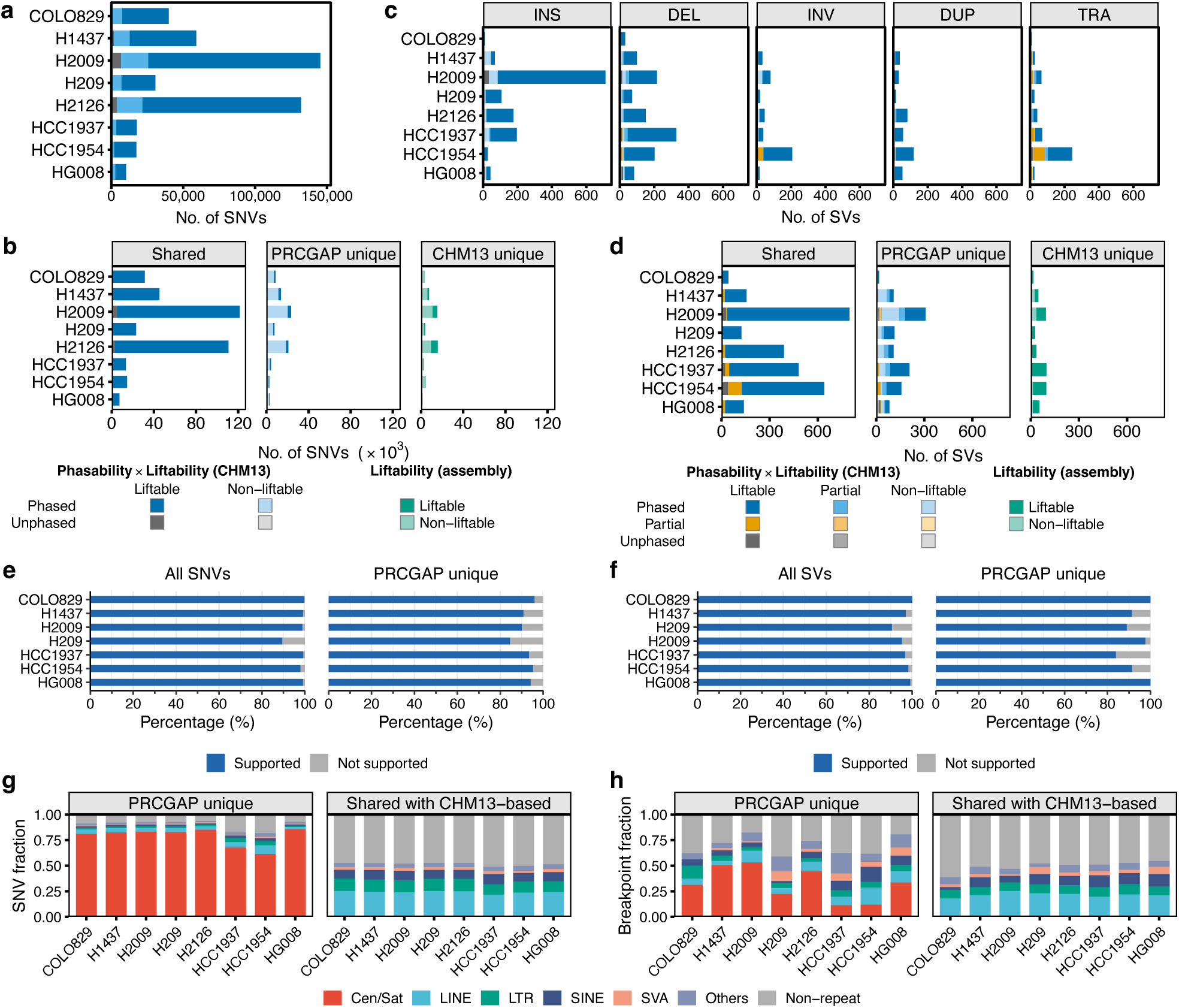
Somatic mutation calling by PRCGAP and comparison with T2T-CHM13-based approach across cancer cell lines. (a) Number of somatic SNVs detected by PRCGAP per cell line, stratified by phasability (Phased or Unphased) and liftability to T2T-CHM13 coordinates (Liftable or Non-liftable). (b) Comparison of somatic SNVs between PRCGAP and the T2T-CHM13-based pipeline, classified as shared (Shared), unique to PRCGAP (PRCGAP unique), or unique to the T2T-CHM13-based pipeline (CHM13 unique). In the Shared and PRCGAP unique panels, SNVs are colored by phasability and liftability (CHM13); in the CHM13 unique panel, by liftability to the personalized reference. (c) Number of somatic SVs detected by PRCGAP per cell line, faceted by SV type (INS, DEL, INV, DUP, and TRA). (d) Comparison of somatic SVs between PRCGAP and the T2T-CHM13-based pipeline, classified as in (b). SVs are colored by phasability and liftability, with the additional “Partial” category indicating that only one of the two breakpoints is phased (for phasability) or liftable (for liftability). (e, f) Validation of PRCGAP somatic SNVs (e) and SVs (f) using Illumina whole-genome sequencing data. For each of seven cell lines with available Illumina data, the percentage of mutations supported by Illumina data is shown for all PRCGAP mutations (left) and the subset unique to PRCGAP (right). (g, h) Fraction of somatic SNVs (g) and SV breakpoints (h) annotated by RepeatMasker, compared between PRCGAP-unique mutations (left) and mutations shared with the T2T-CHM13-based pipeline (right). Repeat classes: Cen/Sat, LINE, LTR, SINE, SVA, Others, and Non-repeat.

To evaluate the variant-calling performance of PRCGAP, we compared somatic variants detected by PRCGAP with those detected using GRCh38- and T2T-CHM13-based approaches. Somatic SNVs, short indels, and SVs detected by PRCGAP overlapped with those from the GRCh38-and T2T-CHM13-based approaches, with median overlap rates of 96.4% (94.2–98.6%) for SNVs, 91.8% (85.9–95.5%) for short indels, and 86.7% (72.4–93.8%) for SVs (Fig. 2b, d, Supplementary Figs. 4b and 5, and Supplementary Table 10). Variants detected only by the GRCh38- or T2T-CHM13-based approaches tended to show low frequencies and to be located in highly repetitive regions, such as centromeric and segmentally duplicated regions (Supplementary Figs. 6 and 7).

To confirm somatic variants detected by PRCGAP, we performed k-mer-based validation using Illumina whole-genome sequencing data (Methods). For the seven cell lines with available Illumina data, 89.4–99.6% (median 99.1%) of SNVs detected by PRCGAP were supported by Illumina data (Fig. 2e), and 90.5–100.0% (median 97.0%) of SVs were supported (Fig. 2f). Focusing on PRCGAP-unique variants that were not detected using the standard reference genomes, 84.5–95.9% (median 93.3%) of SNVs and 83.9–100.0% (median 91.4%) of SVs were supported by Illumina data. These results indicate that somatic variants detected by PRCGAP, including those unique to the personalized reference approach, are largely reliable.

PRCGAP detected a substantial fraction of SNVs and SVs that were missed by standard reference-based approaches: a median of 23.4% (15.8–29.9%) of SNVs and 32.6% (25.3–57.6%) of SVs were PRCGAP-unique (Fig. 2b, d, Supplementary Fig. 5a, c, and Supplementary Table 10). We then explored the characteristics of PRCGAP-unique variants by annotating repeat and segmental duplication information using RepeatMasker^29^ and SEDEF^30^. PRCGAP-unique variants were enriched in regions annotated as centromeres (Fig. 2g, h, Supplementary Fig. 8). Moreover, PRCGAP-unique somatic point mutations were enriched in segmental duplication regions, including regions with sequence similarity exceeding 80% (Supplementary Fig. 9). These results demonstrate that PRCGAP enables the detection of somatic variants in highly repetitive genomic regions, including centromeres and segmental duplications.

### Genomic overview of cancer cell lines

In addition to point mutation and SV calling, PRCGAP generates haplotype-resolved copy-number profiles (Fig. 3a and Supplementary Fig. 10). The cancer cell lines exhibited diverse patterns of copy-number alterations across both haplotypes, with frequent oscillations between gain, loss, and copy-number neutral segments. By integrating these results, we constructed haplotype-resolved genomic overviews of cancer genomes encompassing cancer-associated point mutations, SV breakpoints, and copy number (Fig. 3b and Supplementary Fig. 11).

**Figure 3.**
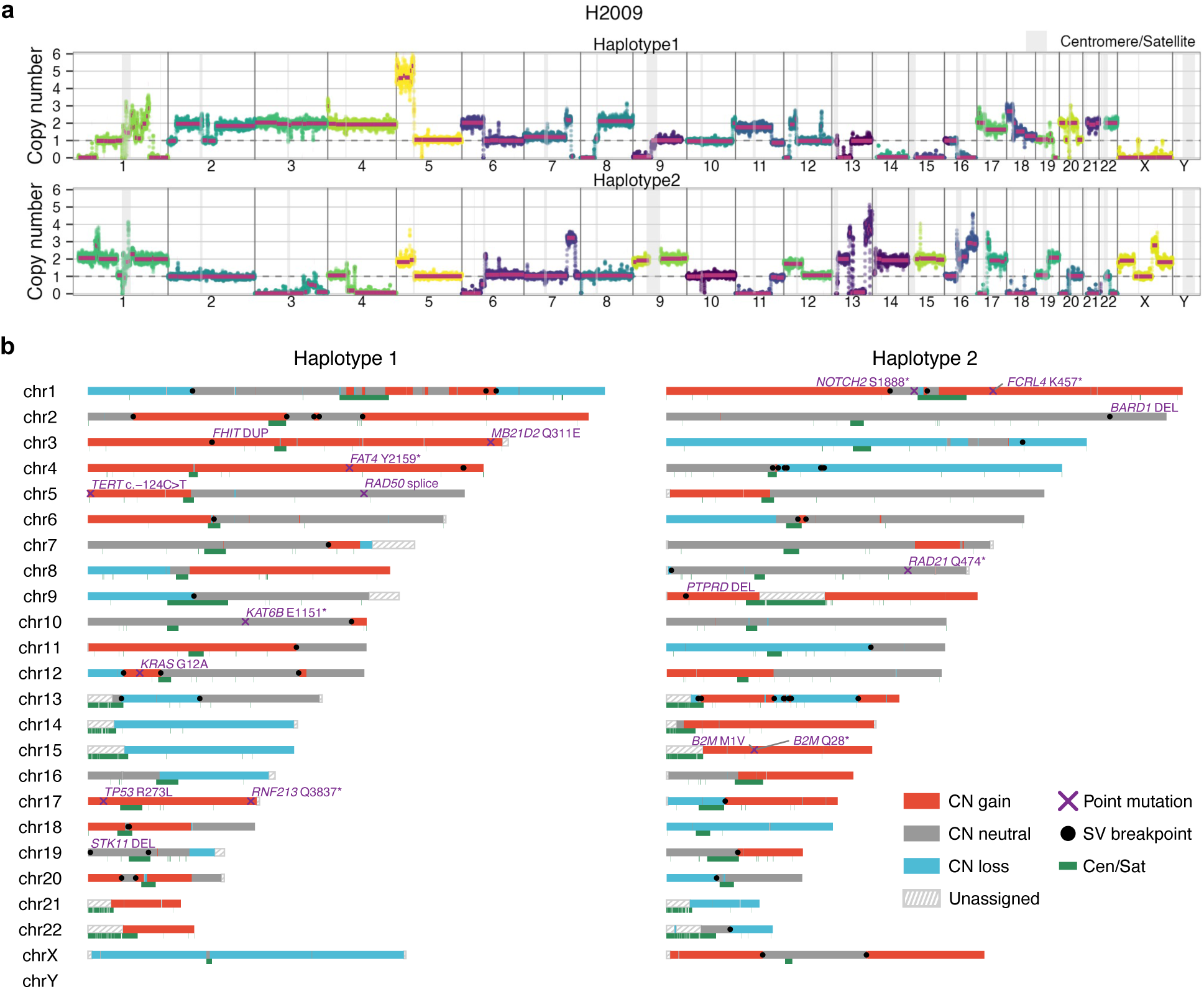
Haplotype-resolved genomic overview of the H2009 cancer cell line. (a) Haplotype-resolved copy-number profile of the H2009 cell line, shown separately for haplotype 1 (top) and haplotype 2 (bottom). The y-axis indicates the estimated copy number, and the x-axis represents genomic positions ordered by chromosome. Points are colored by assembled contigs, and centromere/satellite regions are shaded in gray. (b) Haplotype-resolved genomic overview of the H2009 cell line, integrating cancer-associated point mutations, SV breakpoints, and copy-number status. Chromosome segments are colored by copy-number status (CN gain, red; CN neutral, gray; CN loss, light blue) or coverage status with respect to T2T-CHM13 (Unassigned, hatched gray). Centromere/Satellite (Cen/Sat) regions are marked in green. SV breakpoints are shown as black dots, and cancer-associated somatic point mutations are shown as purple crosses and labeled with the affected gene name.

PRCGAP detected 46 of 56 (82.1%) previously reported cancer-associated point mutations^20,31^ across the eight cell lines (Methods, Supplementary Table 11) and revealed their co-occurrence with copy-number changes and SV breakpoints on the same haplotype^32^. Eight out of the ten undetected mutations were also missed by GRCh38-based analysis, suggesting intrinsic variant-calling challenges rather than a PRCGAP-specific limitation (Supplementary Table 11).

Tumor suppressor inactivation typically involves a somatic mutation combined with loss of heterozygosity (LOH) of the wild-type allele, whereas oncogene activation is often accompanied by copy-number gain of the mutant allele^33^. PRCGAP enabled visualization of both configurations at single-haplotype resolution. Consistent with the tumor suppressor pattern, *TP53* mutations were detected in seven out of eight cancer cell lines, all accompanied by inactivation of the wild-type allele. In five of these (H1437, H209, H2009, H2126, and HCC1937), the mutant haplotype exhibited copy-number gain while the wild-type haplotype was lost, indicating LOH accompanied by mutant allele amplification. In the HG008 cell line, the wild-type haplotype was lost while the mutant haplotype was retained as a single copy, indicating hemizygous LOH. In the HCC1954 cell line, *TP53* mutations were identified on both haplotypes, indicating biallelic inactivation (Supplementary Fig. 11). Consistent with the oncogene pattern, *KRAS* mutations co-occurring with copy-number gain of the mutant haplotype were observed in the H2009 (G12A, lung adenocarcinoma) and HG008 (G12V, pancreatic ductal adenocarcinoma) cell lines, with SV breakpoints at the borders of the amplified regions (Fig. 3b and Supplementary Fig. 11). *KRAS* mutant allele-specific amplification has been associated with poor prognosis in both lung cancer and pancreatic cancer^34,35^. These results demonstrate that haplotype-resolved analysis by PRCGAP enables direct assessment of allele-specific copy-number changes and their relationship to cancer-associated mutations and SVs.

Among the cell lines, HCC1954 exhibited a particularly complex rearrangement pattern. Haplotype 1 of chromosome 5 and haplotype 2 of chromosome 8 showed pronounced copy-number oscillations and a high density of SV breakpoints, consistent with a chromothripsis-like event^36^ (Supplementary Fig. 12a–c). In these same regions, candidate kataegis loci^37^ were identified by inter-mutational distances, with C>T and C>G substitutions enriched on the q arm of chr5 haplotype 1 and the q arm of chr8 haplotype 2. Mutational signature analysis further revealed SBS2 and SBS13 signatures attributable to APOBEC activity^38^ (Supplementary Fig. 12d, e). The co-localization of SV breakpoints and APOBEC-associated point mutations suggests concurrent chromothripsis and kataegis in HCC1954.

### Variants involving centromere regions in cancer cell lines

During the evaluation of PRCGAP variant calling, we found that PRCGAP-unique variants were enriched in centromeric regions annotated by RepeatMasker (Fig. 2g, h). To further characterize these variants, we annotated SNVs and SV breakpoints by centromeric satellite class (α-satellite, human satellite, and other centromeric satellites; Fig. 4a)^39^. The α-satellite regions are subdivided into active higher-order repeat (HOR), inactive HOR, and monomeric α-satellite, and the active HOR harbors locally hypomethylated loci known as centromere dip regions (CDRs), the putative sites of kinetochore formation^40^ (Fig. 4a). Using this annotation, we first compared somatic SNV rates across centromeric satellite classes in the six cell lines whose centromeres passed our quality-control criteria (HCC1954 and HG008 were excluded owing to the limited number of available centromeres; Methods). The somatic SNV rate was elevated within the CDR, reaching 4.3-fold that of centromere-flanking unique regions (Fig. 4b and Supplementary Fig. 13). This elevated somatic mutation rate in the CDR is consistent with the trend recently reported for intergenerational de novo mutations, although our analysis concerns cancer somatic mutations^41^.

**Figure 4.**
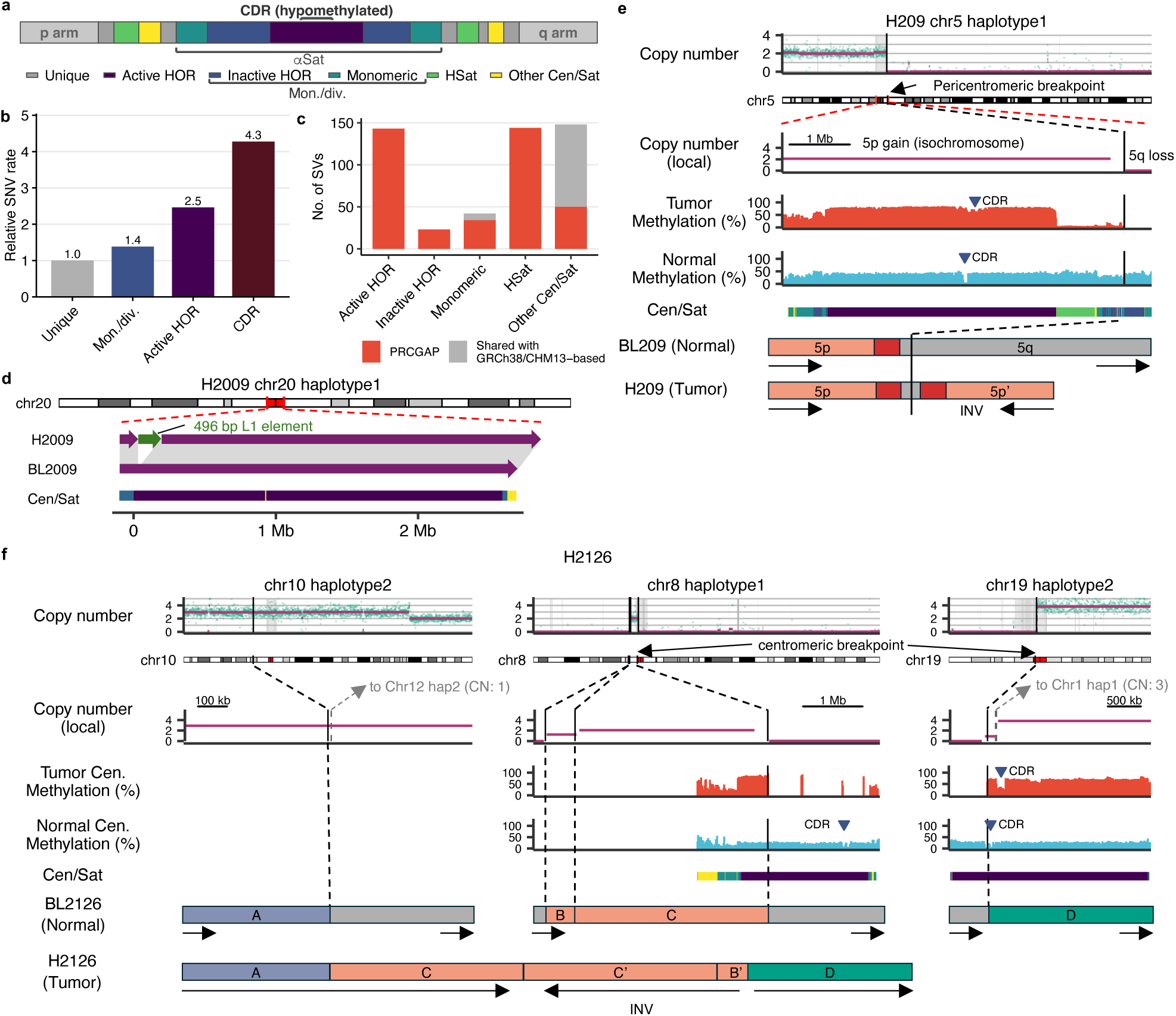
Variants involving centromere regions in cancer cell lines. (a) Schematic of centromere structure. The α-satellite (αSat) region is subdivided into active higher-order repeat (Active HOR), inactive HOR, and monomeric classes, and is flanked by human satellite (HSat) and other centromeric satellite (Other Cen/Sat) regions. The locally hypomethylated centromere dip region (CDR) is located within the active HOR. (b) Relative somatic SNV rate in centromere-annotated regions compared to centromere-flanking unique regions. (c) Number of somatic SVs in the H2009 cell line overlapping centromere-annotated regions, stratified by centromere annotation class (Active HOR, Inactive HOR, Monomeric, HSat, and Other Cen/Sat). Bars are colored by whether the SV was unique to PRCGAP (red) or shared with the GRCh38/T2T-CHM13-based pipelines (gray). (d) A solo L1 insertion within the active HOR region of chromosome 20 haplotype 1 in the H2009 cell line. A 496 bp L1 element is inserted in H2009 but absent from BL2009. (e) Isochromosome of the p arm of chromosome 5 haplotype 1 in the H209 cell line. Global and local copy-number profiles of chromosome 5 haplotype 1 show 5p gain (isochromosome) and 5q loss, together with tumor and normal DNA methylation tracks and the CDR (blue triangle) annotation. The bottom panel shows the reconstructed structure in BL209 (normal) and H209 (tumor). (f) Complex rearrangement in the H2126 cell line involving chromosome 10 haplotype 2 (segment A), chromosome 8 haplotype 1 (segments B and C), and chromosome 19 haplotype 2 (segment D) with global and local copy-number profiles and DNA methylation frequencies in centromere regions. The bottom panels show the reconstructed structure in BL2126 (normal) and H2126 (tumor).

We next examined SVs involving centromere regions. We found that PRCGAP could identify SV breakpoints in α-satellite and human satellite regions (Supplementary Table 12). Most of these SVs were undetectable by GRCh38- or T2T-CHM13-based pipelines (Fig. 4c and Supplementary Fig. 14), suggesting that SV detection in these regions is more challenging with standard reference-based approaches and that PRCGAP can more effectively resolve centromere-involving SVs.

PRCGAP revealed diverse centromere-involving SVs across cell lines. In the H2009 cell line, we identified a solo L1 insertion within the active HOR region of chromosome 20 haplotype 1 (Fig. 4d and Supplementary Fig. 15a) and a derivative chromosome involving chromosome 6 haplotype 1 and chromosome 11 haplotype 2, where the chromosome 6 breakpoint was located in the active HOR (Supplementary Fig. 15b). Analysis of the H209 cell line revealed a pericentromeric breakpoint associated with isochromosome 5p in chromosome 5 haplotype 1, and the resulting isochromosome retained the CDR (Fig. 4e). A more complex rearrangement was observed in the H2126 cell line, involving chromosome 10 haplotype 2 and the active HOR regions of chromosome 8 haplotype 1 and chromosome 19 haplotype 2 (Fig. 4f). In the putative structure, the p arm of chromosome 10 haplotype 2 connected to the p arm of chromosome 8 haplotype 1 with an inversion at the active HOR, followed by translocation into the active HOR of chromosome 19 haplotype 2. This rearrangement harbored a CDR derived from chromosome 19 haplotype 2, and notably, the CDR of H2126 at this locus shifted toward the q arm relative to BL2126 (Fig. 4f). These results demonstrate that PRCGAP enables the detection and characterization of centromere-involving SVs and copy-number alterations, and their interpretation as part of complex genomic rearrangements.

### SVs involving telomere sequences in cancer cell lines

To explore telomere-related SVs in the cancer cell lines, we examined both canonical (two-breakpoint) SVs and single-breakend SVs, in which only one breakpoint could be localized while the partner remained unmapped, and extracted those overlapping telomeric or subtelomeric regions or harboring telomere repeat sequences (Fig. 5a). In total, we identified 13 telomere-related SVs across the eight cancer cell lines (Fig. 5b and Supplementary Table 13), classified into three categories: telomere-associated translocations (n = 4), subtelomeric SVs (n = 7), and telomere-containing single breakends (n = 2). Of these, 11 were uniquely detected by PRCGAP (Fig. 5b).

**Figure 5.**
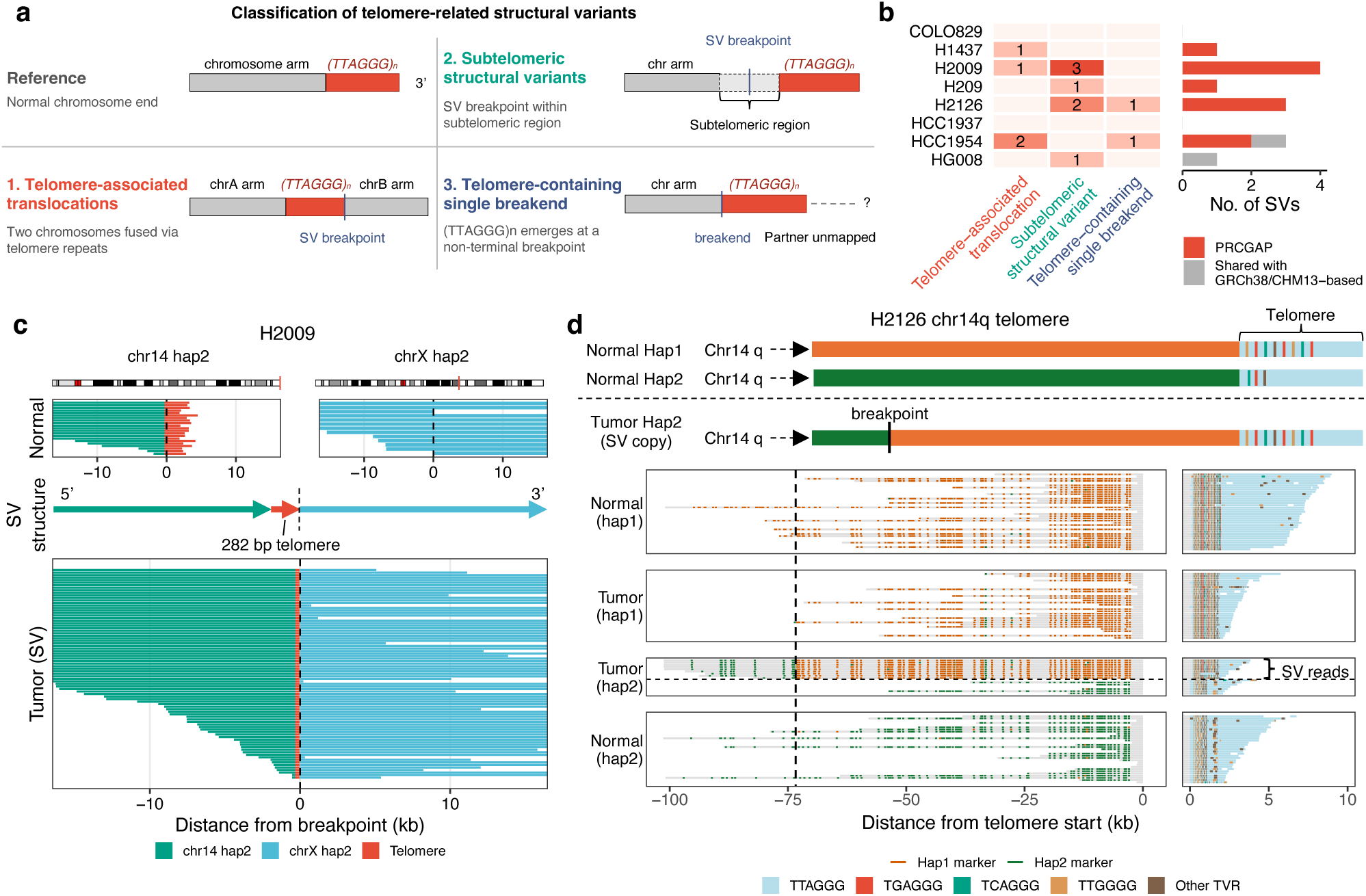
Telomere-related somatic SVs in cancer cell lines. (a) Schematic classification of telomere-related SVs into three categories: telomere-associated translocations (two chromosomes fused via telomere repeats), subtelomeric SVs (breakpoint within a subtelomeric region), and telomere-containing single breakends (telomere repeats at a non-terminal breakpoint with an unmapped partner). A normal chromosome end is shown for reference. (b) Number of telomere-related SVs detected by PRCGAP across cancer cell lines (rows), classified into the three categories defined in a (columns). Heatmap color intensity and the number in each cell indicate the SV count. The right bar plot shows the per-cell-line total, colored by PRCGAP-unique (red) or shared with the GRCh38/T2T-CHM13-based pipelines (gray). (c) Telomere-associated translocation between chromosome 14 haplotype 2 telomere and chromosome X haplotype 2 q arm in the H2009 cell line. Chromosome ideograms (top) show the breakpoint positions in red. The Normal panel (BL2009) shows sequence reads mapped around each breakpoint separately, and the Tumor (SV) panel (H2009) shows chimeric reads spanning the junction. Between them, a schematic SV structure is shown. Arrows indicate the 5’-to-3’ direction of each segment. The x-axis shows distance from the breakpoint (0 kb, vertical dashed line). Reads are colored by origin: chr14 haplotype 2 (green), chrX haplotype 2 (light blue), and telomere repeat sequence (red). (d) Inter-haplotype recombination at the chromosome 14 q-arm telomere in the H2126 cell line. Top schematics show the normal haplotype 1 telomere (Normal Hap1), the normal haplotype 2 telomere (Normal Hap2), and the SV copy of tumor haplotype 2, labeled Tumor Hap2 (SV copy), in which the distal ∼75-kb segment has been replaced by haplotype 1-derived sequence at the breakpoint. Sequence reads mapped around telomere regions are shown with haplotype-specific markers (haplotype 1: orange, haplotype 2: green) and telomere variant repeat (TVR) composition (TTAGGG: light blue, TGAGGG: red, TCAGGG: teal, TTGGGG: orange, and Other TVR: brown). In the Tumor (hap2) panel, the dashed line separates SV-supporting reads (top, indicated by the right-side bracket) from parental hap2 reads (bottom).

Among these, the H2009 cell line harbored a previously reported translocation between the chromosome 14 haplotype 2 telomere and the chromosome X haplotype 2 q arm^28^, with 282 bp of telomere repeats retained at the junction (Fig. 5c). The remaining three telomere-associated translocations are shown in Supplementary Fig. 16.

One telomere-containing single breakend in H2126, located at the chromosome 14 haplotype 2 telomere, showed evidence of inter-haplotype recombination (Fig. 5d). Haplotype-specific marker analysis localized the breakpoint approximately 75 kb from the telomere, where supporting reads transitioned from haplotype 2-specific to haplotype 1-specific markers. The telomere variant repeat composition at the recombined region also matched that of the haplotype 1 telomere, identifying haplotype 1 as the donor template (Fig. 5d). This continuous ∼75-kb donor-derived tract that includes the telomere is reminiscent of break-induced replication, a long-tract recombination mechanism associated with alternative lengthening of telomeres^42^.

### L1 source site detection in cancer cell lines

L1 elements are autonomous retrotransposons that copy themselves into new genomic locations. Identifying the source site, the active L1 element from which an insertion derives, is essential for understanding L1 retrotransposition in cancer^43–45^. L1 insertions are classified into solo L1 insertions and transductions (TDs; classified as partnered or orphan). TDs carry 3′ downstream sequences that enable source site identification by alignment, whereas solo L1 insertions lack such sequences, making source site detection challenging. A recent study addressed this challenge by leveraging the T2T-CHM13 reference genome^46^, although source sites absent from T2T-CHM13 are challenging to identify. To further improve source site detection, we developed a marker-based approach using personalized reference genomes and L1 insertions detected by PRCGAP (Fig. 6a and Methods). We applied this pipeline to all eight cancer cell lines (Fig. 6b and Supplementary Fig. 17a). In total, we detected 427 solo L1 insertions, 142 partnered TDs, and 52 orphan TDs across the eight cell lines. Using a mean of 14 marker sequences per L1 element (9,889 markers per personalized reference genome), we identified source sites for 190 (44.5%) solo L1 insertions (Supplementary Fig. 17a, b), delineating 63 active L1 source sites in total.

**Figure 6.**
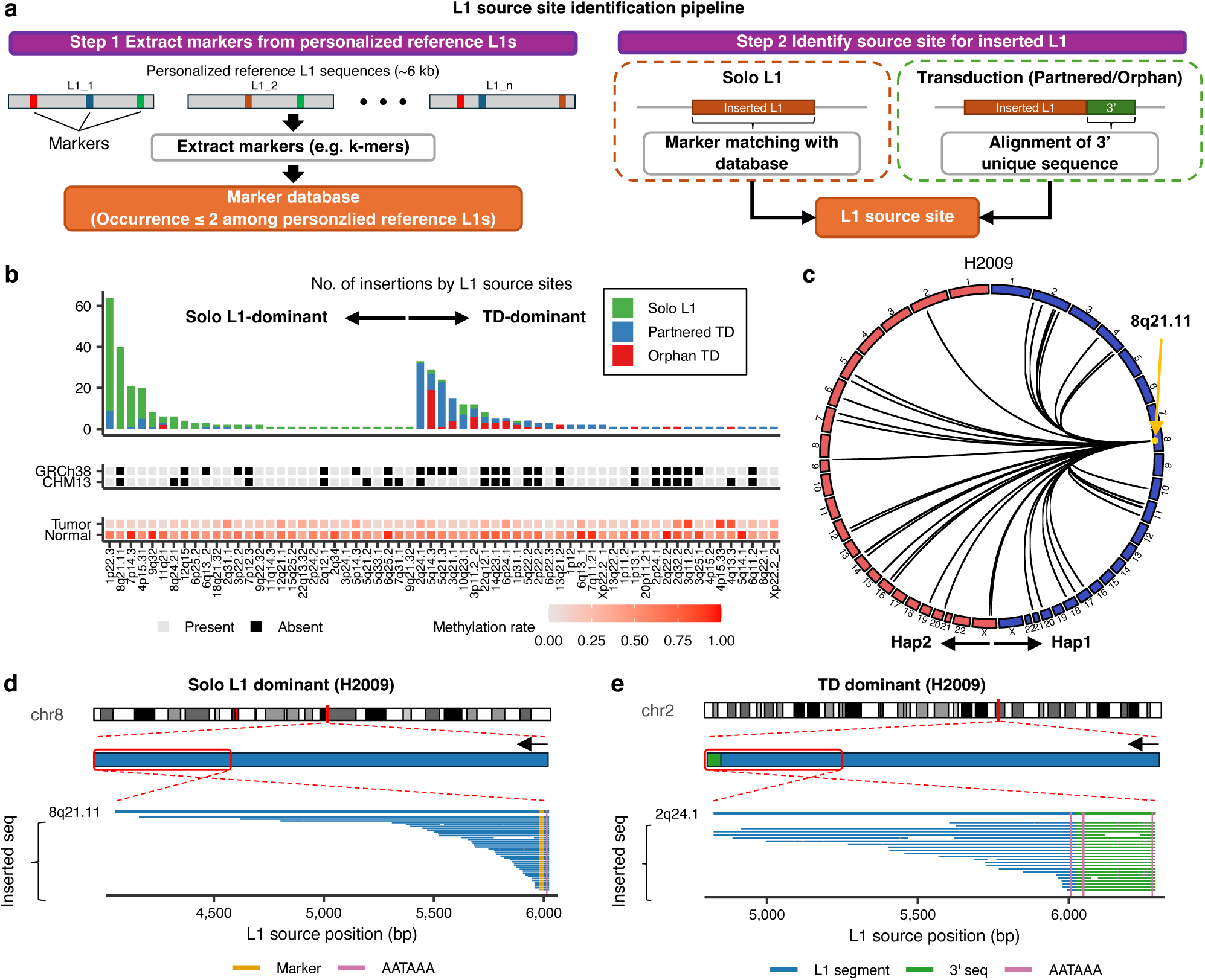
Identification of L1 source sites using the personalized reference genome across cancer cell lines. (a) Schematic of the pipeline to identify L1 source sites for solo L1 insertions and transductions (TDs; see Methods). (b) Number of somatic L1 insertions per source site across cancer cell lines, stratified by insertion type (solo L1, partnered TD, and orphan TD). Source sites are ordered and grouped into those dominated by solo L1 insertions (left) and those dominated by TDs (right). Below the bar plot, heatmap tracks indicate the presence (gray) or absence (black) of each source site in the GRCh38 and T2T-CHM13 reference genomes, and the DNA methylation level at each source site in matched tumor and normal samples. (c) Circos plot of L1 insertions originating from the 8q21.11 source site on haplotype 1 in the H2009 cell line. Lines connect the 8q21.11 source site (arrow) to each target insertion site across the haplotype 1 and haplotype 2 genomes. (d) Example of a solo L1-dominant source site at 8q21.11 (haplotype 1) in the H2009 cell line. The alignment of inserted L1 sequences (horizontal bars) to the reference L1 at the source locus is shown together with rare k-mer markers within the inserted sequences and the internal polyadenylation signal (AATAAA) within the L1 element. (e) Example of a TD-dominant source site at 2q24.1 in the H2009 cell line. The alignment of inserted sequences shows that L1 segments extend into the 3′ downstream sequence, with the polyadenylation signal (AATAAA) located outside the L1 element within the downstream sequence.

To evaluate the performance of our source site detection pipeline, we applied this approach to 48 TDs in H2009 whose source sites had already been identified using the alignment-based method, and 45 (93.8%) were correctly assigned.

Notably, the personalized reference genomes enabled identification of source sites that are not represented in the standard reference. At the source-site level, of the 63 active L1 source sites, 17 corresponded to L1 elements absent from both GRCh38 and T2T-CHM13 (Fig. 6b) and were also detected as structural variations in the Human Genome Structural Variation Consortium Phase 3 and the 1000 Genomes Project ONT datasets^27,47^. L1 sequences of the personalized reference genomes differed from those of T2T-CHM13 (Median edit distance: 6.5 bp; Supplementary Fig. 17c). At the marker level, even for source sites present in T2T-CHM13, 18.4% of markers supporting source sites were absent from T2T-CHM13 (Supplementary Fig. 17d). For example, in H2009, the cell line with the highest somatic L1 burden among the eight cancer cell lines (Supplementary Fig. 17a), the most active L1 element at 1p22.3 generated 55 solo L1 insertions supported by two markers, neither of which was present in T2T-CHM13 (Supplementary Fig. 17e). Collectively, these results demonstrate that a personalized reference genome improves source site detection and enables the identification of novel active L1 source sites that would be missed using a standard reference genome.

Having established the importance of a personalized reference genome for accurate source site identification, we next examined whether L1 source sites exhibit distinct mechanistic patterns. We classified source sites as “Solo L1-dominant” or “TD-dominant” based on the predominant insertion type (Fig. 6b). The second most active L1 element at 8q21.11 generated 40 solo L1 insertions broadly distributed across diverse chromosomes and haplotypes, representing a Solo L1-dominant source site (Fig. 6c, d). In contrast, the active L1 element at 2q24.1 generated 31 partnered TDs with 3′ downstream sequences of the L1 element, representing a TD-dominant source site (Fig. 6e). These findings suggest that L1 source sites can be grouped by their predominant insertion type, providing new insight into the mechanistic diversity of L1 retrotransposition.

### Application of PRCGAP to clinical sequencing data

To assess whether PRCGAP is applicable to clinical sequencing data, we analyzed newly generated data from three pediatric B-ALL patients (Supplementary Table 1). The three cases represented distinct molecular subtypes of B-ALL^48,49^: near-haploid ALL (NH-ALL-001), *DUX4*-rearranged ALL (UTPT2T-02), and hyperdiploid ALL (UTPT2T-04). Across the three cases, PRCGAP detected 1,071–1,542 SNVs, 260–494 short indels, and 19–30 SVs, with chromosome numbers of 26 (NH-ALL-001), 46 (UTPT2T-02), and 65 (UTPT2T-04), consistent with their subtype classifications^50–52^.

Previous reports suggest that chromosomal losses in near-haploid ALL and gains in hyperdiploid ALL do not show consistent parental-origin bias, arguing against a major role for genomic imprinting in these subtypes^51,53,54^. We examined whether this pattern was also observed in NH-ALL-001 and UTPT2T-04 using PRCGAP.

For NH-ALL-001, copy-number analysis identified allelic loss of each chromosome except chromosomes 6, 10, 21, and X, consistent with a previous report^51^ (Fig. 7a). Parent-of-origin analysis based on imprinted differentially methylated regions (iDMRs)^55^ revealed that the parental origin of lost chromosomes varied by chromosome, consistent with prior reports of random parental loss^56^ (Fig. 7a). PRCGAP also identified breakpoints of the 1q gain in the maternal haplotype involving the human satellite region (Fig. 7b and Supplementary Fig. 19a).

**Figure 7.**
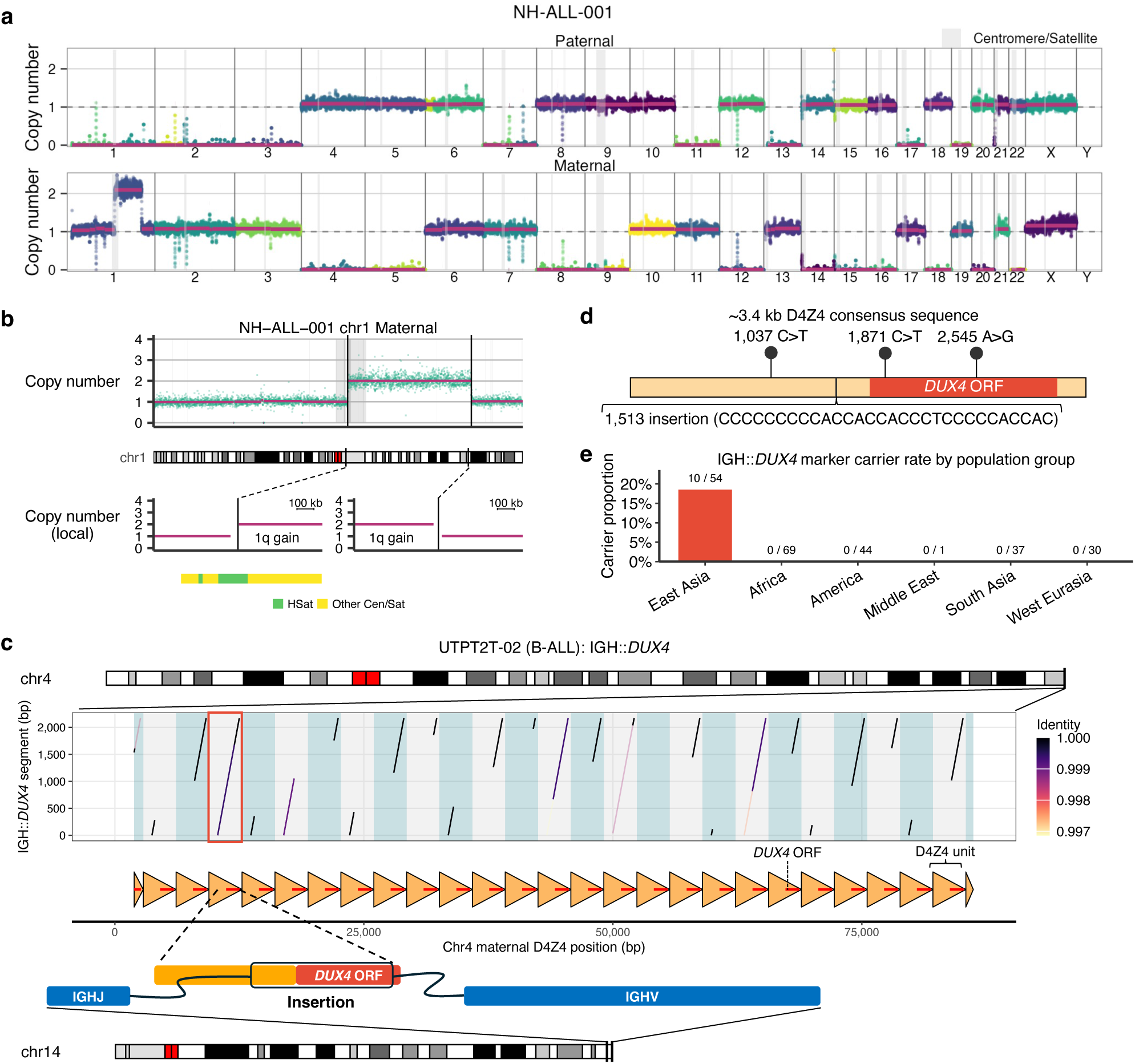
Haplotype-resolved analysis of pediatric B-ALL samples. (a) Haplotype-resolved copy-number profile of the NH-ALL-001 sample after parental origin assignment, shown separately for the paternal (top) and maternal (bottom) haplotypes. The y-axis indicates the estimated copy number, and the x-axis represents genomic positions ordered by chromosome. Centromere/satellite regions are shaded in gray. (b) Detail of the 1q gain on chromosome 1 maternal haplotype in NH-ALL-001. The global and local copy-number profile of chromosome 1 shows the duplication. The Cen/Sat annotation below shows that one breakpoint overlaps a human satellite (HSat) region, indicating a pericentromeric tandem duplication. (c) Structure of the IGH::*DUX4* fusion in the UTPT2T-02 sample. Dot plot shows an alignment of the inserted *DUX4* segment (y-axis) against the D4Z4 repeat array on the maternal chromosome 4 (x-axis), divided into 2-kb segments and colored by sequence identity (threshold: 0.995). Orange arrows below indicate the D4Z4 repeat units, with the *DUX4* open reading frame (ORF) annotated. The bottom schematic shows the reconstructed fusion structure on chromosome 14, in which a D4Z4 repeat sequence containing the *DUX4* ORF is inserted between IGHJ and IGHV segments. (d) Variants identified in the inserted *DUX4* segment relative to the ∼3.4 kb D4Z4 consensus sequence. (e) Carrier rate of the IGH::*DUX4* markers across six population groups. The bar height indicates the proportion of individuals carrying the markers, and the fraction of carriers is labeled above each bar. The markers were detected only among the East Asian population (10/54).

For UTPT2T-04, the personalized reference genome was constructed from trio data, so parental origins were assigned to chromosomes during de novo assembly. We observed a hyperdiploid karyotype with copy-number gain of all chromosomes except chromosomes 1, 7, 9, 13, 15, 19, and X (Supplementary Fig. 18a and Supplementary Fig. 19b). Chromosomes 10 and 21 exhibited gains of both haplotypes, indicating that both the paternal and maternal copies were duplicated. The remaining chromosomes with trisomy included gains of both paternal and maternal homologs across chromosomes, consistent with previous reports suggesting that genomic imprinting does not play a major pathogenetic role in hyperdiploid ALL^53,54^. Notably, we identified a chromosome 1q gain in the paternal haplotype with breakpoints proximal to those of NH-ALL-001 (Supplementary Fig. 19b), an alteration recurrent in B-ALL and other hematological malignancies with established prognostic relevance^57,58^.

For UTPT2T-02, no copy-number alteration was observed across the genome (Supplementary Fig. 18b and Supplementary Fig. 19c). The IGH::*DUX4* fusion gene was identified in this case, with its genomic structure analyzed in detail below.

These results demonstrate that PRCGAP is applicable to clinical cancer sequencing data, elucidating cancer genome structures in a haplotype-resolved manner.

### IGH::*DUX4* fusion gene structure in a pediatric B-ALL sample

The IGH::*DUX4* fusion is a recurrent oncogenic event in B-ALL^48,59,60^, and recent long-read sequencing-based approaches have enabled its systematic detection^61^. However, resolving the genomic structure of the IGH::*DUX4* fusion is challenging because the *DUX4* gene is located in a highly repetitive D4Z4 macrosatellite array at the subtelomeric regions of chromosomes 4 and 10^62^. The D4Z4 array consists of a variable number of 3.3-kb repeat units, each harboring a *DUX4* open reading frame (ORF). Functional full-length *DUX4* mRNA is produced only from the distal-most repeat unit, which uses a polyadenylation signal located immediately downstream of the array; the internal *DUX4* ORFs are therefore considered pseudogenes^63,64^. To address this challenge, we determined the origin of the *DUX4* segment involved in the IGH::*DUX4* fusion gene in the UTPT2T-02 sample by combining sequence alignment with per-repeat variant analysis.

First, we annotated a personalized reference genome of the UTPT2T-02 sample with D4Z4 repeat units using D4Z4End2End^65^. The number of D4Z4 repeat units varied across haplotypes and chromosomes (Fig. 7c and Supplementary Fig. 20). The IGH::*DUX4* segment was mapped with high concordance (identity >0.999) to the fourth of 27 D4Z4 repeat units on the maternal chromosome 4 (Fig. 7c and Supplementary Fig. 20). Strikingly, the IGH::*DUX4* segment in this case derived from an internal *DUX4* pseudogene rather than the canonical full-length distal *DUX4* with the polyadenylation signal. This finding indicates that fusion to the IGH locus can transform a normally silent *DUX4* pseudogene into a functional oncogenic driver.

To further validate this finding, we identified sequence variants distinguishing individual D4Z4 repeat units by aligning all D4Z4 units to their consensus sequence (Fig. 7d, Methods). The IGH::*DUX4* segment harbored all variants of the fourth D4Z4 repeat unit on the maternal chromosome 4, a combination unique to that unit in UTPT2T-02. These results indicate that the IGH::*DUX4* segment is derived from this specific repeat unit. In contrast, no D4Z4 repeat unit in the T2T-CHM13 reference aligned to the full-length IGH::*DUX4* segment with >0.995 identity (Supplementary Fig. 21), and no unit harbored all four IGH::*DUX4*-specific variants. This indicates that the source D4Z4 repeat unit of this IGH::*DUX4* fusion is not represented in the standard reference and could not have been identified without the personalized reference.

Finally, to assess the population distribution of this IGH::*DUX4*-associated variant pattern, we matched these IGH::*DUX4*-specific variants against D4Z4 repeat units in 235 phased genome assemblies (Methods). We found that this combination of variants was observed only among East Asian assemblies in this panel (Fig. 7e), suggesting that this IGH::*DUX4*-derived sequence may be more prevalent in East Asian populations. Collectively, these results demonstrate that the genomic structure of the IGH::*DUX4* fusion gene can be more precisely elucidated using personalized reference genomes.

## Discussion

In this study, we developed PRCGAP, to our knowledge, the first comprehensive cancer genome analysis pipeline based on personalized diploid reference genomes. Unlike previous personalized reference-based studies, which have addressed only a single sample or a single class of somatic alteration, PRCGAP integrates haplotype-resolved point mutation, structural variant, copy-number, and methylation analysis within a single unified framework, and we further extended it to pediatric B-ALL clinical samples. PRCGAP recovered the majority of somatic variants identified by standard reference-based approaches while additionally revealing variants in highly repetitive regions, such as centromeres and telomeres, where standard approaches are limited.

The personalized reference genome-based approach enabled several major advances in cancer genome analysis. First, PRCGAP provided a more detailed and integrated view of cancer genome architecture by combining haplotype-resolved copy number, SVs, and point mutations within a single framework. This integration clarified the relationships among well-established cancer-associated mutations, such as the different forms of *TP53* inactivation (LOH, hemizygous LOH, and biallelic inactivation) and *KRAS* mutant allele-specific amplification, at single-haplotype resolution. In clinical pediatric ALL samples, PRCGAP determined the parental origin of chromosomal losses in near-haploid ALL and gains in hyperdiploid ALL. These findings are consistent with prior reports of random parental origin, which argue against a major role for genomic imprinting in these subtypes^51,53,54^.

Second, a substantial fraction of somatic variants were unique to PRCGAP (23.4% of SNVs and 32.6% of SVs), and these were enriched in repetitive regions that are largely inaccessible to standard references. In centromeres, the somatic mutation rate within CDRs, the putative sites of kinetochore formation, was 4.3-fold higher than in the flanking unique regions. PRCGAP also identified centromere-involving SVs accompanied by copy-number and methylation changes as well as inter-haplotype telomere recombination suggestive of break-induced replication. In a pediatric B-ALL sample, PRCGAP identified a 1q gain, a clinically relevant alteration in B-ALL, whose breakpoint fell within a human satellite array and could be resolved only on the personalized reference.

Third, the personalized reference genomes served as a foundation for sequence-aware analyses. We identified novel active L1 source sites absent from standard reference genomes and enabled mechanistic classification of source sites as Solo L1-dominant or TD-dominant, suggesting mechanistic heterogeneity among active L1 source elements. Similarly, structural analysis of the IGH::*DUX4* fusion gene in a pediatric B-ALL sample traced the fused *DUX4* segment to a specific internal *DUX4* pseudogene in a D4Z4 repeat unit whose sequence variant was observed only among East Asian assemblies in the limited population panel. A previous long-read study localized the translocated *DUX4* in the Nalm6 cell line to *DUX4L13* on chromosome 10 using standard reference-based mapping^66^. However, the D4Z4 macrosatellite, which comprises ∼3.3-kb units present on both the 4q35 and 10q26 subtelomeres that share extensive sequence identity, is itself highly variable, differing not only between individuals but also between the two haplotypes within an individual^67^. Standard linear references capture only a single configuration of this array and therefore cannot represent such individual- and haplotype-specific arrangements. Our findings indicate that resolving the source D4Z4 unit at single-unit resolution requires personalized references to identify the true source of the IGH::*DUX4* fusion gene.

Several limitations should be noted. First, PRCGAP requires deep long-read sequencing of both tumor and matched normal samples, typically combining HiFi, ONT-UL, and Hi-C, Pore-C, or trio phasing data. This sequencing requirement poses challenges in terms of sequencing cost, operational complexity, computational resources, and the requirement of substantial high-molecular-weight DNA from fresh-frozen samples for routine clinical application. To mitigate these burdens, the use of pangenome references as an alternative to individual de novo assembly may be a practical strategy^9–12^. In addition, recent advances in ONT-only diploid assembly with hifiasm^68^ or Verkko with read correction^69^ may alleviate the data-type complexity by enabling high-quality assembly from ONT data alone. Second, haplotype switch errors in the assembled reference genomes, although infrequent, may affect the accuracy of haplotype-resolved analyses in the affected regions. Third, genomic regions lacking haplotype-specific markers, such as ribosomal DNA arrays and highly similar segmental duplications, remain difficult to resolve even with personalized references, as evidenced by the residual multi-mapped reads observed there.

Recent perspectives anticipate that complete, diploid-genome-based sequencing, integrating haplotype-resolved assembly, pangenome references, and advanced variant interpretation, will converge into a unified paradigm for clinical genomics^70,71^. Personalized reference genome-based analysis is a natural step toward this goal, yet a comprehensive framework spanning the full spectrum of somatic alterations has been lacking. PRCGAP provides an early example of such a framework that anticipates this shift and can extend personalized reference genome-based cancer genome analysis from cell lines to clinical samples.

## Methods

### Sample collection

The cell lines used in this study (H209, BL209, H2126, BL2126, HCC1954, and HCC1954BL) were obtained from the American Type Culture Collection (ATCC). All cell lines were cultured according to the provider’s instructions. A patient-derived xenograft of NH-ALL-001 was established to obtain a sufficient yield of tumor samples. For the other tumor samples (UTPT2T-02 and UTPT2T-04), cryopreserved mononuclear cells of diagnostic bone marrow samples were used. EBV-transformed B-lymphoblastoid cell lines were established from the patients’ peripheral blood mononuclear cells (PBMCs) by infection with EBV and culture in RPMI-1640 with 10% FBS at 37°C with 5% CO2 and were used as matched normal samples. Parental genomic DNA for pediatric B-ALL samples was obtained from peripheral blood of both parents.

### Publicly available sequence data for tumor-normal cell-line pairs

For H1437, H2009, and HCC1937 cell lines and their matched normal cell lines, we obtained PacBio HiFi, ONT-UL R10, Pore-C, and Illumina short-read sequencing data from the Cancer Standards Long-read Evaluation panel (CASTLE, BioProject ID: PRJNA1086849) and ran PRCGAP^19^.

For COLO829 and COLO829BL, PacBio HiFi, ONT-UL R10, and Illumina short-read sequencing data, along with the de novo assembly of COLO829BL, were obtained from the SMaHT data portal (https://data.smaht.org/data/benchmarking/COLO829 and https://data.smaht.org/data/benchmarking/COLO829#BL)^14,21^.

For HG008, PacBio HiFi, standard ONT R10, and Illumina short-read sequencing data from pancreatic normal tissue, Hi-C data from duodenal tissue, and the normal-sample de novo assembly were obtained from the Genome in a Bottle Consortium^20^.

For HCC1954 and HCC1954BL, Illumina short-read whole-genome sequencing data were additionally obtained from a previous study (BioProject ID: PRJDB10898)^28^.

### Sequencing and basecalling

For ONT ultra-long sequencing, genomic DNA was extracted from 5–6 million cells using the Monarch® HMW DNA Extraction Kit for Tissue (NEB, T3060), and libraries were prepared with the ONT Ultra-Long DNA Sequencing Kit (SQK-ULK001 or SQK-ULK114). Sequencing was performed on PromethION using R9.4.1 (FLO-PRO002) or R10.4.1 (FLO-PRO114M) flow cells, and basecalling was performed in super-accuracy mode with 5mC CpG methylation calling using Guppy v6.4.8 or Dorado v0.7.3/v0.9.1 (https://github.com/nanoporetech/dorado), depending on the sample (Supplementary Table 14).

PacBio HiFi sequencing was performed as previously described^72^. HiFi reads were generated by CCS (v6.3.0, v6.4.0, or v8.0.0 with “--hifi-kinetics”) with consensus accuracy further improved by

DeepConsensus v1.2.0^73^ for HCC1954/HCC1954BL and NH-ALL-001. 5mC CpG methylation was predicted by Primrose (v1.3.0) or Jasmine (v2.2.0; Supplementary Table 14).

Hi-C libraries were generated for normal samples (HCC1954BL, NH-ALL-001, BL209, and BL2126) using the MboI restriction enzyme as previously described^72,74,75^.

For short-read whole-genome sequencing, genomic DNA from H209, BL209, and parents of pediatric B-ALL samples was sequenced on the Illumina NovaSeq6000 platform with a standard 151-bp paired-end read protocol according to the manufacturer’s instructions.

### De novo assemblies of matched normal samples

In PRCGAP, de novo assemblies of matched normal samples were generated using hifiasm and Verkko, with different phasing strategies depending on the available data type.

For Hi-C-phased samples (HCC1954BL, NH-ALL-001, BL209, and BL2126), hifiasm was run with Hi-C reads (“--h1” and “--h2”) and ONT-UL reads (“--ul”) together with PacBio HiFi reads, and Verkko was run with Hi-C reads (“--hic1” and “--hic2”) together with PacBio HiFi (“--hifi”) and ONT (“--nano”) reads.

For trio-phased samples (UTPT2T-02 and UTPT2T-04 matched normals), paternal and maternal k-mer databases were generated from parental short-read data using yak (v0.1, https://github.com/lh3/yak) or merqury^76^ (v1.3). Hifiasm was then run with paternal (−1) and maternal (−2) yak databases together with PacBio HiFi reads, and Verkko was run with the “--hap-kmers” option (paternal and maternal Meryl databases) in trio mode together with PacBio HiFi and ONT reads.

For Pore-C-phased samples (BL1437, BL2009, and HCC1937BL), only Verkko was used because hifiasm does not natively support Pore-C input as phasing data, with Pore-C reads (“--porec”) together with PacBio HiFi and ONT reads.

Sample-specific tool versions are listed in Supplementary Table 15. Then, we removed short contigs less than 100 kb as in the HPRC workflow (https://github.com/human-pangenomics/hpp_production_workflows/blob/master/assembly/wdl/tasks/assembly_clean_up.wdl)^9^.

### Annotation of genome assembly

De novo assemblies were annotated with gene, repeat, segmental duplication, and centromere/satellite information. Gene models were transferred from GRCh38 to each assembly using liftoff (v1.6.3) with the Ensembl v112 GTF file as input, and the resulting GFF3 was converted to GTF using GffRead (v0.12.7)^77^. Repeat elements were identified using RepeatMasker (v4.1.6) with the Dfam 3.8 database. From the RepeatMasker results, two subsets were extracted: simple repeats for downstream filtering, and full-length LINE-1 elements (≥5.8 kb; subfamilies L1HS, L1PA2–L1PA5) for SV insertion classification. Segmental duplications were detected using SEDEF (v1.1-35-g5acd139) after masking alpha satellite, human satellite, and simple repeats (>1,000 bp), and the output was filtered to retain alignments with length ≥1,000 bp, gap-compressed identity ≥90%, SEDEF score ≤50, gap percentage ≤50%, and satellite coverage ≤70%. Centromeric and satellite regions were annotated using modified scripts from the alphaAnnotation pipeline (https://github.com/kmiga/alphaAnnotation/tree/main), which integrates alpha satellite HOR classification (HumAS-HMMER), HSat2/3 identification, rDNA detection, and RepeatMasker-based satellite subtype extraction into a unified centromere/satellite annotation.

### Evaluation of genome assembly

De novo assemblies were evaluated for contiguity, completeness, base-level accuracy, structural correctness, and phasing accuracy. Total length, N50, and maximum contig length were calculated, and telomeric repeats at contig ends were detected using seqtk telo (https://github.com/lh3/seqtk). Contigs were aligned to T2T-CHM13 using MashMap (v3.1.3) with one-to-one mapping, 99% identity, and 100 kb minimum segment length^78–80^. T2T chromosomes were defined as contigs with telomere sequences at both ends and coverage exceeding 95% of the corresponding T2T-CHM13 chromosome, with relaxed thresholds of 90% for chromosome 9 and 60% for chromosome Y as in a previous study^27^. Quality values were calculated from PacBio HiFi reads using yak (v0.1). Read alignments were generated using minimap2 (v2.28)^81,82^ with map-hifi and map-ont presets, and assembly errors were detected using Flagger (v1.1.0) with platform-specific HMM models^9^, NucFlag (v0.3.3)^24,25^, and Inspector (v1.3)^23^, defining errors as regions flagged by at least two pipelines. Gene completeness was assessed using compleasm (v0.2.6) in BUSCO mode with the primates_odb10 database^22^. Phasing accuracy was evaluated using yak trioeval for trio-phased samples and pstools (v0.2.a3) for Hi-C-phased samples^83^, noting that pstools tends to underestimate switch error rates compared to yak trioeval^11^. Centromere regions were evaluated by NucFlag (v0.3.3)^24,25^ for α-satellite HOR regions >500 kb to identify misassembled contigs, defined as those containing any MISJOIN, COLLAPSE, or COLLAPSE_VAR region >1 Mb or ≥4 such regions of 100 bp–1 Mb^72^. A centromere was considered correctly assembled when a single non-misassembled contig spanned the full H1L interval of the corresponding T2T-CHM13 chromosome (CenSat v2.1 annotation).

### Coordinate conversion between personalized reference genomes and standard human reference genomes

To enable bidirectional coordinate conversion between personalized reference genomes and standard human reference genomes (GRCh38 and T2T-CHM13 v2.0), two sets of chain files were generated. For converting personalized-reference-based coordinates to GRCh38/T2T-CHM13-based coordinates, each standard reference was aligned to each haplotype of the personalized reference. Conversely, for converting GRCh38/T2T-CHM13-based coordinates to personalized-reference-based coordinates, each haplotype of the personalized reference was aligned to each standard reference. Alignments were performed with minimap2 (v2.28) with the “asm5” preset, and chain files were generated with transanno (v0.4.1). Coordinate conversion in both directions was performed using coordconv (v0.1.2). Coordinates are classified as liftable when they align to the target reference (Match) and non-liftable when they do not. Non-liftable coordinates have two states: “Gap with anchors”, where flanking coordinates are liftable, and “Unmapped”, where they do not align to any coordinates in the target reference (Fig. 1c).

### Personalized reference genome-based cancer genome analysis pipeline (PRCGAP)

PRCGAP performs haplotype-aware read alignment refinement, point mutation calling, SV calling, copy-number analysis, and methylation profiling from personalized diploid reference genomes and long-read sequencing data (PacBio HiFi and/or ONT).

#### Haplotype-aware read mapping refinement

Long-read sequencing data of both tumor and matched normal samples were mapped to the personalized reference genome using minimap2 (v2.28, asm5 for HiFi, asm10 for ONT). These presets were chosen for two reasons: (i) the personalized reference is derived from the matched normal sample, so sequence divergence between reads and the reference should be low; and (ii) the stricter alignment criteria of asm presets prevent over-extension of alignments across SV breakpoints, preserving sharp breakpoint resolution. Haplotype assignments were refined with bam_refiner (v0.3.6) using haplotype-specific k-mers (k = 21, occurring once in each haplotype). Haplotype-specific k-mers were extracted as follows: all 21-mers were generated from each haplotype assembly using Meryl (v1.4)^76^, and 21-mers occurring exactly once in one haplotype and absent in the other were retained as haplotype-specific markers. For each read, the haplotype-specific k-mer coverage ratio was defined as the fraction of haplotype-specific k-mers in the aligned region of the personalized reference supported by the read.

#### Point mutation calling

Somatic point mutations were called using DeepSomatic (v1.8.0) with the PACBIO model^19^, retaining only PASS-filtered variants. For haplotype assignment, ±100 bp flanking sequences around each variant were extracted from the personalized diploid reference and realigned to themselves using bwa mem^84^ (“-a -k 50 -c 1000000”) to identify corresponding positions between haplotypes. Supporting reads were extracted using samtools^85^ mpileup with “--output-QNAME”, and haplotype assignment was performed by mutation_postprocess (v0.1.3) with a sequence similarity threshold of 0.98. To prevent false-positive detection due to mismapping, point mutations were filtered by the haplotype-specific k-mer coverage ratio of supporting reads against the personalized reference (threshold: 0.8). Each somatic point mutation was phased based on the haplotypes of its supporting reads, which had been assigned during haplotype-aware alignment from haplotype-specific k-mers. For a mutation supported by reads from a single haplotype, the mutation was assigned to that haplotype (haplotype1 or haplotype2). When SNVs were supported by reads from both haplotypes, the mutation was regarded as present on both haplotypes (homozygous). A mutation was labeled “unphased” only when none of its supporting reads were assigned to a haplotype.

#### SV calling

SVs were detected from ONT data using nanomonsv (v0.8.0) with “--qv15” preset. Because position-only reads mapped to the diploid personalized reference can be misassigned to the wrong haplotype, post-processing was applied with nanomonsv_postprocess (v0.2.5): ±100 bp sequences around SV breakpoints were extracted from the personalized reference, followed by all-versus-all global alignment, and SVs with ≥98% sequence similarity were defined as identical. Haplotype information was assigned to each breakpoint based on supporting read haplotypes. To prevent false-positive detection due to mismapping, SVs were filtered by the haplotype-specific k-mer coverage ratio of supporting reads against the personalized reference (threshold: 0.6). Each breakpoint was phased independently using the haplotypes of its supporting reads, which had been assigned during haplotype-aware alignment from haplotype-specific markers. A breakpoint was assigned to a haplotype when its supporting reads consistently pointed to that haplotype and was otherwise left unphased. Because the two breakpoints of an SV were phased independently, an SV could be fully phased, partially phased (only one breakpoint assigned; the “partial” phasability category defined above), or unphased. For insertion-type classification, the nanomonsv insert_classify module was modified to handle diploid reference genomes.

#### Copy-number analysis

T2T-CHM13 was aligned to each haplotype assembly using minimap2 (v2.28, asm5 preset) after masking satellite regions with dna-brnn (v0.1)^86^. A reference table was generated from PAF alignments by filtering for mapping quality ≥20 and sequence match ≥100,000 bp, computing coverage in 1 Mb bins, and merging bins with ≥80% coverage into contiguous segments; for male samples, sex chromosomes were consolidated to the appropriate haplotype. Sequence depth was calculated at each position using samtools depth (v1.17) with “-Q 40” for tumor and matched normal PacBio HiFi data. Tumor/normal depth ratios were computed in 50,000 bp bins, corrected by total sequencing coverage, median depth ratio, and expected ploidy, and smoothed with a three-bin rolling mean.

#### Methylation profiling

CpG methylation was called using pb-CpG-tools (v2.3.2, https://github.com/PacificBiosciences/pb-cpg-tools) pileup mode for HiFi data and modkit pileup (v0.4.1, https://github.com/nanoporetech/modkit) with “--cpg --combine-strands” for ONT data, with primary alignments.

### Variant calling on standard reference genomes

For cross-comparison with PRCGAP, somatic variants were also called using GRCh38 and T2T-CHM13 (v2.0) reference genomes. HiFi and ONT reads were aligned to each reference with minimap2 (v2.28) using the “map-hifi” and “map-ont” presets, respectively. Somatic point mutations were called with DeepSomatic (v1.8.0) using the PACBIO model^19^. Somatic SVs were called with nanomonsv (v0.8.0) using the “--qv25” and “--qv15” presets using HiFi and ONT data, respectively^28^. Germline variants used for downstream filtering were called with DeepVariant (v1.8.0) on normal-sample data^87^.

### Filtering and annotation of point mutations

Point mutations were retained if normal and tumor depths were both ≥10 and VAF was ≥0.05, excluding those overlapping misassembly regions. Germline variants arising from reference differences were removed by filtering mutations matching GRCh38- or T2T-CHM13-based normal-sample DeepVariant variants (v1.8.0)^87^. Mutations mapping to the same position on both haplotypes after liftover were flagged as homozygous. Mutation coordinates were converted to GRCh38 and T2T-CHM13 using coordconv (v0.1.2) and matched against GRCh38- and T2T-CHM13-based DeepSomatic calls. Each mutation was annotated with gene, repeat, centromere/satellite, and segmental duplication information, and polymorphic variants were identified using gnomAD v4.1 merged with ToMMo (54KJPN).

To identify cancer-associated mutations, variants were lifted over to GRCh38 using GATK LiftoverVcf (v4.6.1.0)^88^ and annotated with Ensembl VEP (v105.0)^89^ using Cancer Gene Census (https://cancer.sanger.ac.uk/cosmic, accessed 2024-01-17), PCAWG (obtained from the ICGC Data Portal), PanCanAtlas (v0.2.8), GENIE (v11.1), and SpliceAI databases (https://basespace.illumina.com/analyses/194103939/files). Mutations were flagged as cancerous if they were hotspot mutations (recurrent in PCAWG, PanCanAtlas, or GENIE), truncating mutations in Cancer Gene Census genes, or splicing mutations (splice donor/acceptor variants, or splice region variants with SpliceAI score ≥0.38) in Cancer Gene Census genes.

To benchmark cancer-associated mutation detection, PRCGAP-flagged cancerous mutations were compared against previously reported cancer-associated mutations. For the seven cancer cell lines (H2009, HCC1954, COLO829, H1437, H209, H2126, and HCC1937), DepMap 26Q1 somatic mutations (OmicsSomaticMutations.csv) flagged as Hotspot, LikelyLoF, HessDriver, OncogeneHighImpact, or TumorSuppressorHighImpact in Cancer Gene Census genes were taken as the reference set, with the *TERT* promoter substitution additionally rescued. For HG008, a curated list of pancreatic ductal adenocarcinoma driver mutations (*KRAS* p.G12V, *TP53* p.K132T, and *SMAD4* frameshift) previously reported was used^20^.

### Filtering and annotation of SVs

SVs detected by PRCGAP were retained if normal and tumor read depths were both ≥10 and variant frequency was ≥0.05 (depth thresholds reduced to 5 for COLO829 due to lower coverage of ONT data). SVs overlapping misassembly regions at either breakpoint were excluded. SV breakpoint coordinates were converted to GRCh38 and T2T-CHM13 using coordconv (v0.1.2) and matched against GRCh38-based and T2T-CHM13-based nanomonsv calls (HiFi and ONT) with a 50 bp breakpoint margin. Each SV was annotated with gene, repeat, centromere/satellite, and segmental duplication information.

### Validation of somatic variants

Somatic SNVs and SVs were independently validated by k-mer-based analysis using Illumina whole-genome sequencing data with Meryl (v1.4) to count k-mers generated from tumor Illumina, matched normal Illumina, and normal HiFi (for SNVs) or normal ONT (for SVs) reads (k = 21).

For SNVs, the analysis was restricted to mutations with VAF ≥0.15 to exclude subclonal variants^90^. The most frequent 21-mer centered on the variant position among supporting reads in tumor HiFi data was adopted as the mutant 21-mer, and the corresponding reference 21-mer was extracted from the personalized reference. We retained mutations whose mutant 21-mer was absent from both the personalized reference and normal HiFi counts and excluded those in highly repetitive regions (reference 21-mer count in normal Illumina >10× normal Illumina depth). A mutation was classified as “Supported” when the mutant 21-mer was observed ≥5 times in tumor Illumina and the normal Illumina count did not exceed the expected sequencing-error false-positive count (error rate 0.001).

For SVs, consensus breakpoint sequences were obtained with nanomonsv (“--debug” option). For each SV, two 21-mers centered on each breakpoint position (±10 bp) were extracted from the consensus sequence. SVs overlapping germline variants called by dipcall (v0.3)^91^ from the matched normal assembly against GRCh38 and T2T-CHM13 were removed. An SV was evaluable if both junction 21-mers were absent from the personalized reference and normal ONT 21-mers, and the SV was supported by >4 reads. Evaluable SVs were classified as “Supported” when both junction 21-mers were detected ≥5 times in tumor Illumina and at most once in normal Illumina.

### Kataegis and mutational signature analysis

Kataegis candidates were detected from the chromosomal-arm-stratified VCF files for each haplotype, calculating inter-mutation distances (IMD) between consecutive SNVs per contig, and seeding a cluster wherever six consecutive SNVs had a mean IMD ≤1,000 bp as previously defined^92^; each seed was then extended forwards and backwards one variant at a time while the cumulative mean IMD remained ≤1,000 bp. For each stratified VCF, the SBS96 trinucleotide spectrum was computed and decomposed into COSMIC mutational signatures (v3.4)^38^ using the VCF2SPECTRUM (https://github.com/ryansohny/VCF2SPECTRUM)^14^.

### Somatic SNV rate analysis in centromere regions

Using the evaluation results of centromere regions, we excluded HCC1954 and HG008 owing to the limited number of available centromeres (less than 30 centromeres of 46 chromosomes). For each sample we classified the centromere and its flank of the matched normal assembly into four sequence classes, computed independently for each haplotype. Centromere boundaries were defined from the dna-brnn (v0.1) calls, merging calls within 10 kb into arrays and keeping the arrays that contained an active HOR annotation from the centromere/satellite (Cen/Sat) annotation as the centromere locus. Each locus was extended by 500 kb on both sides to form the flanking region. The Unique class was the flanking sequence carrying no RepeatMasker annotation, representing single copy pericentromeric sequence. The Mon./div. class (monomeric and divergent α-satellite) comprised the Cen/Sat Monomeric, Inactive HOR, and mixed-alpha annotations. The Active HOR class was the Cen/Sat Active HOR annotation. The CDR class was the hypomethylated dip within the Active HOR array, identified by CDR-Finder (https://github.com/logsdon-lab/CDR-Finder) from the tumor Hi-Fi methylation signal. Somatic SNVs passing the filters described above were each assigned to a sequence class by the position of the variant in the assembly. For each class the mutation rate was the number of SNVs divided by the total size of the class in base pairs.

### Identification of telomere-associated SVs

Telomere-associated SVs were identified from both two-breakpoint SVs and single-breakend (SBND) events detected by nanomonsv. Candidate two-breakpoint SVs were extracted if the breakpoint annotation contained telomere repeats (Simple repeat class matching TTAGGG/CCCTAA hexameric permutations), subtelomeric satellites (Satellite/subtelo class), or the inserted sequence contained ≥5 consecutive telomere repeat hexamers. Supporting reads were extracted from the tumor BAM and analyzed with RepeatMasker to identify telomere repeats; SVs were retained if ≥1 supporting read contained ≥60 bp of telomere repeats or if the SV was subtelomeric by breakpoint annotation. Each supporting read was structurally classified by parsing primary and supplementary alignments into Breakpoint1, Breakpoint2, Telomere, or Inserted_seq segments. To remove likely false-positive translocations arising from telomere-to-telomere mismapping, translocations were filtered out when all supporting reads had only a single breakpoint with telomere annotation and the breakpoint overlapped a telomere region.

For SBND events, candidates with consensus sequences containing ≥1,000 bp of RepeatMasker-annotated telomere repeats were selected. Supporting reads were extracted from the tumor BAM (soft-clipped regions within 50 bp of the breakpoint, soft-clip length ≥100 bp). SBNDs were retained if ≥1 supporting read contained ≥60 bp of telomere repeats. Reads were classified into Breakpoint, Telomere, and Soft_clipped segments, and SBNDs were removed as false positives when the breakpoint overlapped a telomere region and all supporting reads contained only Telomere and Breakpoint segments.

### LINE-1 source site detection

To identify source elements of somatic L1HS retrotransposition, we developed a k-mer-based approach matching unique sequences of L1HS elements in the personalized reference to inserted sequences of somatic L1 insertions (k = 21).

L1HS elements were extracted from RepeatMasker annotations, their sequences retrieved using samtools (v1.17) with reverse-strand elements converted using seqtk (v1.4), and all 21-mers extracted using Meryl (v1.4). 21-mers with genome-wide occurrence ≤2 were retained as rare 21-mers distinguishing individual L1HS copies, and their coordinates were extracted using bam_refiner (v0.3.6). Consecutive 21-mers were clustered into non-overlapping groups of up to 21, with a representative 21-mer (shortest homopolymer run) selected per cluster. Representatives with >80% single-base composition were removed. L1HS coordinates were converted to T2T-CHM13 and GRCh38 using coordconv (v0.1.2), and elements mapping to the same T2T-CHM13 locus (start and end within 50 bp) were identified as allelic pairs.

For each somatic solo L1 insertion, the inserted sequence was searched for representative 21-mers (forward or reverse complement) from each L1HS element, and a source element was assigned when the 21-mer coverage rate (the fraction of matched clusters among clusters spanning from the first match to the 3′ end of the element) was ≥0.5. Partnered and orphan L1 transductions were assigned to source elements using the modified nanomonsv insert_classify module described above (see SV calling section), and partnered transductions were reclassified as solo L1 if the alignment boundary was within 30 bp of the source element boundary.

Source elements were excluded from the solo L1 analysis if (1) the 1-kb 5’-end region was not covered by tumor HiFi reads (typically due to somatic deletion), preventing methylation measurement, or (2) the tumor methylation frequency was ≥0.5, as hypomethylation of the L1HS promoter was expected for active retrotransposition.

### Parent-of-origin analysis

Parental origin was assigned to each contig using allele-specific methylation at known imprinted differentially methylated regions (iDMRs), following the approach described previously^55^. A curated catalog of iDMRs with known parent-of-origin methylation status defined in GRCh38 coordinates was converted to personalized reference coordinates using chain files. For each CpG site mapped to both haplotypes, methylation frequencies were extracted from normal-sample HiFi or ONT methylation calls, and informative sites were used to assign each contig to the parent with the higher cumulative evidence score.

### IGH::*DUX4* structure analysis

The structure of the IGH locus on the breakpoint-containing haplotype was reconstructed using IGLoo^93^. Assembly contigs corresponding to the D4Z4 arrays on chromosomes 4q35 and 10q26 were extracted with samtools (v1.17) and annotated with D4Z4End2End^65^ to identify D4Z4 repeat units, *DUX4* ORFs, and the polyadenylation signal on both haplotypes.

Individual D4Z4 units downstream of p13E-11 (the proximal marker of the D4Z4 array) were extracted, with minus-strand sequences reverse-complemented using seqtk (v1.4). Multiple sequence alignment of all D4Z4 units was performed using MAFFT^94^ (v7.526, “--auto --reorder” option). A consensus sequence generated by EMBOSS cons^95^ (v6.6.0.0) was used as a reference, against which all units including IGH::*DUX4* were realigned with minimap2 (v2.28, asm20, “--eqx” option). Per-unit variants were extracted using samtools mpileup, and D4Z4 units sharing the same variant pattern as the IGH::*DUX4* at all variant positions were identified as candidate sources. To further localize the source unit, each D4Z4 array was split into 2-kb windows, the IGH::*DUX4* sequence was aligned against each window using minimap2 (asm20 --eqx), with sequence identity threshold of 0.995.

To assess the population distribution of the IGH::*DUX4*-associated variant pattern, 235 phased genome assemblies from the OpenHGL (https://github.com/lh3/OpenHGL) with unambiguous chromosome assignment and super-population information were analyzed. Chromosome 4 and 10 contigs were extracted using AGC (v3.2.0)^96^, and annotated with D4Z4End2End^65^. Four marker sequences derived from IGH::*DUX4*-specific variants were searched by exact substring matching (forward and reverse complement); an assembly was classified as a carrier when all four markers co-occurred within a single D4Z4 unit downstream of p13E-11. Carrier proportions were calculated per super-population.

## Data availability

Sequencing data for the tumor-normal cancer cell line pairs will be deposited in a public sequence repository under open access prior to publication. Sequencing data for the three pediatric B-ALL clinical samples, including tumor, matched normal, and parental samples, will be deposited in a controlled-access repository in accordance with the informed consent obtained and ethics approval. Accession numbers will be provided upon publication. Previously published data used in this study are accessible from the original publications and repositories cited in the Methods. The per-sample analysis results, comprising haplotype-resolved copy-number profiles, somatic SNVs and indels, SVs, genomic overviews, and assembly annotations, are available at Zenodo (10.5281/zenodo.21231495).

## Code availability

The PRCGAP pipeline is available on GitHub at https://github.com/yos-sk/PRCGAP. Associated tools (bam_refiner, nanomonsv_postprocess, mutation_postprocess, and coordconv) are available in the same GitHub organization.

## Acknowledgements

This work was supported by Japan Society for the Promotion of Science (JSPS) KAKENHI Grant Number JP24K20896 (to Y.Sa.), JP24H00009 (to S.O.), JP24K19223 (to Y.O.), and JP26K21772 (to K.K.); Grants-in-Aid from the Japan Agency for Medical Research and Development (AMED; JP24ama221538 to Y.Sh., JP24tk0124003h0002, 24zf0127009h0003, and 25ama221530h0002 to S.O., and JP24ck0106791, JP25ck0106019, and JP25kk0305028 to Y.O., JP23ck0106876 to M.K.); National Cancer Center Research and Development Funds (2025-A-03 to Y.Sh.); the Japan Science and Technology Agency (JST) FOREST Program (JPMJFR220L to Y.O.); the Japan Science and Technology Agency Moonshot R&D Program (JPMJMS2022 to K.K.). This work was partially supported by Platform Project for Supporting Drug Discovery and Life Science Research (Basis for Supporting Innovative Drug Discovery and Life Science Research (BINDS)) from AMED under Grant Number JP24ama121021. We thank S. Tarumoto, M. Shinagawa, and staff at the Single-Cell Genome Information Analysis Core (SignAC) at WPI-ASHBi, Kyoto University for the long-read sequence analyses. We thank Y. Ito for technical assistance. Some data used in this work are from a NIH Common Fund-supported effort, the SMaHT Network, and were provided by the SMaHT Data Analysis Center (DAC) [1UM1DA058230] on behalf of the SMaHT Network. More information about the SMaHT Network is available online at https://smaht.org/. We acknowledge the GIAB Consortium for providing the HG008 sequencing data. We also acknowledge the CASTLE panel for providing long-read sequencing data for the H1437, H2009, and HCC1937 cell lines (BioProject PRJNA1086849). We used ChatGPT (OpenAI) and Claude Code (Anthropic) in order to assist in language editing, proofreading, and reviewing computational scripts. We used the NIG supercomputer (provided by ROIS National Institute of Genetics) and SHIROKANE supercomputer (provided by Human Genome Center, the Institute of Medical Science, The University of Tokyo).

## Author contributions

Y.Sh. and M.K. designed the study. Y.Sa., with assistance from K.C., A.O., and Y.Sh., designed and implemented the PRCGAP workflow. Y.Sa., with assistance from S.K., M.S., H.S., Y.I., and Y.Sh., performed the downstream cancer genome analyses based on personalized reference genomes. Y.O., Y.Kogu., Y.T., T.T., T.Y., H.M., S.O., and K.K. contributed to cell-line sample preparation and sequencing experiments. Y.Koga collected the near-haploid ALL clinical sample (NH-ALL-001). S.K., A.S.-O., and M.K. collected the pediatric B-ALL clinical samples and established the EBV-transformed lymphoblastoid cell lines used as matched normal controls. T.M., I.K., and J.T. established the patient-derived xenograft. G.N. and H.A. performed ONT-UL sequencing for the clinical samples. Y.Sa. and Y.Sh. wrote the manuscript with input from all authors.

## References

1. International Cancer Genome Consortium et al. International network of cancer genome projects. Nature 464, 993–998 (2010).

2. Cancer Genome Atlas Research Network et al. The Cancer Genome Atlas Pan-Cancer analysis project. Nat. Genet. 45, 1113–1120 (2013).

3. ICGC/TCGA Pan-Cancer Analysis of Whole Genomes Consortium. Pan-cancer analysis of whole genomes. Nature 578, 82–93 (2020).

4. Nakagawa, H. & Fujita, M. Whole genome sequencing analysis for cancer genomics and precision medicine. Cancer Sci. 109, 513–522 (2018).

5. Nurk, S. et al. The complete sequence of a human genome. Science 376, 44–53 (2022).

6. Cheng, H., Concepcion, G. T., Feng, X., Zhang, H. & Li, H. Haplotype-resolved de novo assembly using phased assembly graphs with hifiasm. Nat. Methods 18, 170–175 (2021).

7. Cheng, H., Asri, M., Lucas, J., Koren, S. & Li, H. Scalable telomere-to-telomere assembly for diploid and polyploid genomes with double graph. Nat. Methods 21, 967–970 (2024).

8. Rautiainen, M. et al. Telomere-to-telomere assembly of diploid chromosomes with Verkko. Nat. Biotechnol. 41, 1474–1482 (2023).

9. Liao, W.-W. et al. A draft human pangenome reference. Nature 617, 312–324 (2023).

10. Gao, Y. et al. A pangenome reference of 36 Chinese populations. Nature 619, 112–121 (2023).

11. Nassir, N. et al. A draft UAE-based Arab pangenome reference. Nat. Commun. 16, 6747 (2025).

12. Ghorbani, M. et al. Near-complete Middle Eastern genomes refine autozygosity and enhance disease-causing and population-specific variant discovery. Nat. Genet. 57, 1119–1131 (2025).

13. Xiao, C. et al. Personalized genome assembly for accurate cancer somatic mutation discovery using tumor-normal paired reference samples. Genome Biol. 23, 237 (2022).

14. Sohn, M.-H., et al. A telomere-to-telomere map of somatic mutation burden and functional impact in cancer. bioRxiv (2025) doi:10.1101/2025.10.10.681725.

15. Wagner, J., et al. A complete human pancreatic cancer genome. bioRxiv (2026) doi:10.64898/2026.05.01.722316.

16. Qin, Q., Heinz, J. & Li, H. Improving long-read somatic structural variant calling with pangenome and de novo personal genome assembly. bioRxiv (2025) doi:10.1101/2025.10.28.685154.

17. Zhang, Y., Qu, H., Qin, Q., Li, H. & Park, P. J. Diploid donor-specific assembly enhances somatic structural variant detection in cancer genomes. bioRxiv (2025) doi:10.1101/2025.10.28.685155.

18. Mack, T., et al. Donor-specific assemblies enhance somatic structural variant detection in complex genomic regions. bioRxiv (2026) doi:10.64898/2026.02.20.707061.

19. Park, J. et al. Accurate somatic small variant discovery for multiple sequencing technologies with DeepSomatic. Nat. Biotechnol. (2025) doi:10.1038/s41587-025-02839-x.

20. McDaniel, J. H. et al. Development and extensive sequencing of a broadly-consented Genome in a Bottle matched tumor-normal pair. Sci. Data 12, 1195 (2025).

21. Coorens, T. H. H. et al. The Somatic Mosaicism across Human Tissues Network. Nature 643, 47–59 (2025).

22. Huang, N. & Li, H. compleasm: a faster and more accurate reimplementation of BUSCO. Bioinformatics 39, btad595 (2023).

23. Chen, Y., Zhang, Y., Wang, A. Y., Gao, M. & Chong, Z. Accurate long-read de novo assembly evaluation with Inspector. Genome Biol. 22, 312 (2021).

24. Vollger, M. R. et al. Long-read sequence and assembly of segmental duplications. Nat. Methods 16, 88–94 (2019).

25. Mc Cartney, A. M., et al. Chasing perfection: validation and polishing strategies for telomere-to-telomere genome assemblies. Nat. Methods 19, 687–695 (2022).

26. Vollger, M. R. et al. Segmental duplications and their variation in a complete human genome. Science 376, eabj6965 (2022).

27. Logsdon, G. A. et al. Complex genetic variation in nearly complete human genomes. Nature 644, 430–441 (2025).

28. Shiraishi, Y. et al. Precise characterization of somatic complex structural variations from tumor/control paired long-read sequencing data with nanomonsv. Nucleic Acids Res. 51, e74 (2023).

29. Tarailo-Graovac, M. & Chen, N. Using RepeatMasker to identify repetitive elements in genomic sequences. Curr. Protoc. Bioinformatics Chapter 4, 4.10.1–4.10.14 (2009).

30. Numanagic, I. et al. Fast characterization of segmental duplications in genome assemblies. Bioinformatics 34, i706–i714 (2018).

31. Tsherniak, A. et al. Defining a cancer dependency map. Cell 170, 564–576.e16 (2017).

32. Bairoch, A. The cellosaurus, a cell-line knowledge resource. J. Biomol. Tech. 29, 25–38 (2018).

33. Bielski, C. M. et al. Widespread selection for oncogenic mutant allele imbalance in cancer. Cancer Cell 34, 852–862.e4 (2018).

34. Sasaki, H. et al. Evaluation of Kras gene mutation and copy number gain in non-small cell lung cancer. J. Thorac. Oncol. 6, 15–20 (2011).

35. Mueller, S. et al. Evolutionary routes and KRAS dosage define pancreatic cancer phenotypes. Nature 554, 62–68 (2018).

36. Stephens, P. J. et al. Massive genomic rearrangement acquired in a single catastrophic event during cancer development. Cell 144, 27–40 (2011).

37. Nik-Zainal, S. et al. Mutational processes molding the genomes of 21 breast cancers. Cell 149, 979–993 (2012).

38. Alexandrov, L. B. et al. The repertoire of mutational signatures in human cancer. Nature 578, 94–101 (2020).

39. Altemose, N. et al. Complete genomic and epigenetic maps of human centromeres. Science 376, eabl4178 (2022).

40. Gershman, A. et al. Epigenetic patterns in a complete human genome. Science 376, eabj5089 (2022).

41. Gao, S., et al. A global view of human centromere variation and evolution. bioRxiv (2025) doi:10.64898/2025.12.09.693231.

42. Dilley, R. L. et al. Break-induced telomere synthesis underlies alternative telomere maintenance. Nature 539, 54–58 (2016).

43. Tubio, J. M. C. et al. Mobile DNA in cancer. Extensive transduction of nonrepetitive DNA mediated by L1 retrotransposition in cancer genomes. Science 345, 1251343 (2014).

44. Burns, K. H. Transposable elements in cancer. Nat. Rev. Cancer 17, 415–424 (2017).

45. Rodriguez-Martin, B. et al. Pan-cancer analysis of whole genomes identifies driver rearrangements promoted by LINE-1 retrotransposition. Nat. Genet. 52, 306–319 (2020).

46. Zumalave, S. et al. Concurrent L1 retrotransposition events promote reciprocal translocations in human tumorigenesis. Science 392, eaee4513 (2026).

47. Schloissnig, S. et al. Structural variation in 1,019 diverse humans based on long-read sequencing. Nature 644, 442–452 (2025).

48. Zhang, J. et al. Deregulation of DUX4 and ERG in acute lymphoblastic leukemia. Nat. Genet. 48, 1481–1489 (2016).

49. Brady, S. W. et al. The genomic landscape of pediatric acute lymphoblastic leukemia. Nat. Genet. 54, 1376–1389 (2022).

50. Holmfeldt, L. et al. The genomic landscape of hypodiploid acute lymphoblastic leukemia. Nat. Genet. 45, 242–252 (2013).

51. Safavi, S. & Paulsson, K. Near-haploid and low-hypodiploid acute lymphoblastic leukemia: two distinct subtypes with consistently poor prognosis. Blood 129, 420–423 (2017).

52. Haas, O. A. & Borkhardt, A. Hyperdiploidy: the longest known, most prevalent, and most enigmatic form of acute lymphoblastic leukemia in children. Leukemia 36, 2769–2783 (2022).

53. Paulsson, K. et al. Formation of trisomies and their parental origin in hyperdiploid childhood acute lymphoblastic leukemia. Blood 102, 3010–3015 (2003).

54. Paulsson, K. & Johansson, B. High hyperdiploid childhood acute lymphoblastic leukemia. Genes Chromosomes Cancer 48, 637–660 (2009).

55. Akbari, V. et al. Parent-of-origin detection and chromosome-scale haplotyping using long-read DNA methylation sequencing and Strand-seq. Cell Genom 3, 100233 (2023).

56. Lundin-Ström, K. B. et al. Parental origin of monosomic chromosomes in near-haploid acute lymphoblastic leukemia. Blood Cancer J. 10, 51 (2020).

57. Forero-Castro, M. et al. Genome-wide DNA copy number analysis of acute lymphoblastic leukemia identifies new genetic markers associated with clinical outcome. PLoS One 11, e0148972 (2016).

58. Hanamura, I. et al. Frequent gain of chromosome band 1q21 in plasma-cell dyscrasias detected by fluorescence in situ hybridization: incidence increases from MGUS to relapsed myeloma and is related to prognosis and disease progression following tandem stem-cell transplantation. Blood 108, 1724–1732 (2006).

59. Yasuda, T. et al. Recurrent DUX4 fusions in B cell acute lymphoblastic leukemia of adolescents and young adults. Nat. Genet. 48, 569–574 (2016).

60. Tanaka, Y. et al. Chromatin architecture modulation in B-cell acute lymphoblastic leukemia carrying DUX4 fusions. Blood 134, 1240–1240 (2019).

61. Kato, S. et al. Genome profiling with targeted adaptive sampling long-read sequencing for pediatric leukemia. Blood Cancer J. 14, 145 (2024).

62. Bakewell, J., Moorman, A. V. & Ryan, S. L. DUX4-rearranged B-ALL: deciphering a biological and clinical conundrum. Leukemia 39, 2835–2847 (2025).

63. Lemmers, R. J. L. F. et al. A unifying genetic model for facioscapulohumeral muscular dystrophy. Science 329, 1650–1653 (2010).

64. van der Maarel, S. M., Tawil, R. & Tapscott, S. J. Facioscapulohumeral muscular dystrophy and DUX4: breaking the silence. Trends Mol. Med. 17, 252–258 (2011).

65. Xiao, L. C. et al. Complete genetic and epigenetic architecture of D4Z4 macrosatellites in FSHD, BAMS, and reference cohorts with D4Z4End2End. Genome Res. 36, 827–848 (2026).

66. Tian, L. et al. Long-read sequencing unveils IGH-DUX4 translocation into the silenced IGH allele in B-cell acute lymphoblastic leukemia. Nat. Commun. 10, 2789 (2019).

67. Ranallo-Benavidez, T. R., et al. KaryoScope: rapid, alignment-free sequence annotation for the pangenome era. bioRxiv (2026) doi:10.64898/2026.05.15.725544.

68. Cheng, H. et al. Efficient near-telomere-to-telomere assembly of nanopore simplex reads. Nature (2026) doi:10.1038/s41586-026-10105-6.

69. Stanojević, D., Lin, D., Nurk, S., de Sessions, P. F. & Šikić, M. Telomere-to-telomere assembly using HERRO-corrected Simplex nanopore reads. Nature (2026) doi:10.1038/s41586-026-10563-y.

70. Sabbagh, Q., Gilissen, C., Yntema, H. G., Vissers, L. E. L. M. & Hoischen, A. Near-perfect genome sequencing in medical genetics. Nat. Genet. (2026) doi:10.1038/s41588-026-02645-4.

71. Ashley, E. A. et al. Harmonizing standards and resources for the medical genome. Nature 655, 47–58 (2026).

72. Shiraishi, Y., et al. Rare k-mers reveal centromere haplogroups underlying human diversity and cancer translocations. bioRxiv (2025) doi:10.1101/2025.07.26.666712.

73. Baid, G. et al. DeepConsensus improves the accuracy of sequences with a gap-aware sequence transformer. Nat. Biotechnol. 41, 232–238 (2023).

74. Rao, S. S. P. et al. A 3D map of the human genome at kilobase resolution reveals principles of chromatin looping. Cell 162, 687–688 (2015).

75. Ochi, Y. et al. Combined cohesin-RUNX1 deficiency synergistically perturbs chromatin looping and causes myelodysplastic syndromes. Cancer Discov. 10, 836–853 (2020).

76. Rhie, A., Walenz, B. P., Koren, S. & Phillippy, A. M. Merqury: reference-free quality, completeness, and phasing assessment for genome assemblies. Genome Biol. 21, 245 (2020).

77. Pertea, G. & Pertea, M. GFF utilities: GffRead and GffCompare. F1000Res. 9, 304 (2020).

78. Jain, C., Dilthey, A., Koren, S., Aluru, S. & Phillippy, A. M. A fast approximate algorithm for mapping long reads to large reference databases. J. Comput. Biol. 25, 766–779 (2018).

79. Jain, C., Koren, S., Dilthey, A., Phillippy, A. M. & Aluru, S. A fast adaptive algorithm for computing whole-genome homology maps. Bioinformatics 34, i748–i756 (2018).

80. Kille, B., Garrison, E., Treangen, T. J. & Phillippy, A. M. Minmers are a generalization of minimizers that enable unbiased local Jaccard estimation. Bioinformatics 39, btad512 (2023).

81. Li, H. Minimap2: pairwise alignment for nucleotide sequences. Bioinformatics 34, 3094–3100 (2018).

82. Li, H. New strategies to improve minimap2 alignment accuracy. Bioinformatics 37, 4572–4574 (2021).

83. Garg, S. Towards routine chromosome-scale haplotype-resolved reconstruction in cancer genomics. Nat. Commun. 14, 1358 (2023).

84. Li, H. Aligning sequence reads, clone sequences and assembly contigs with BWA-MEM. arXiv [q-bio.GN] (2013) doi:10.48550/arXiv.1303.3997.

85. Danecek, P. et al. Twelve years of SAMtools and BCFtools. Gigascience 10, giab008 (2021).

86. Li, H. Identifying centromeric satellites with dna-brnn. Bioinformatics 35, 4408–4410 (2019).

87. Poplin, R. et al. A universal SNP and small-indel variant caller using deep neural networks. Nat. Biotechnol. 36, 983–987 (2018).

88. McKenna, A. et al. The Genome Analysis Toolkit: a MapReduce framework for analyzing next-generation DNA sequencing data. Genome Res. 20, 1297–1303 (2010).

89. McLaren, W. et al. The Ensembl Variant Effect Predictor. Genome Biol. 17, 122 (2016).

90. Dentro, S. C. et al. Characterizing genetic intra-tumor heterogeneity across 2,658 human cancer genomes. Cell 184, 2239–2254.e39 (2021).

91. Li, H. et al. A synthetic-diploid benchmark for accurate variant-calling evaluation. Nat. Methods 15, 595–597 (2018).

92. Alexandrov, L. B. et al. Signatures of mutational processes in human cancer. Nature 500, 415–421 (2013).

93. Lin, M.-J., Langmead, B. & Safonova, Y. IGLoo enables comprehensive analysis and assembly of immunoglobulin heavy-chain loci in lymphoblastoid cell lines using PacBio high-fidelity reads. Cell Rep. Methods 5, 101033 (2025).

94. Katoh, K. & Standley, D. M. MAFFT multiple sequence alignment software version 7: improvements in performance and usability. Mol. Biol. Evol. 30, 772–780 (2013).

95. Rice, P., Longden, I. & Bleasby, A. EMBOSS: The European molecular biology open software suite. Trends Genet. 16, 276–277 (2000).

96. Deorowicz, S., Danek, A. & Li, H. AGC: compact representation of assembled genomes with fast queries and updates. Bioinformatics 39, btad097 (2023).

