## Supplementary Figure for "Personalized reference genome-based pipeline reveals comprehensive haplotype-resolved views of cancer genomes"

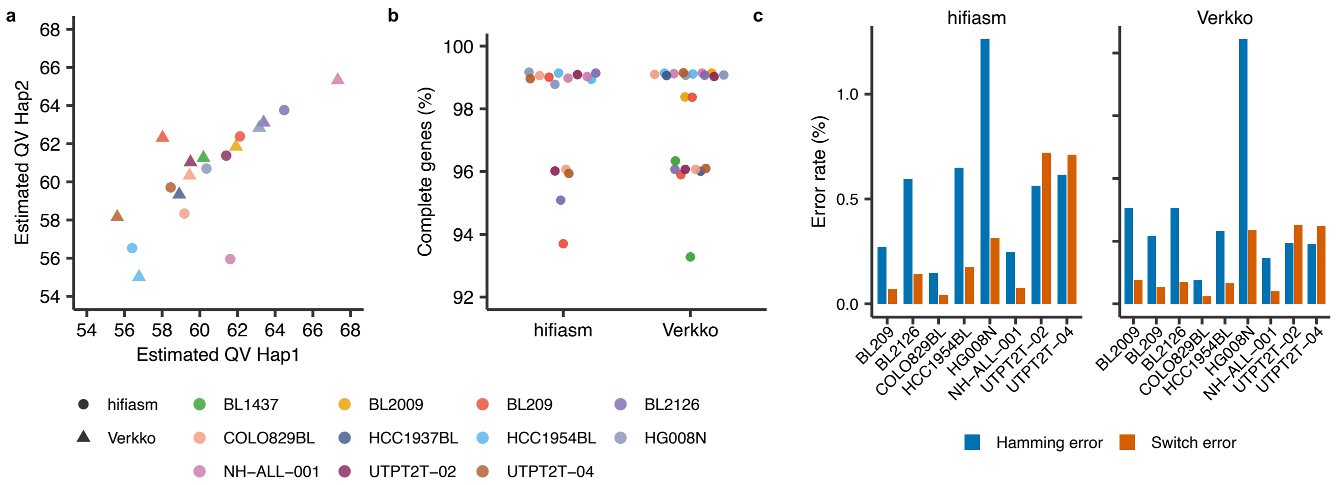


**Supplementary Figure 1. Quality evaluation of personalized reference genome assemblies.** (a) Scatter plot of estimated per-haplotype quality value (QV) for haplotype 1 (x-axis) versus haplotype 2 (y-axis) across all assemblies. Point shapes indicate the assembler (circle, hifiasm; triangle, Verkko) and colors indicate the sample. (b) Percentage of complete genes per assembly, shown separately for hifiasm and Verkko. (c) Haplotype phasing accuracy of each assembly measured by Hamming error (blue) and switch error (orange) rates, shown separately for hifiasm and Verkko.

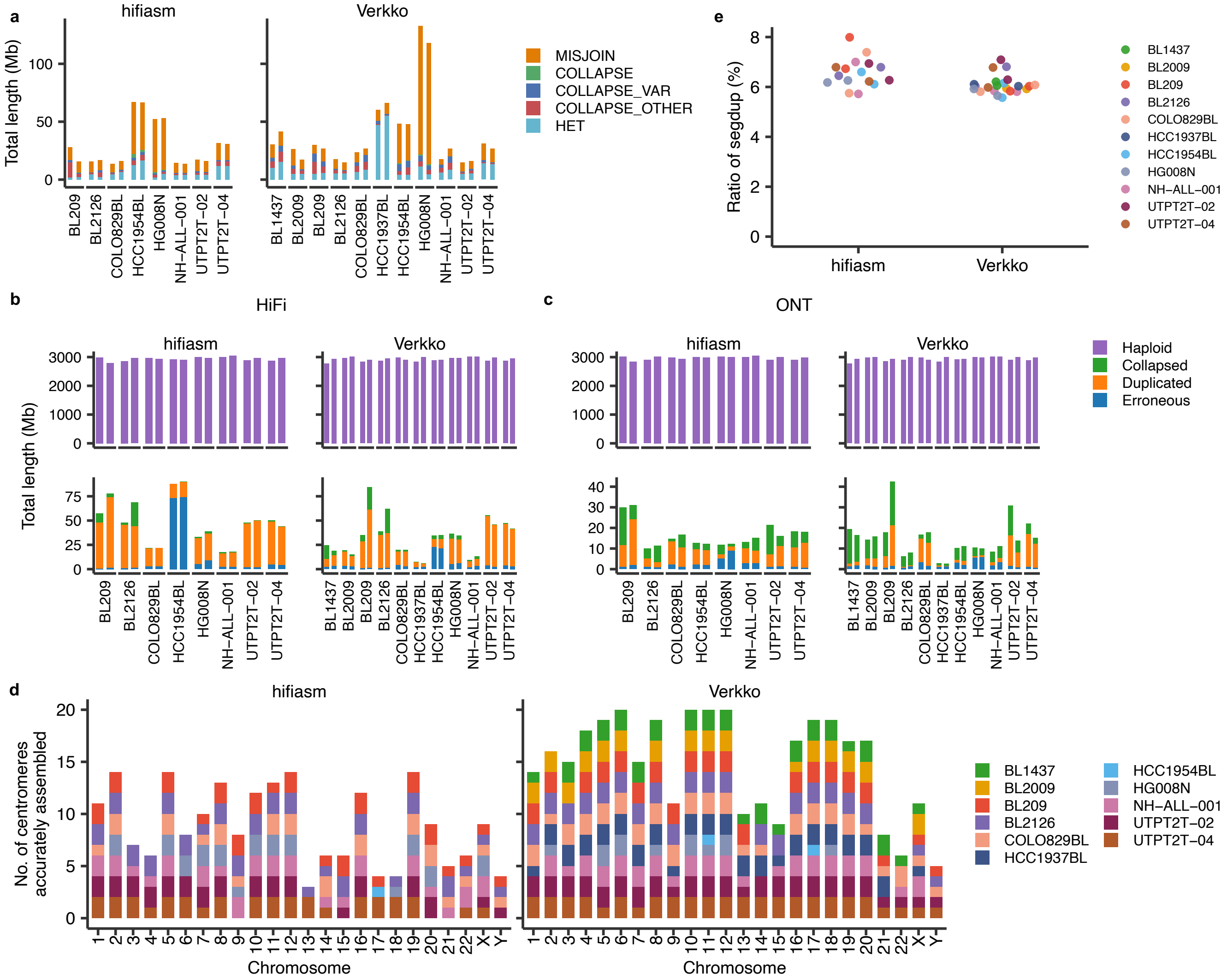


**Supplementary Figure 2. Structural correctness metrics of personalized reference genome assemblies.** (a) Total length of misassembly-flagged regions identified by NucFlag per assembly, stratified by error class (MISJOIN, COLLAPSE, COLLAPSE_VAR, COLLAPSE_OTHER, HET), shown separately for hifiasm and Verkko. (b, c) Total length of assembly regions classified by Flagger into Haploid, Collapsed, Duplicated, and Erroneous categories using HiFi (b) and ONT (c) reads, shown separately for hifiasm and Verkko. (d) Number of accurately assembled centromeres per chromosome identified by NucFlag, shown separately for hifiasm and Verkko. Stacked bars are colored by sample. (e) Ratio of segmental duplication content (segdup) per assembly, shown separately for hifiasm and Verkko.

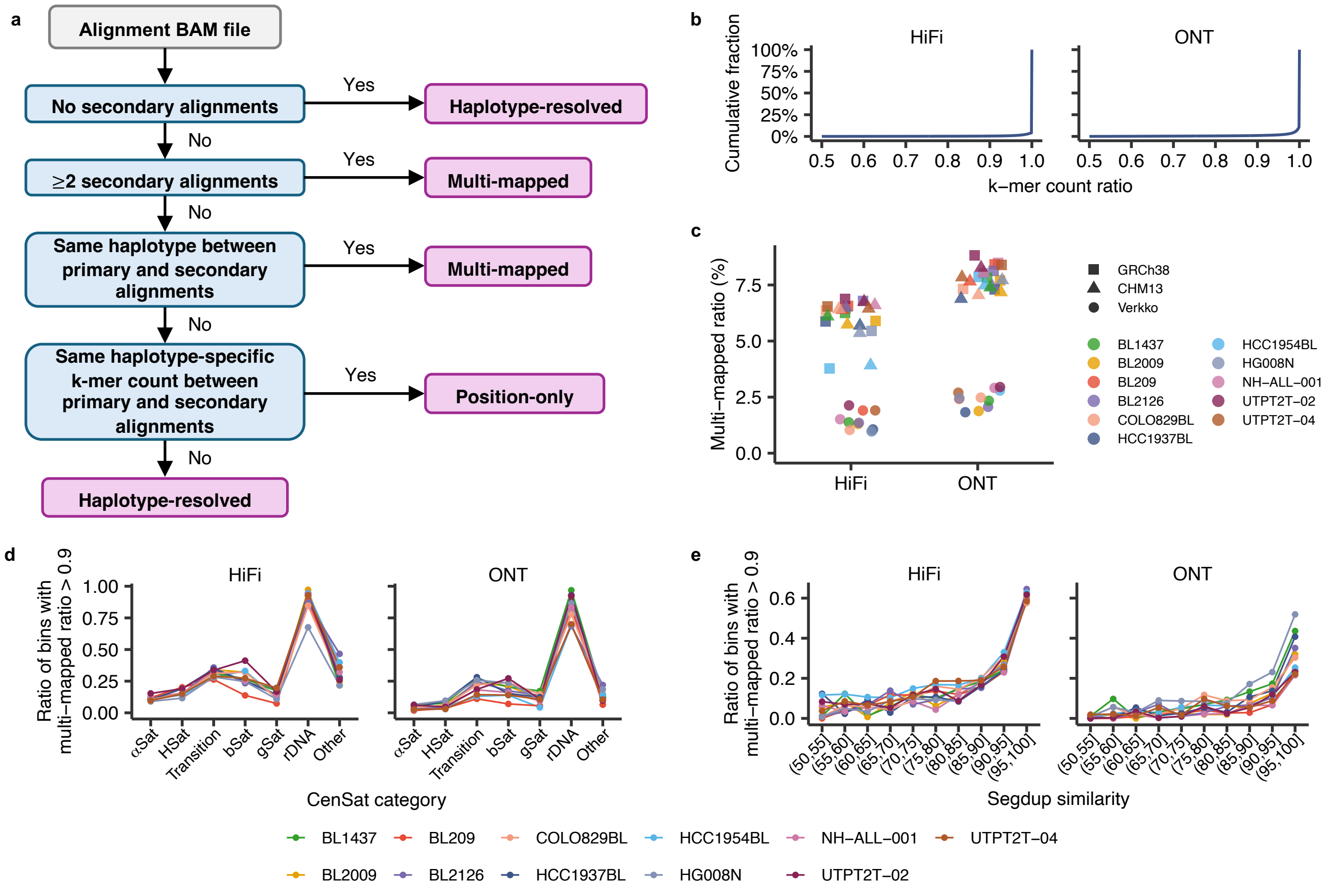


**Supplementary Figure 3. Characterization of multi-mapped reads using haplotype-aware mapping results.** (a) Flowchart of the read classification procedure applied to each alignment. Reads without secondary alignments are classified as Haplotype-resolved. Reads with two or more secondary alignments, or with primary and secondary alignments on the same haplotype, are classified as Multi-mapped. Reads whose primary and secondary alignments have identical counts of haplotype-specific k-mers are classified as Position-only. The remaining reads are classified as Haplotype-resolved. (b) Cumulative distribution of haplotype-specific k-mer ratios for haplotype-resolved reads. The k-mer ratio is defined as the assigned haplotype-specific k-mer count divided by the sum of hap1- and hap2-specific k-mer counts on each read. The sharp peaks at 1 indicate that haplotype-resolved reads were assigned to a haplotype with high confidence. (c) Percentage of multi-mapped reads per sample for HiFi (left) and ONT (right) alignments, shown for alignments against GRCh38 (square), T2T-CHM13 (triangle), and the Verkko-based personalized reference (circle). Colors indicate the sample. (d) Ratio of genomic bins with multi-mapped ratio over 0.9, stratified by Centromere/Satellite category (αSat, HSat, Transition, bSat, gSat, rDNA, and Other), shown for HiFi (left) and ONT (right). (e) Ratio of genomic bins with multi-mapped ratio over 0.9 as a function of segmental duplication similarity (binned from 50% to 100%), shown for HiFi (left) and ONT (right). Lines are colored by sample.

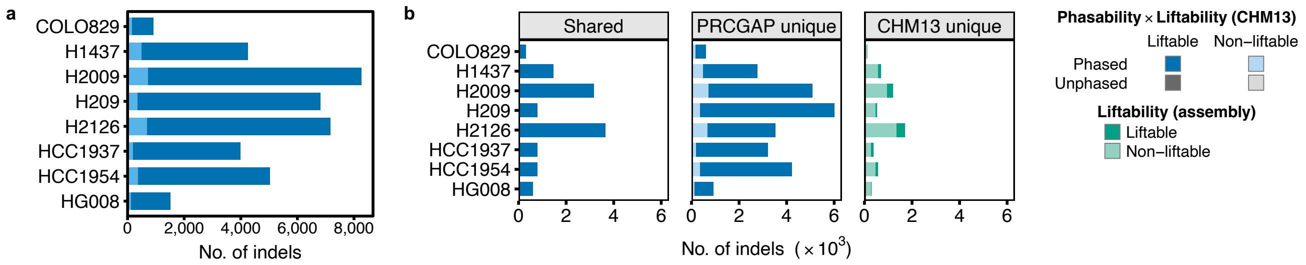


**Supplementary Figure 4. Somatic indel calling by PRCGAP in comparison with the T2T-CHM13-based pipeline.** (a) Total number of somatic indels detected by PRCGAP per cell line. Phasability and liftability (T2T-CHM13) composition is shown by color. (b) Comparison of somatic indels between PRCGAP and the T2T-CHM13-based pipeline, classified into Shared, PRCGAP unique, and CHM13 unique. In the Shared and PRCGAP unique panels, indels are colored by phasability × liftability (T2T-CHM13); in the CHM13 unique panel, by liftability to the personalized reference assembly.

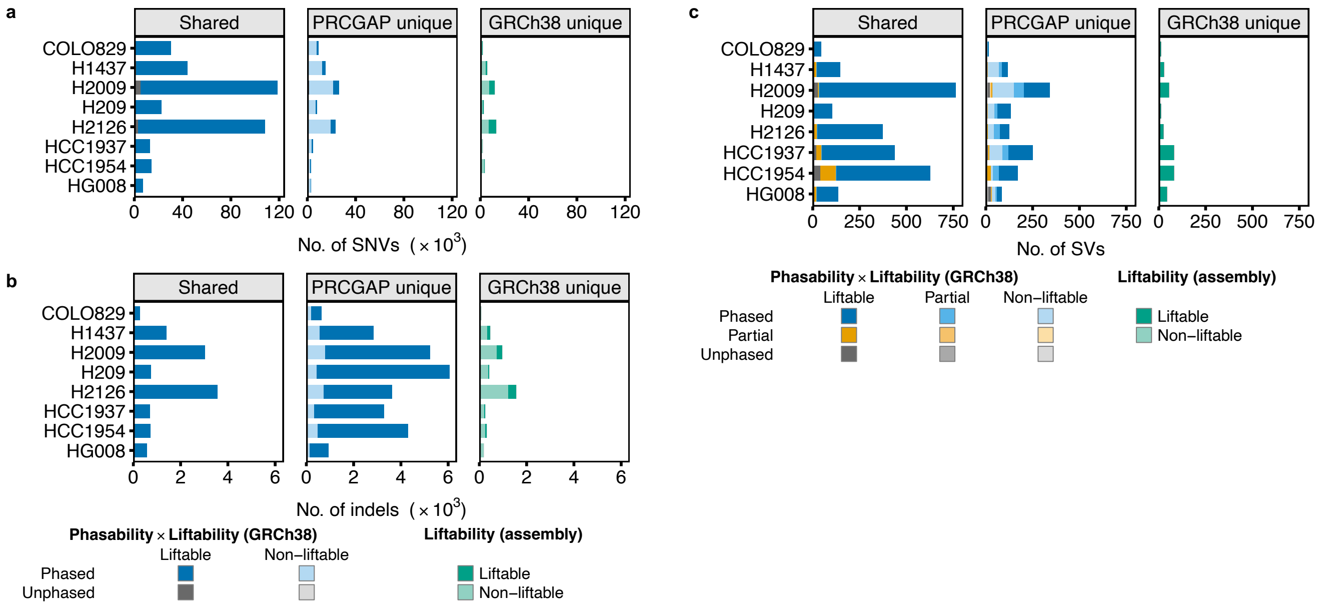


**Supplementary Figure 5. Somatic SNV, indel, and SV calling by PRCGAP in comparison with the GRCh38-based pipeline.** (a) Comparison of somatic SNVs between PRCGAP and the GRCh38-based pipeline, classified into Shared, PRCGAP unique, and GRCh38 unique. In the Shared and PRCGAP unique panels, SNVs are colored by phasability × liftability (GRCh38); in the GRCh38 unique panel, by liftability to the personalized reference assembly. (b) Comparison of somatic indels between PRCGAP and the GRCh38-based pipeline, shown as in a. (c) Comparison of somatic SVs between PRCGAP and the GRCh38-based pipeline, colored by phasability × liftability (GRCh38) including the "Partial" category indicating that only one of the two breakpoints is phased (for phasability) or liftable (for liftability).

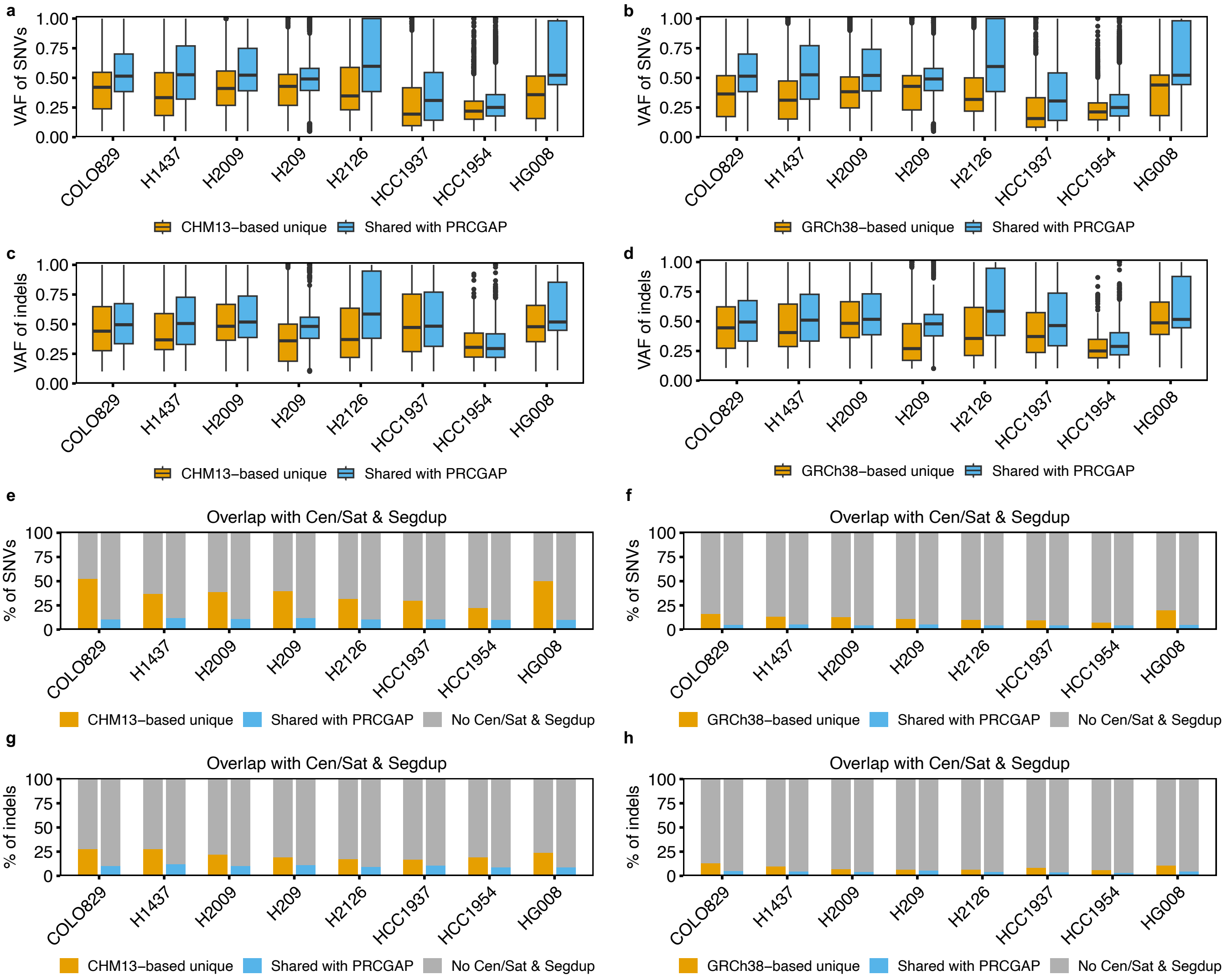


**Supplementary Figure 6. Variant allele frequency and repeat-region distribution of PRCGAP-unique versus standard-reference-unique somatic SNVs and indels.** (a, b) Box plot of variant allele frequency (VAF) of somatic SNVs detected only by the T2T-CHM13-based (a) or GRCh38-based pipeline (b; CHM13/GRCh38-based unique, orange) versus SNVs shared with PRCGAP (Shared with PRCGAP, blue) for each cell line. (c, d) Box plot of variant allele frequency (VAF) of somatic indels detected only by the T2T-CHM13-based (c) or GRCh38-based pipeline (d; CHM13/GRCh38-based unique, orange) versus indels shared with PRCGAP (Shared with PRCGAP, blue) for each cell line. (e, f) Percentage of somatic SNVs located in satellite or segmental duplication regions for T2T-CHM13-based (e) or GRCh38-based (f) unique SNVs versus SNVs shared with PRCGAP. (g, h) Percentage of somatic indels in centromere/satellite (Cen/Sat) or segmental duplication (Segdup) regions for the T2T-CHM13-based (g) or GRCh38-based (h) unique indels versus indels shared with PRCGAP.

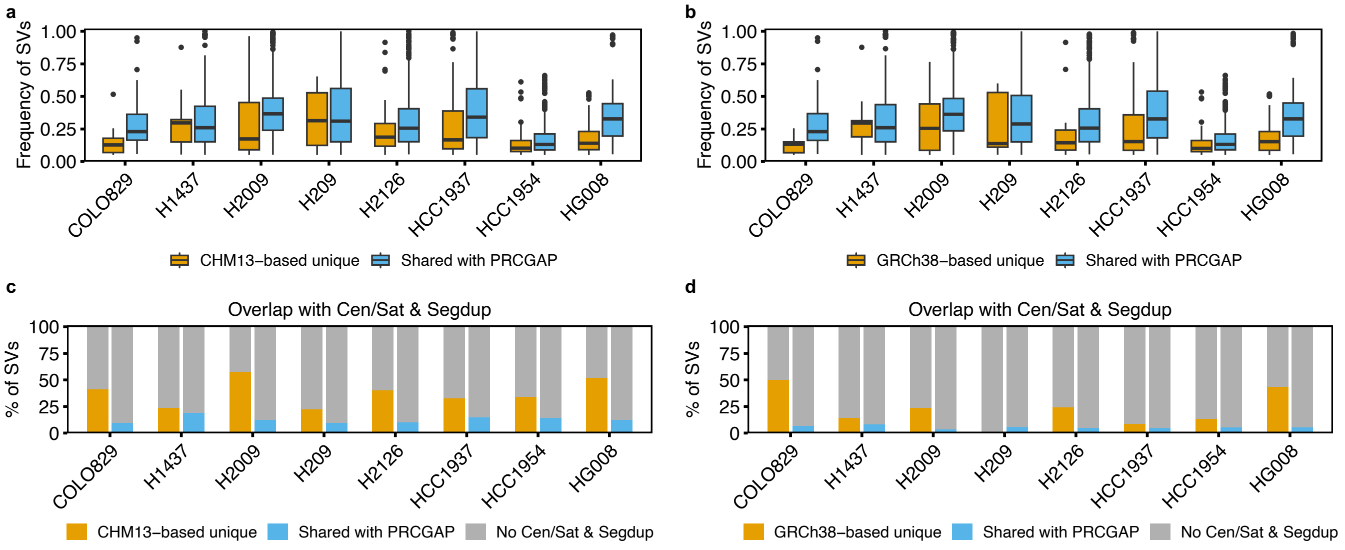


**Supplementary Figure 7. Variant frequency and repeat-region distribution of standard-reference-unique somatic SVs.** (a, b) Box plot of variant frequency of somatic SVs detected only by the T2T-CHM13-based (a) or GRCh38-based pipeline (b; CHM13/GRCh38-based unique, orange) versus SVs shared with PRCGAP (Shared with PRCGAP, blue) for each cell line. (c, d) Percentage of somatic SVs located in centromere/satellite (Cen/Sat) or segmental duplication (Segdup) regions for the SVs detected only by the T2T-CHM13-based (c) or GRCh38-based (d) pipeline versus SVs shared with PRCGAP.

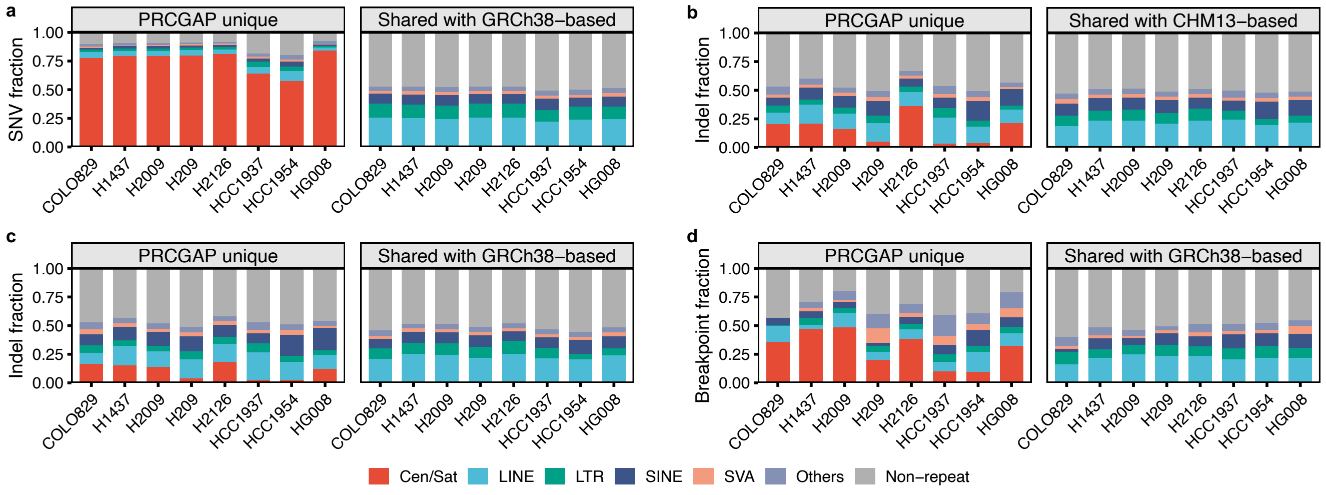


**Supplementary Figure 8. RepeatMasker annotation of PRCGAP-unique versus shared somatic mutations.** (a) Bar plot shows the fraction of somatic SNVs annotated by RepeatMasker, compared between PRCGAP-unique SNVs (left) and SNVs shared with the GRCh38-based pipeline (right). Repeat classes are colored as Cen/Sat, LINE, LTR, SINE, SVA, Others, and Non-repeat. (b) Fraction of somatic indels annotated by RepeatMasker, comparing PRCGAP-unique indels with indels shared with the T2T-CHM13-based pipeline, shown as in a. (c) Fraction of somatic indels annotated by RepeatMasker, comparing PRCGAP-unique indels with indels shared with the GRCh38-based pipeline, shown as in a. (d) Fraction of somatic SV breakpoints annotated by RepeatMasker, comparing PRCGAP-unique breakpoints with breakpoints shared with the GRCh38-based pipeline, shown as in a.

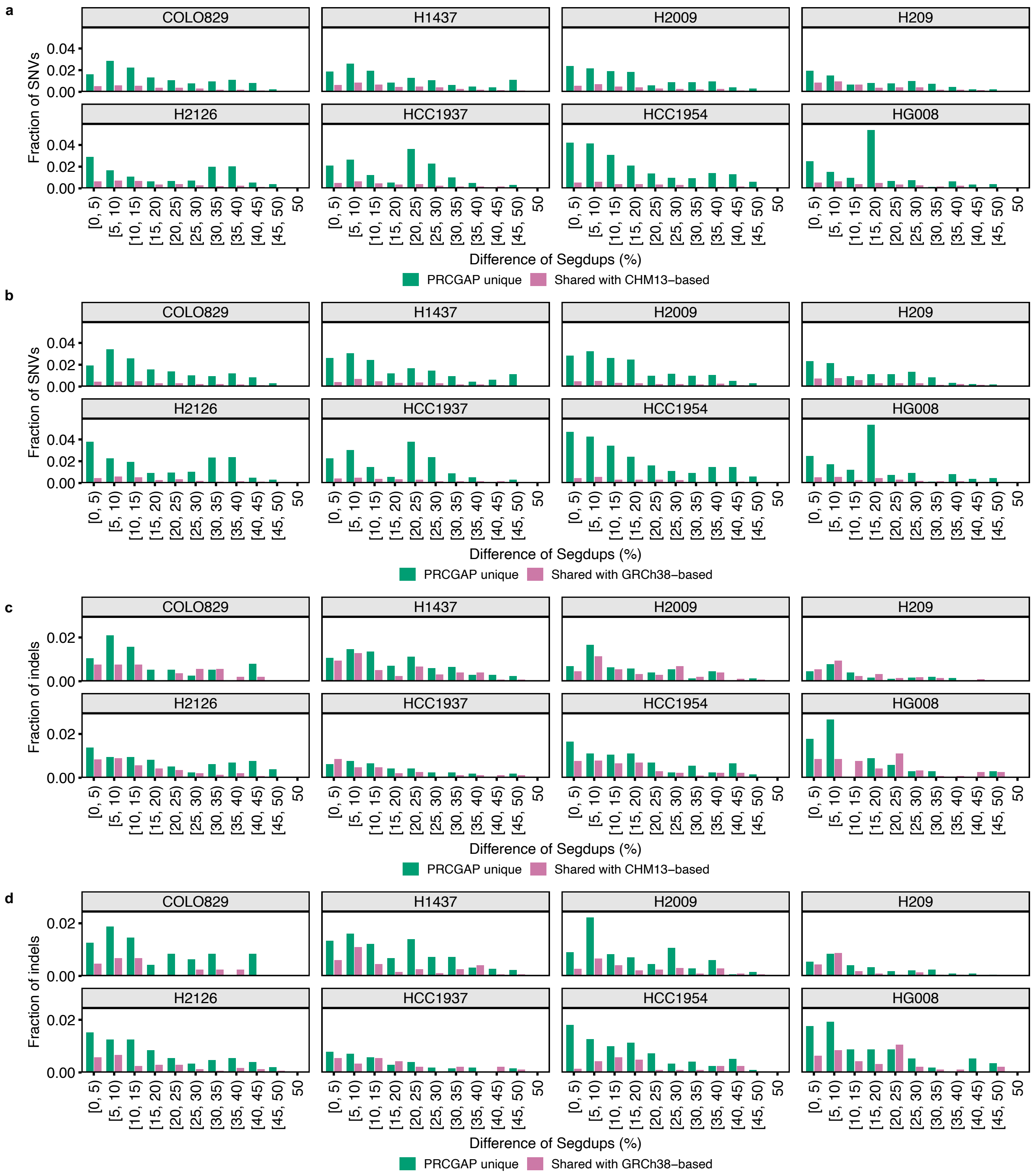


**Supplementary Figure 9. Distribution of PRCGAP-unique and shared somatic mutations relative to segmental duplication divergence.** (a, b) Fraction of somatic SNVs by the percent difference of segmental duplications (binned from [0, 5) to 50), shown per cell line and compared between PRCGAP-unique SNVs (green) and SNVs shared with the T2T-CHM13-based (a) or GRCh38-based (b) pipeline (pink). (c, d) Fraction of somatic indels by the percent difference of segmental duplications (binned from [0, 5) to 50), shown per cell line and compared between PRCGAP-unique indels (green) and indels shared with the T2T-CHM13-based (c) or GRCh38-based (d) pipeline (pink).

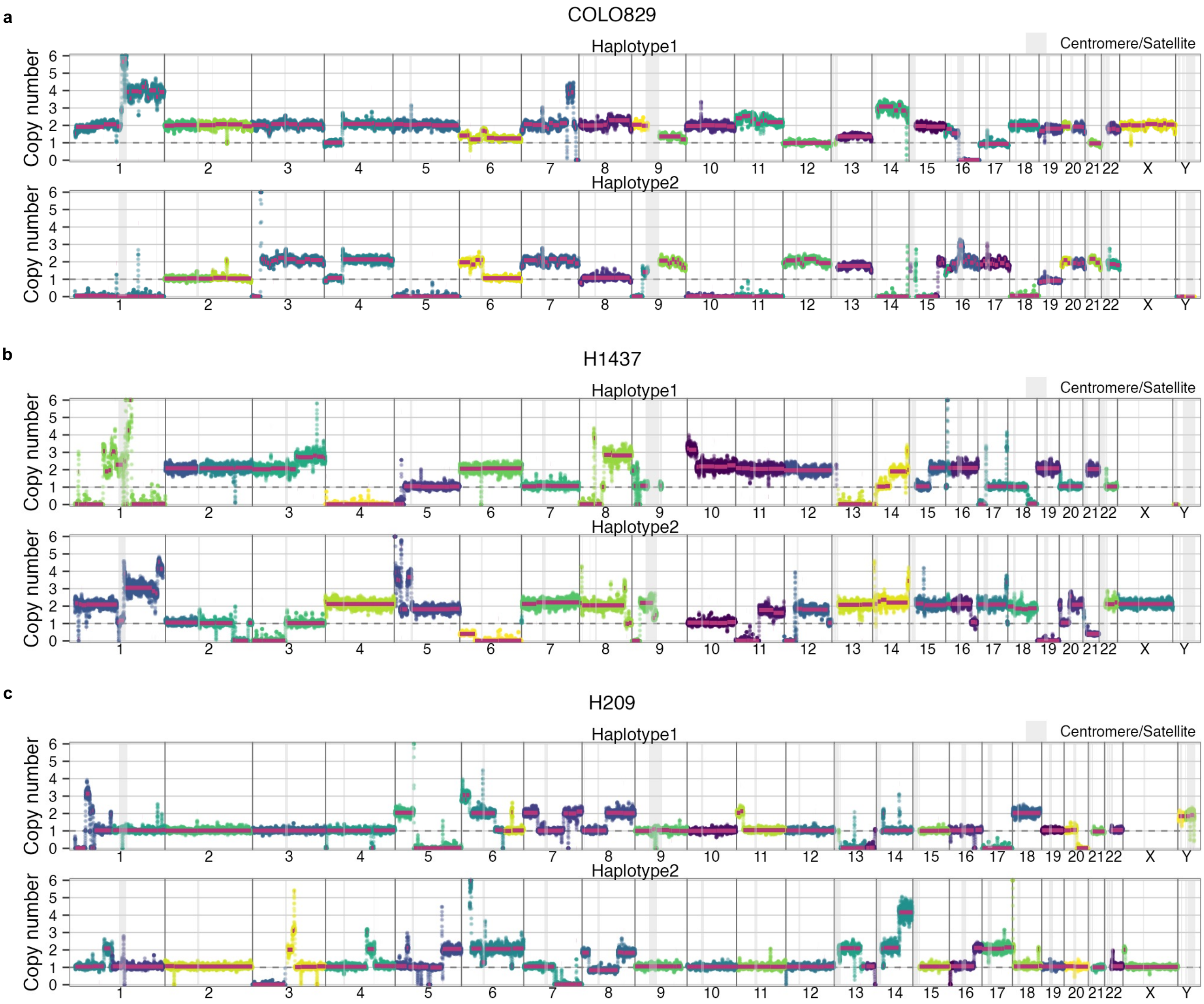


**Supplementary Figure 10. Haplotype-resolved copy-number profiles of cancer cell lines.** (a–g) Haplotype-resolved copy-number profiles of the COLO829 (a), H1437 (b), H209 (c), H2126 (d), HCC1937 (e), HCC1954 (f), and HG008 (g) cell lines. For each sample, haplotype 1 (top) and haplotype 2 (bottom) are shown separately. The y-axis indicates the estimated copy number, and the x-axis represents genomic positions ordered by chromosome. Points are colored by assembled contigs, and centromere/satellite regions are shaded in gray.

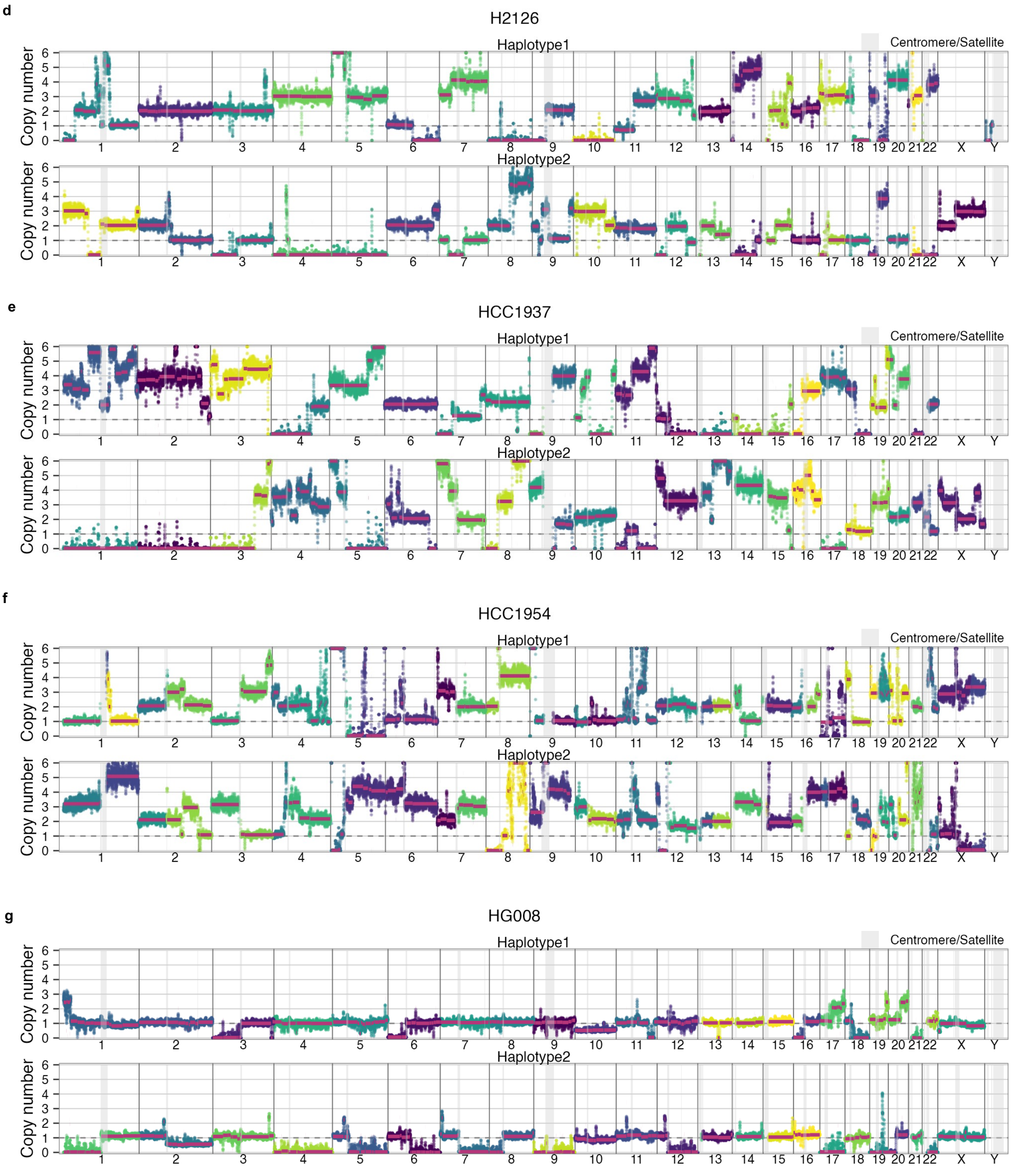


Supplementary Figure 10 (continued)

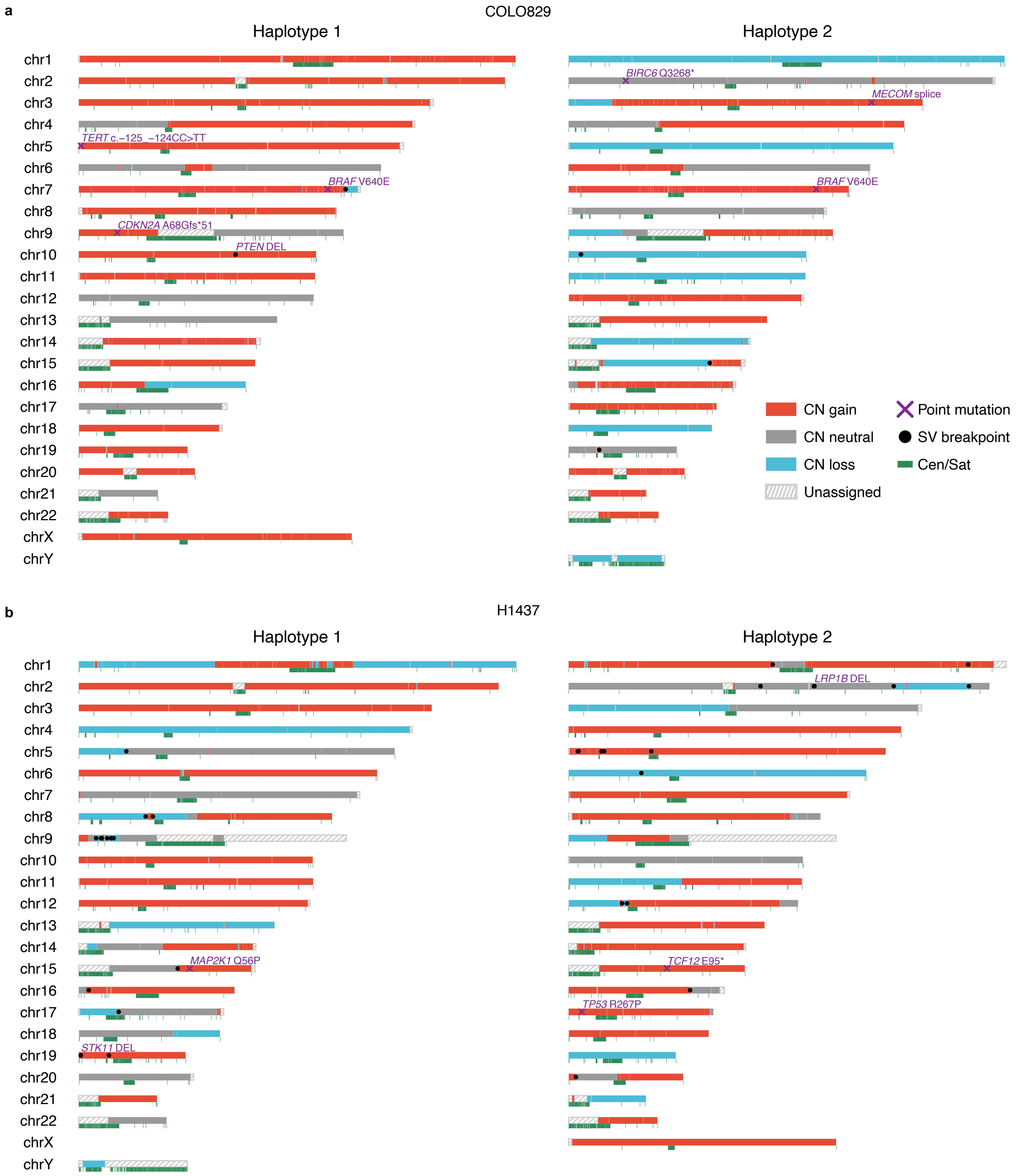


**Supplementary Figure 11. Haplotype-resolved genomic overview of cancer cell lines integrating copy-number status, SV breakpoints, and cancer-associated point mutations.** (a–g) Haplotype-resolved genomic overviews of the COLO829 (a), H1437 (b), H209 (c), H2126 (d), HCC1937 (e), HCC1954 (f), and HG008 (g) cell lines. Each horizontal track represents a chromosome, shown separately for haplotype 1 (left) and haplotype 2 (right). Chromosome segments are colored by copy-number status (CN gain, red; CN neutral, gray; CN loss, light blue; Unassigned, hatched gray). Centromere/satellite regions (Cen/Sat) are marked in green. SV breakpoints are shown as black dots, and cancer-associated somatic point mutations are shown as purple crosses and labeled with the affected gene name.

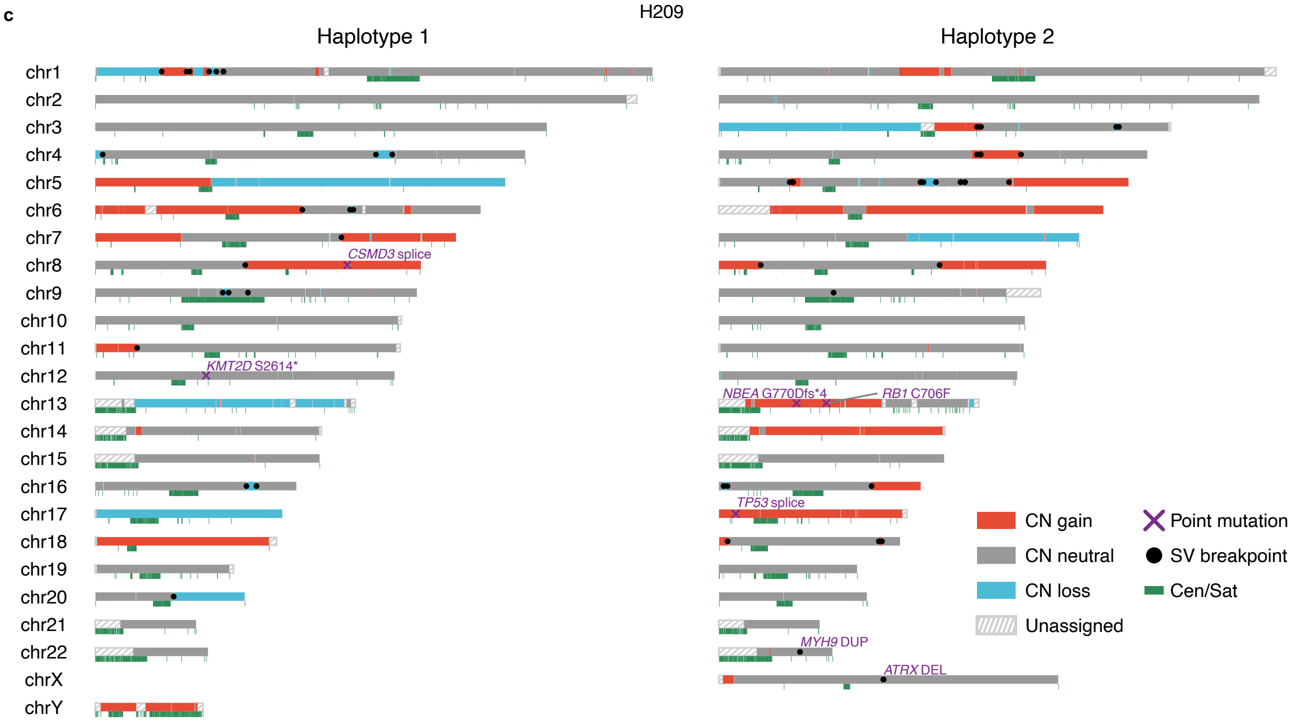


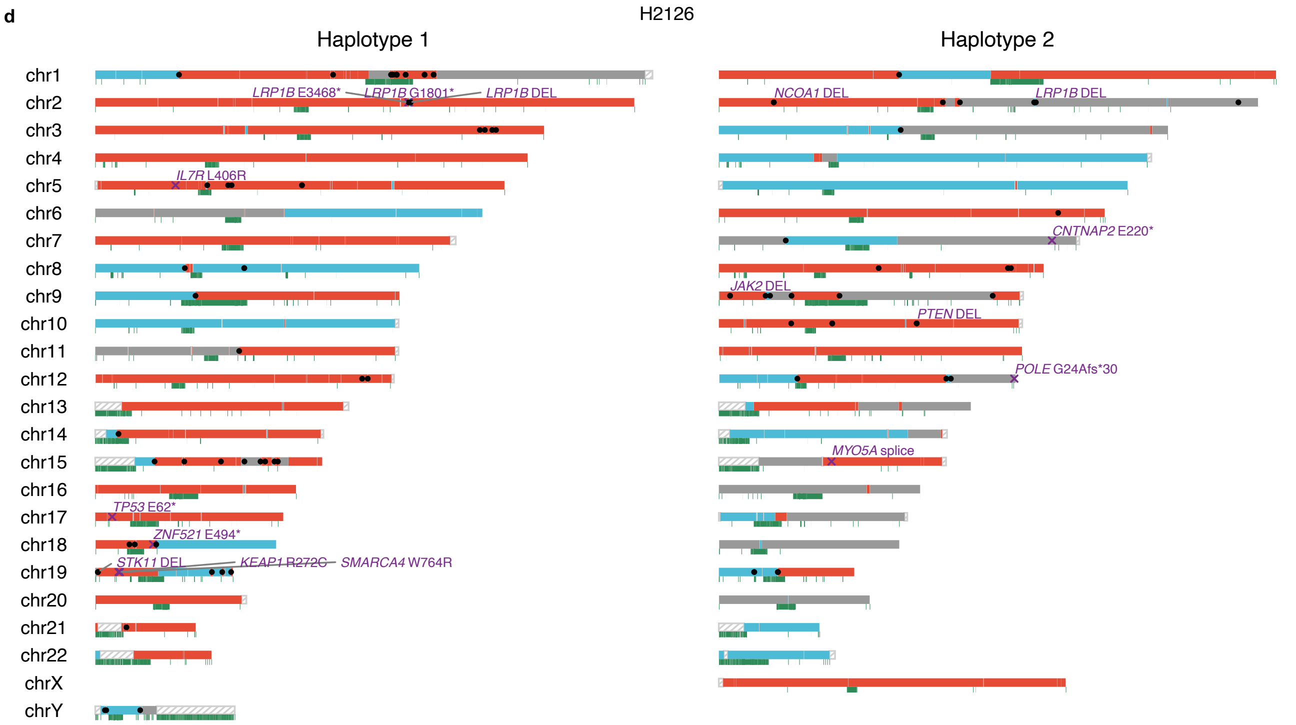


Supplementary Figure 11 (continued)

e

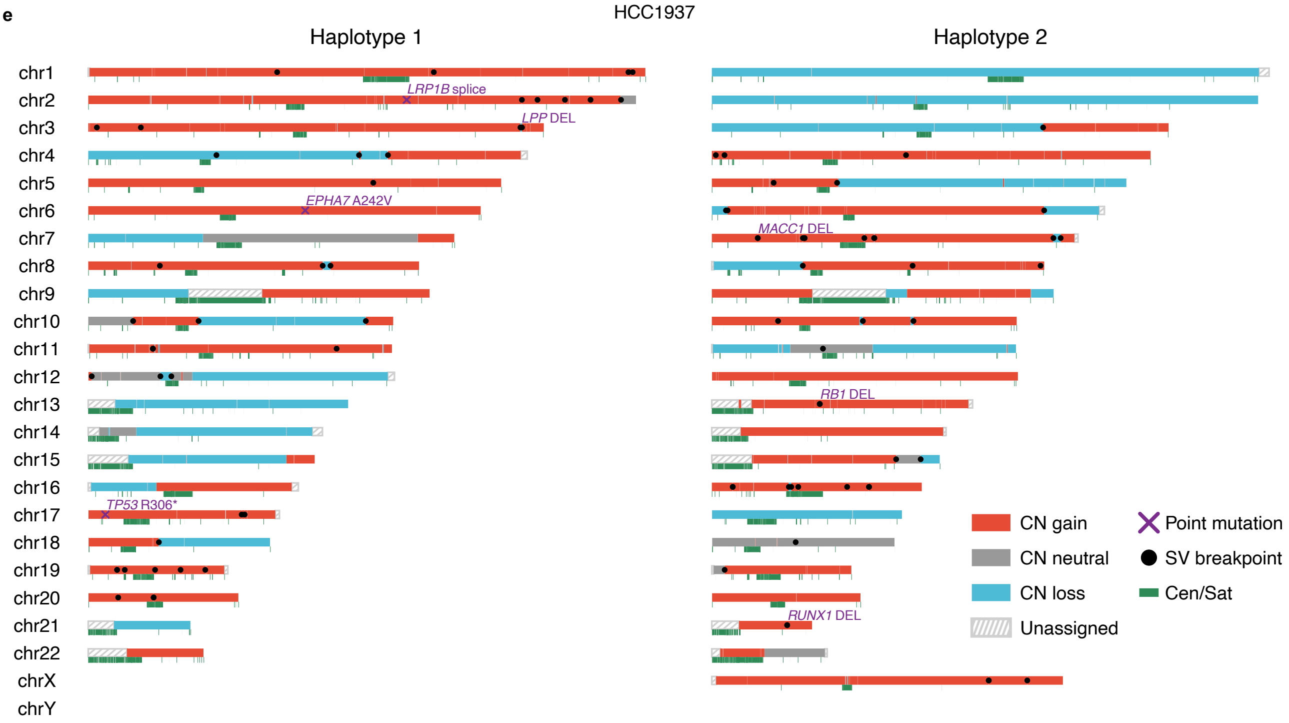


f

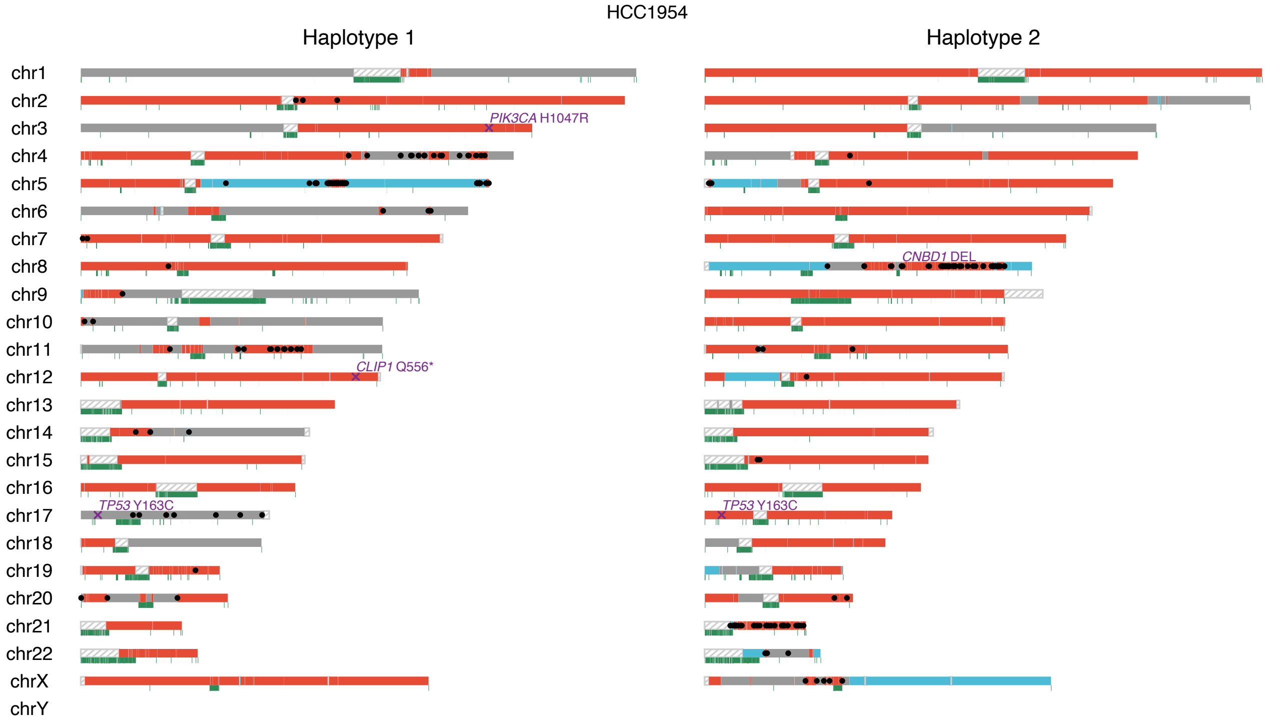


Supplementary Figure 11 (continued)

g

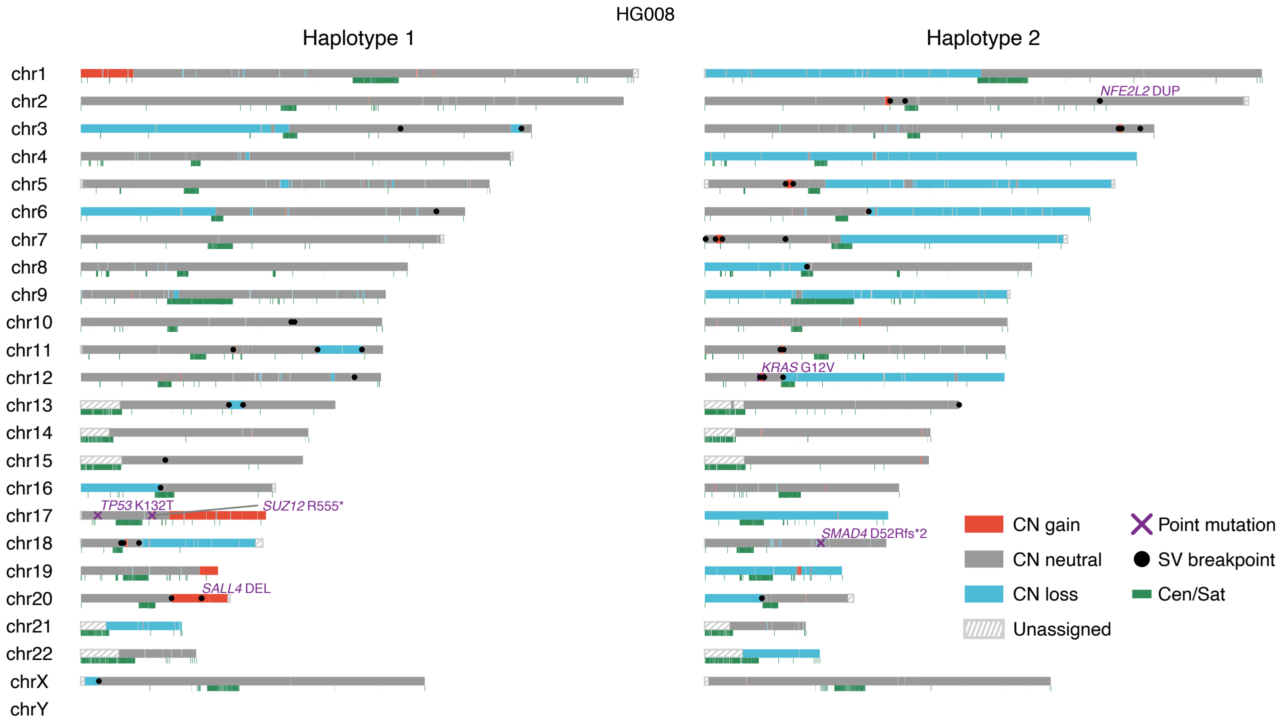


Supplementary Figure 11 (continued)

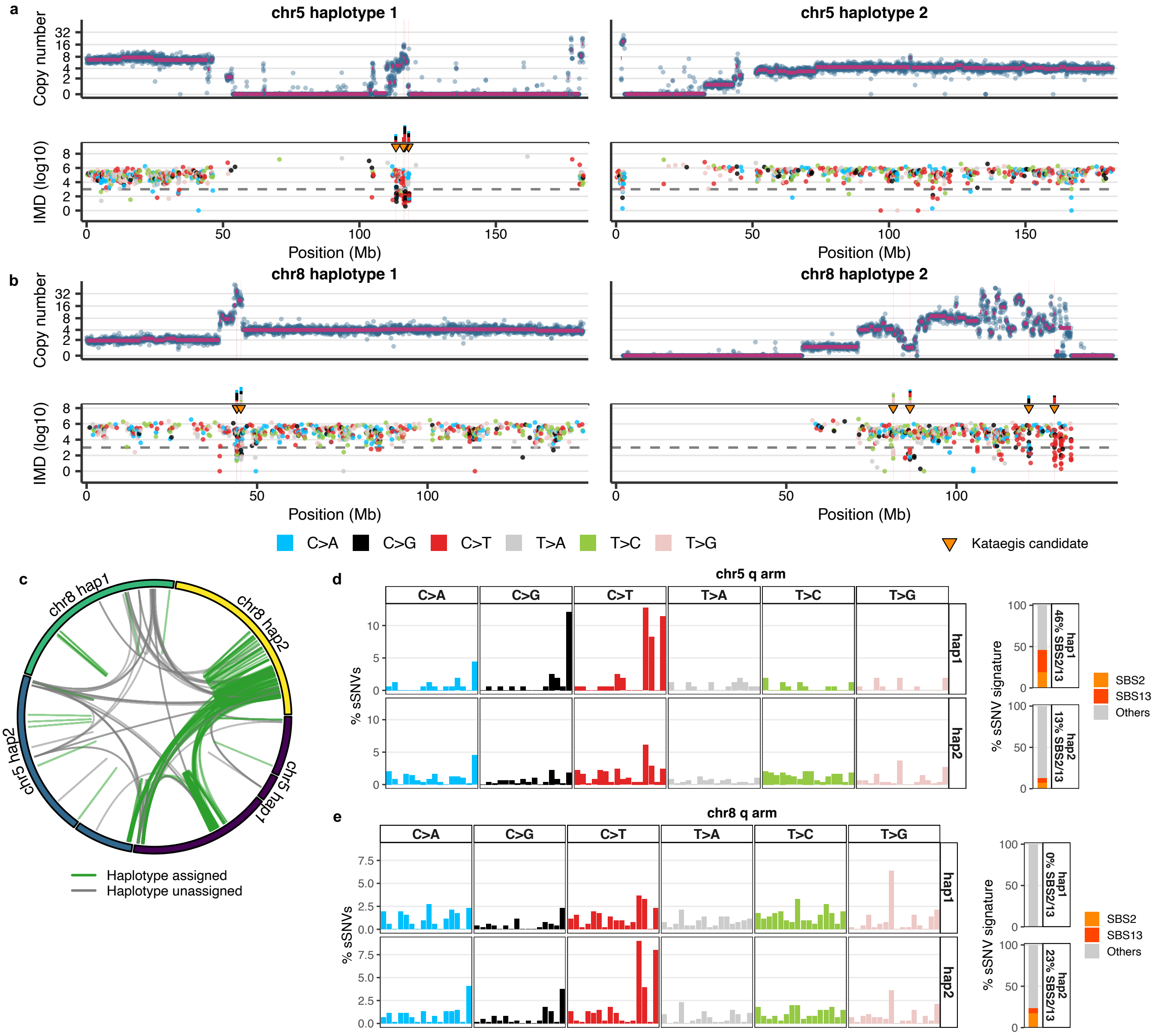


**Supplementary Figure 12. Haplotype-resolved copy-number, kataegis, and mutational signature analysis of chromothripsis-like events on chromosomes 5 and 8 in the HCC1954 cell line.** (a, b) Haplotype-resolved view of chromosome 5 (a) and chromosome 8 (b) in HCC1954 (haplotype 1, left; haplotype 2, right). From top to bottom: copy-number profile, inter-mutation distance (IMD; log10) of somatic SNVs colored by substitution class, and the trinucleotide context of kataegis-candidate SNVs (IMD below the dashed threshold). Kataegis candidate clusters are marked (orange triangles). (c) Circos plot of SVs involving chromosomes 5 and 8 in HCC1954. Haplotype-assigned SVs are shown in green and haplotype-unassigned SVs in gray. (d, e) Trinucleotide mutational spectrum of somatic SNVs on the chromosome 5 q arm (d) and chromosome 8 q arm (e) in HCC1954, shown separately for haplotype 1 (top) and haplotype 2 (bottom). The stacked bars on the right show SBS signature contributions per haplotype (SBS2, SBS13, and Others), with the combined SBS2/13 percentage indicated for each haplotype.

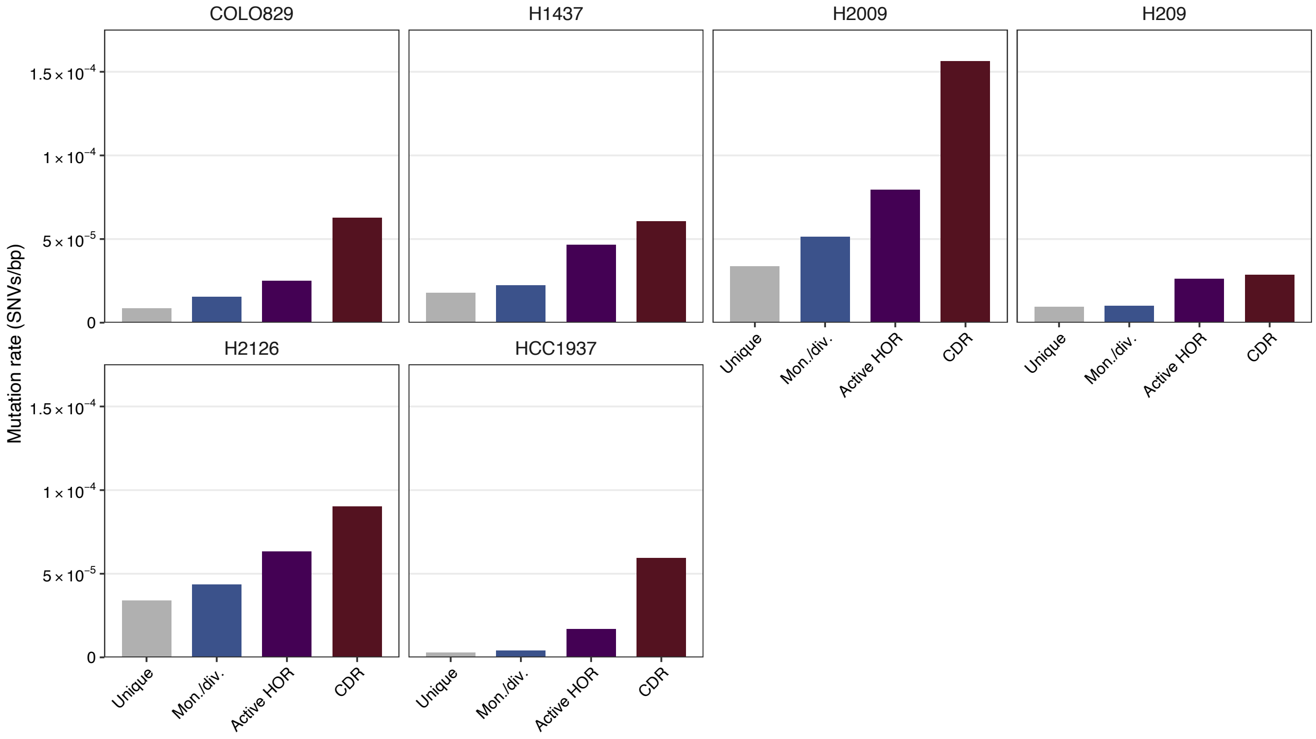


**Supplementary Figure 13. Somatic SNV rate of cancer cell lines in centromeric regions.** Somatic SNV rate (SNVs/bp) of COLO829, H1437, H2009, H209, H2126, and HCC1937 cell lines, stratified by centromere annotation class (Unique, Mon./div., Active HOR, and CDR). HCC1954 and HG008 were excluded owing to the limited number of available centromeres (Methods).

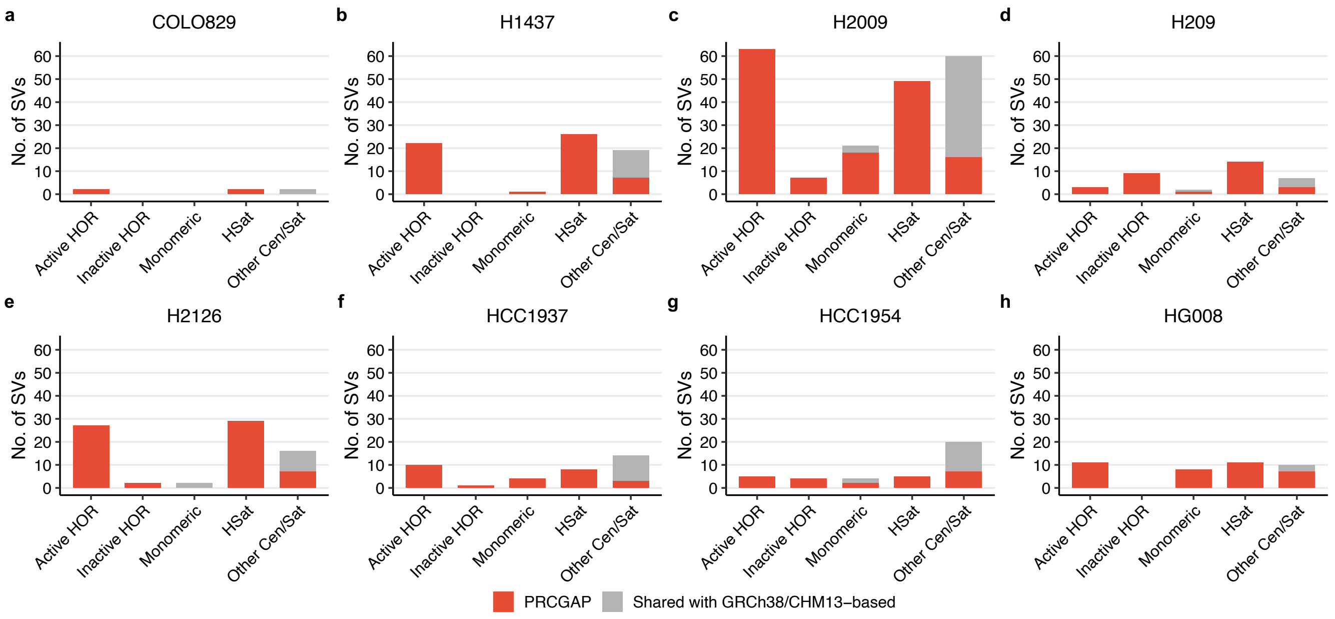


**Supplementary Figure 14. Centromere-associated somatic SVs across cancer cell lines.** (a–h) Number of somatic SVs overlapping centromere-annotated regions in the COLO829 (a), H1437 (b), H2009 (c), H209 (d), H2126 (e), HCC1937 (f), HCC1954 (g), and HG008 (h) cell lines, stratified by centromere annotation class (Active HOR, Inactive HOR, Monomeric, HSat, and Other Cen/Sat). Bars are colored by whether the SV was unique to PRCGAP (red) or shared with the GRCh38/T2T-CHM13-based pipelines (gray).

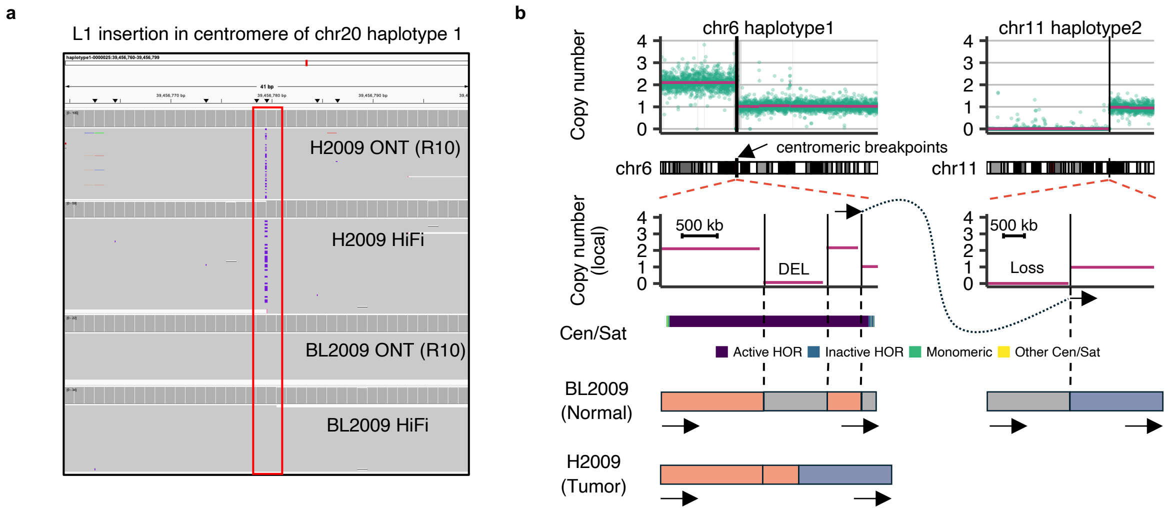


**Supplementary Figure 15. Somatic SVs within centromeric regions of the H2009 cell line.** (a) IGV shows read alignments of a somatic L1 insertion within the centromere of chromosome 20 haplotype 1. Tracks from top to bottom show H2009 ONT (R10), H2009 HiFi, BL2009 ONT (R10), and BL2009 HiFi alignments, with the inserted L1 region highlighted. (b) Centromere-involving structural rearrangement in H2009 between chromosome 6 haplotype 1 and chromosome 11 haplotype 2. A deletion occurred in the centromere region of chromosome 6 haplotype 1 and then a translocation occurred between the centromere region of chromosome 6 haplotype 1 and chromosome 11 haplotype 2 q arm with copy-number changes. The reconstructed structure in BL2009 (normal) versus H2009 (tumor), with arrows indicating chromosomal orientation is shown at the bottom.

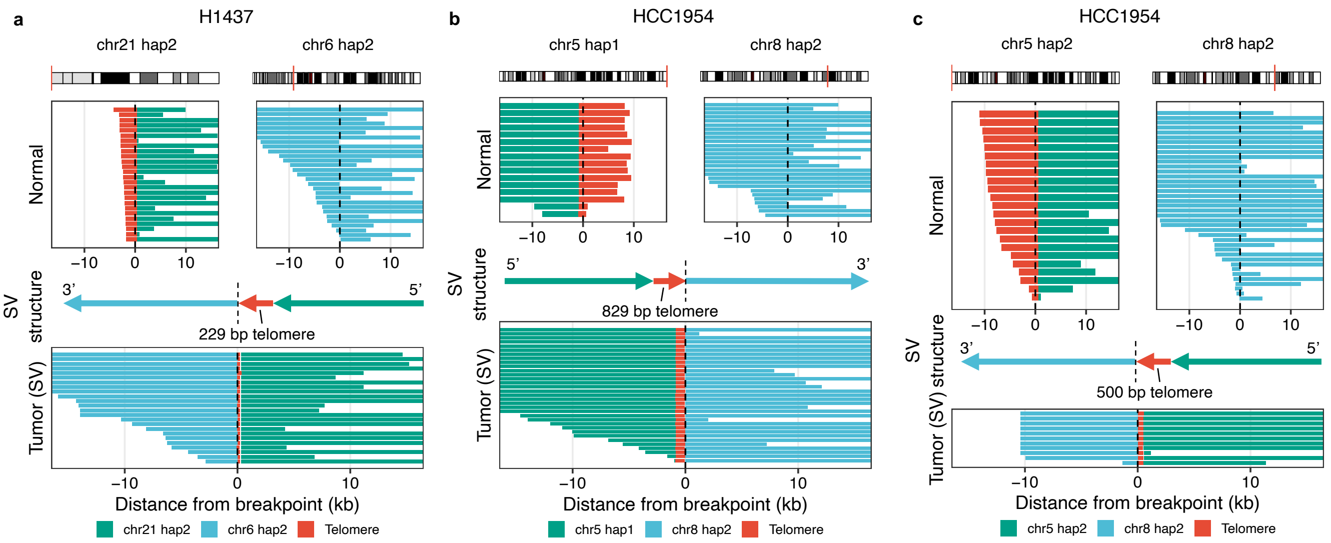


**Supplementary Figure 16. Examples of telomere-associated translocations.** (a) Telomere-associated translocation between chromosome 21 haplotype 2 and chromosome 6 haplotype 2 in the H1437 cell line. The chromosome ideograms are shown at the top, and read alignments around the breakpoint (0 kb, dashed line) are shown for the normal (BL1437) and tumor (H1437) samples. Between the Normal and Tumor panels, a schematic SV structure is shown with arrows indicating the 5'-to-3' direction on each segment and the length of the telomere block (229 bp) annotated. Reads are colored by origin: chr21 haplotype 2 (green), chr6 haplotype 2 (light blue), and telomere repeat sequence (red). (b) Telomere-associated translocation between chromosome 5 haplotype 1 and chromosome 8 haplotype 2 in the HCC1954 cell line (telomere block, 829 bp), shown as in a. (c) Telomere-associated translocation between chromosome 5 haplotype 2 and chromosome 8 haplotype 2 in the HCC1954 cell line (telomere block, 500 bp), shown as in a.

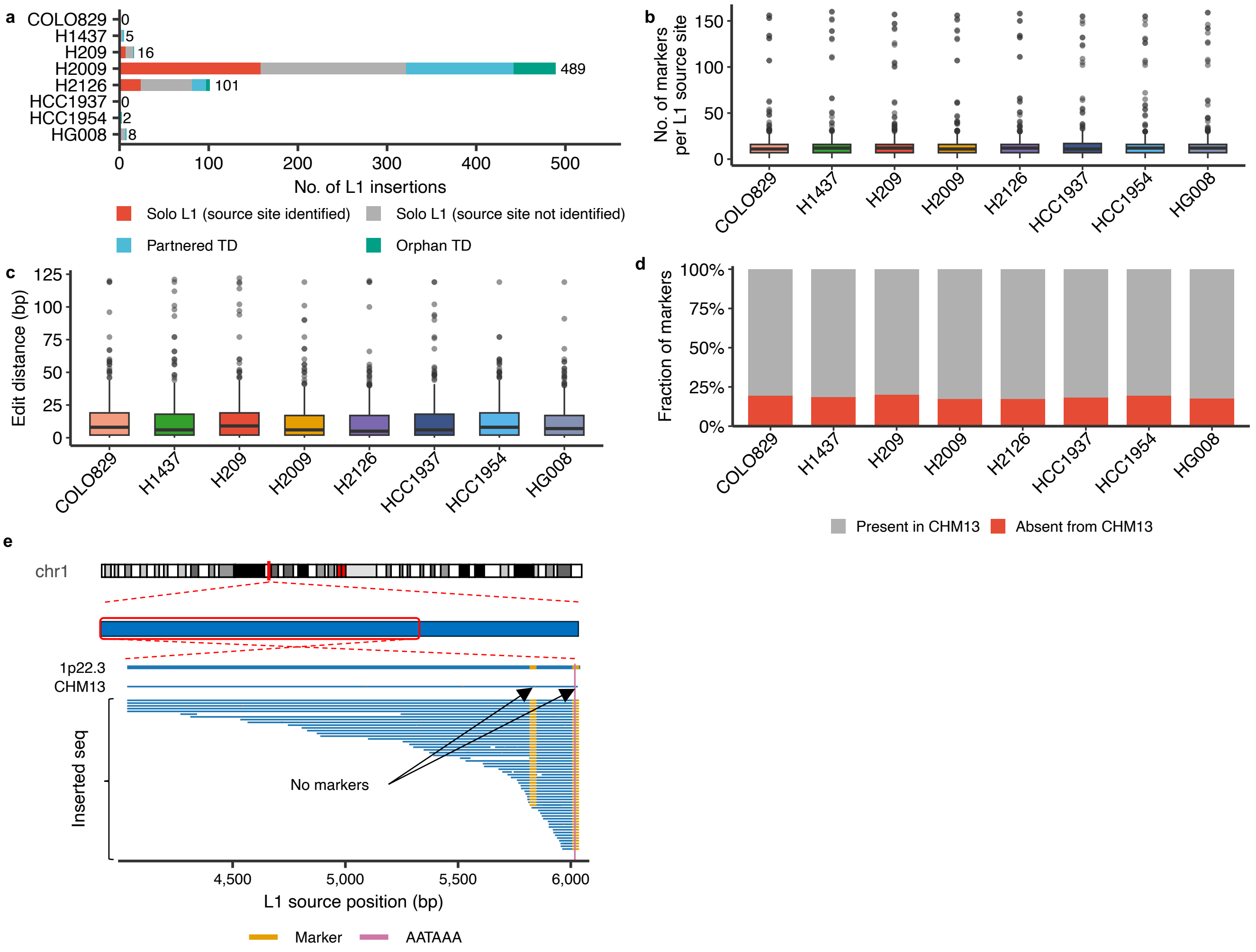


**Supplementary Figure 17. Characterization of somatic L1 insertions and the marker database used for L1 source-site identification across cancer cell lines.** (a) Number of somatic L1 insertions detected per cancer cell line (COLO829, H1437, H209, H2009, H2126, HCC1937, HCC1954, and HG008), classified into solo L1 insertions with identified source site (red), solo L1 insertions with unidentified source site (gray), partnered TD (light blue), and orphan TD (green). The H2009 cell line harbored the largest number of somatic L1 insertions. (b) Distribution of the number of markers per L1 source site, shown as a boxplot for each cell line. (c) Distribution of the edit distance between each full-length L1HS element in the personalized reference and the corresponding L1HS element in T2T-CHM13, shown as a boxplot for each cell line. (d) Fraction of markers in each cell line classified as present (gray) or absent (red) in T2T-CHM13. (e) Example of an L1 source site at 1p22.3 in the H2009 cell line. The chromosome 1 ideogram (top) and the alignment of inserted L1 sequences against the L1 source position, along with the marker positions and the internal polyadenylation signal (AATAAA) (bottom), are shown. The alignment of the corresponding L1 element in T2T-CHM13 (CHM13) is shown for comparison and illustrates that the markers supporting this source site are absent from T2T-CHM13, indicating that this source site would be challenging to identify using T2T-CHM13.

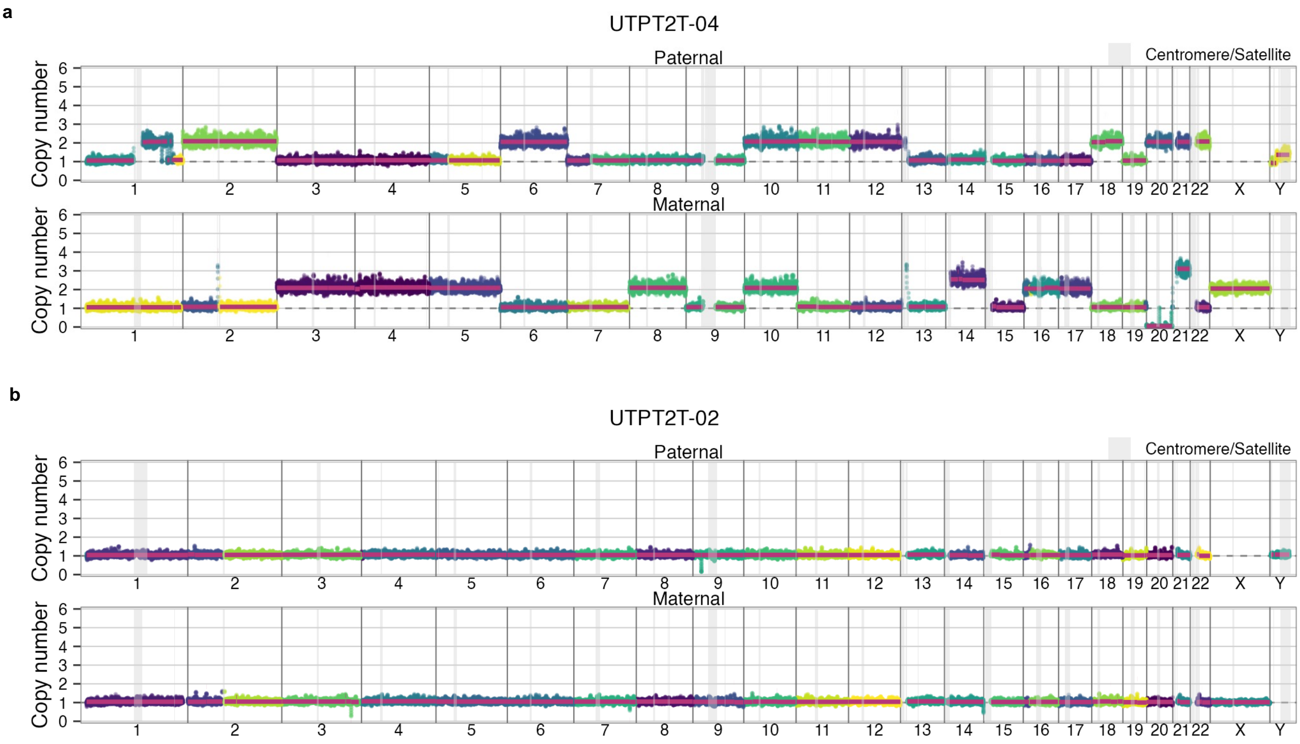


**Supplementary Figure 18. Haplotype-resolved copy-number profiles of pediatric cancer clinical samples.** (a, b) Haplotype-resolved copy-number profiles of the UTPT2T-04 (a) and UTPT2T-02 (b) clinical samples, shown separately for paternal (top) and maternal (bottom) haplotypes. Axes, coloring, and shading are as in Supplementary Fig. 10.

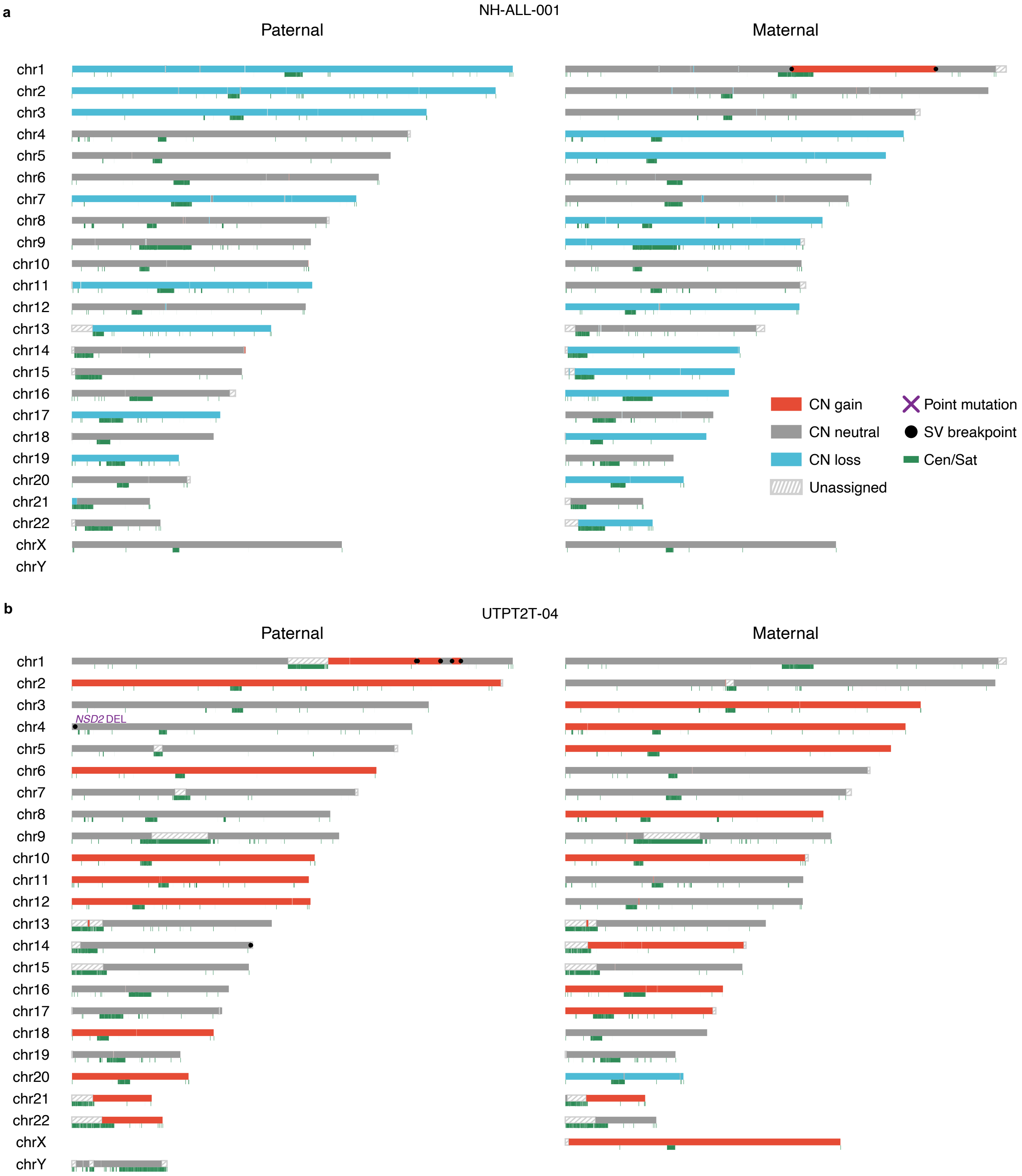


**Supplementary Figure 19. Haplotype-resolved genomic overview of pediatric cancer clinical samples.** (a–c) Haplotype-resolved genomic overviews of the NH-ALL-001 (a), UTPT2T-04 (b), and UTPT2T-02 (c) clinical samples, shown as in Supplementary Fig. 11.

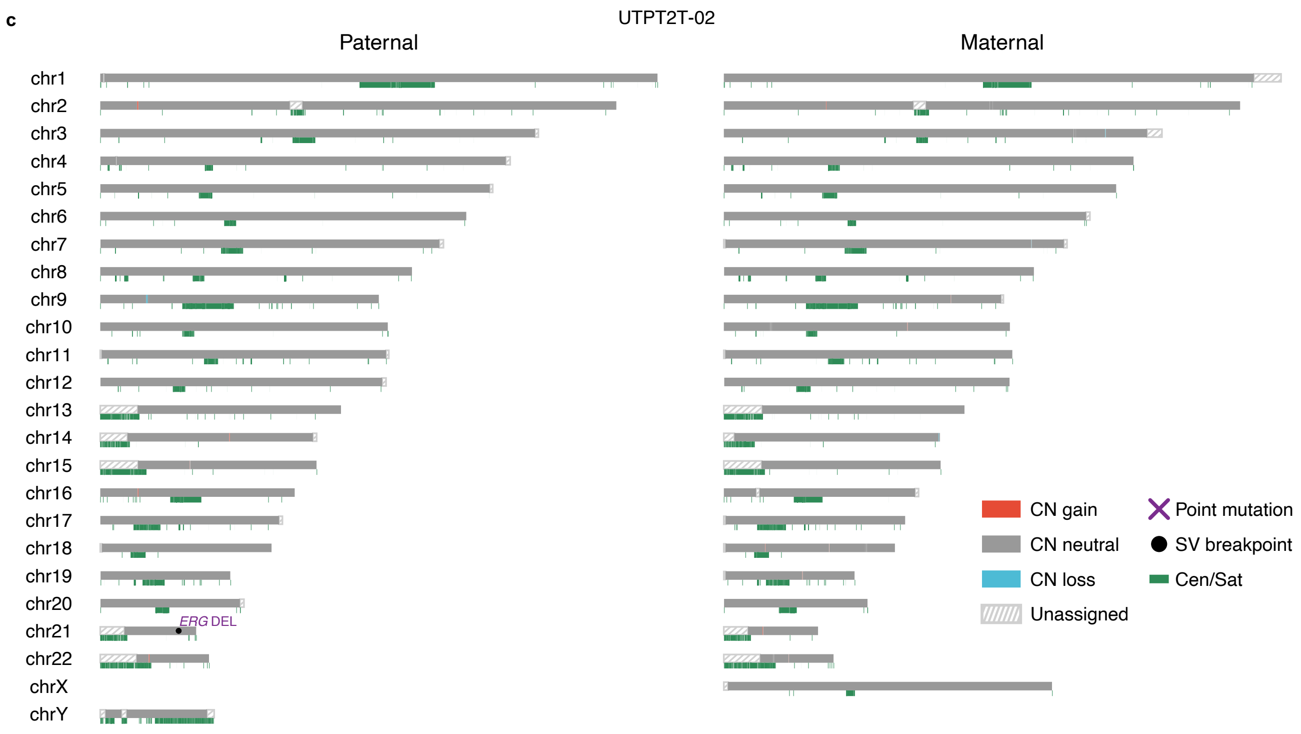


Supplementary Figure 19 (continued)

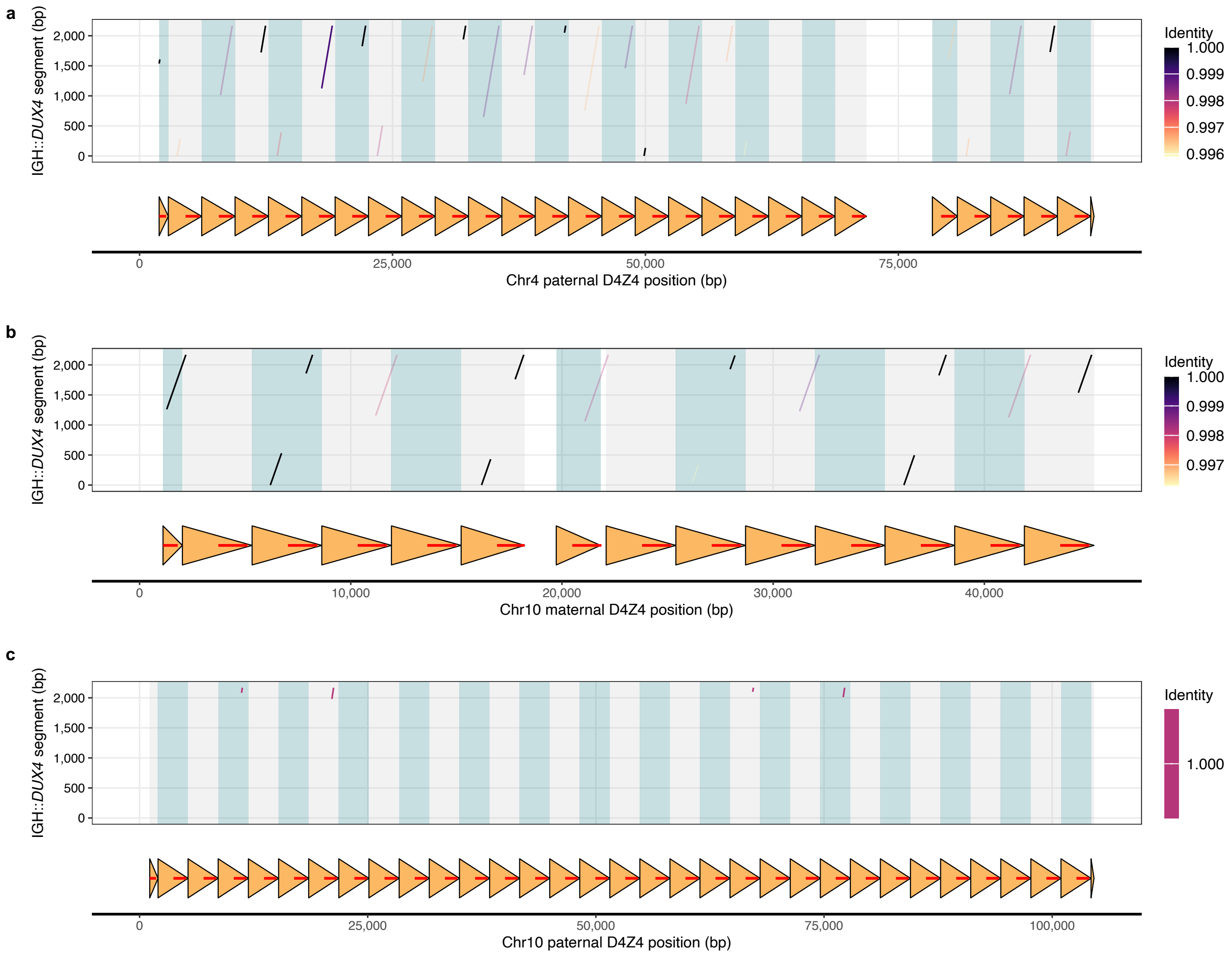


**Supplementary Figure 20. Alignment of the IGH::*DUX4* fusion segment to D4Z4 repeat arrays other than the maternal chromosome 4 array.** (a) Dot plot of the IGH::*DUX4* fusion segment (y-axis) aligned to the paternal chromosome 4 D4Z4 repeat array (x-axis). The color scale indicates percent identity. The schematic below shows D4Z4 repeat units (arrows) along the array. (b) Alignment of the IGH::*DUX4* fusion segment to the maternal chromosome 10 D4Z4 repeat array, shown as in a. (c) Alignment of the IGH::*DUX4* fusion segment to the paternal chromosome 10 D4Z4 repeat array, shown as in a. Only a small number of low-identity hits are observed, supporting that the fusion originated from a specific D4Z4 unit on the maternal chromosome 4 array.

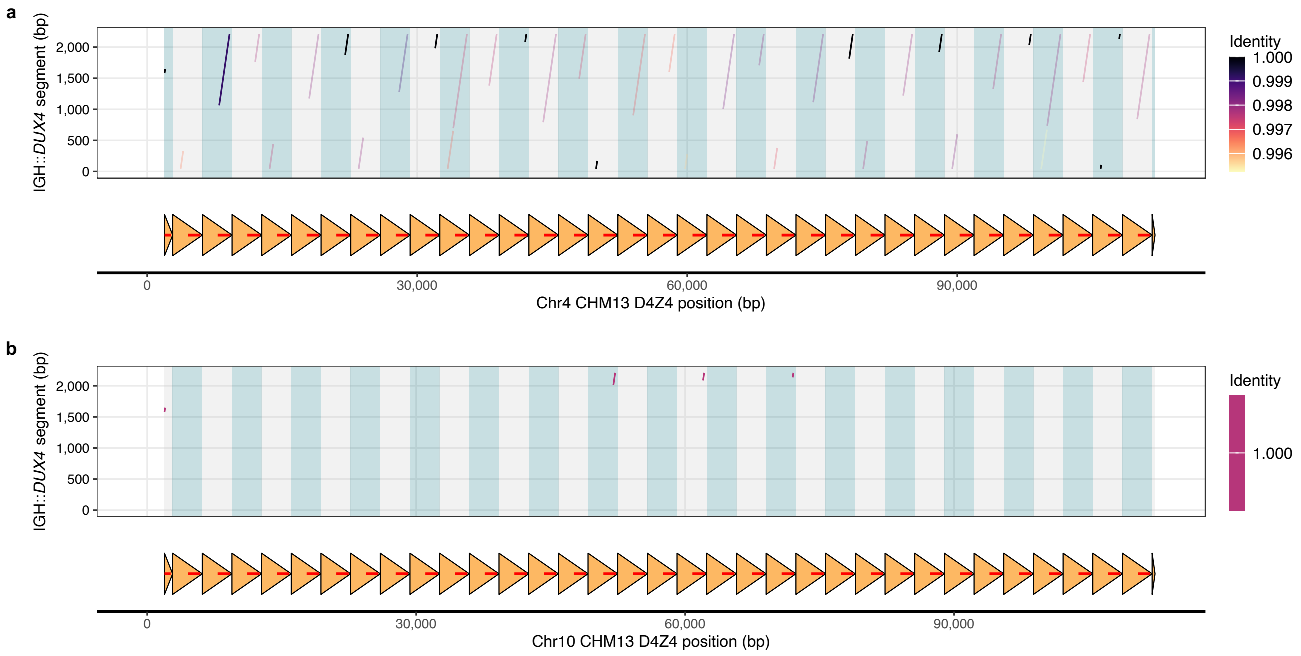


**Supplementary Figure 21. Alignment of the IGH::*DUX4* fusion segment to D4Z4 repeat arrays in T2T-CHM13.** (a) Dot plot of the IGH::*DUX4* fusion segment (y-axis) aligned to the T2T-CHM13 chromosome 4 D4Z4 repeat array (x-axis). The color scale indicates percent identity. The schematic below shows D4Z4 repeat units (arrows) along the array. (b) Alignment of the IGH::*DUX4* fusion segment to the T2T-CHM13 chromosome 10 D4Z4 repeat array, shown as in a.
